# Region-specific blood-brain barrier transporter changes leads to increased sensitivity to amisulpride in Alzheimer’s disease

**DOI:** 10.1101/582387

**Authors:** Gayathri Nair Sekhar, Alice L. Fleckney, Sevda Tomova Boyanova, Huzefa Rupawala, Rachel Lo, Hao Wang, Doaa B. Farag, Khondaker Miraz Rahman, Martin Broadstock, Suzanne Reeves, Sarah Ann Thomas

## Abstract

Research into amisulpride use in Alzheimer’s disease (AD) implicates blood-brain barrier (BBB) dysfunction in antipsychotic sensitivity. Solute carrier function in AD has not been widely studied. This study tests the hypothesis that organic cation transporters contribute to the BBB delivery of antipsychotics and is disrupted in AD.

I*n vitro* BBB studies indicated that [^3^H]amisulpride and [^3^H]haloperidol were transported by OCT1. Amisulpride also utilized PMAT. Molecular docking predicted that amisulpride and haloperidol are OCT1, PMAT and MATE1 substrates, and amisulpride is not a P-gp substrate. Amisulpride brain uptake increased in 3xTgAD compared to wildtype mice. PMAT and MATE1 expression was reduced in brain from AD patients compared to controls. The increased sensitivity of Alzheimer’s patients to amisulpride is related to previously unreported changes in OCT1, PMAT and MATE1 function/expression at the BBB. Dose adjustments may be required for drugs that are substrates of these transporters when prescribing for AD patients.

## 1. Background

Antipsychotic drugs are associated with significant harm in older people, particularly those with dementia who are more susceptible to antipsychotic drug related morbidity (parkinsonism, postural hypotension, stroke) and mortality than other diagnostic groups (Ballard and Howard 2006)(Schneider *et al.* 2006). This has led to restrictions on the NHS use of this class of drugs in the pharmacological management of psychosis and agitation in dementia. Emerging evidence from research into amisulpride use in older people with Alzheimer’s disease (AD) psychosis suggests that blood-brain barrier (BBB) dysfunction may be an important contributor to this heightened sensitivity (Reeves *et al.* 2017)(Caravaggio Fernando; Graff-Guerrero 2017).

Amisulpride is a benzamide derivative, second generation antipsychotic drug, used to treat schizophrenia (Mauri *et al.* 2014) and a drug for which the optimal dose (400-800mg/day), blood concentration (100-319ng/ml) and striatal dopamine D2/3 receptor occupancy range to avoid non-response and parkinsonism (40-70%) are well established in young adults with schizophrenia (Hiemke *et al.* 2011)(Sparshatt *et al.* 2009)(Lako *et al.* 2013). Despite being highly selective for dopamine D2/3 receptors *(in vitro* (K_i_=2.8nM) and D3 (K_i_=3.2nM)) amisulpride has a low propensity to induce parkinsonism, due to its poor BBB penetration and mesolimbic selectivity (Schoemaker *et al.* 1997). In an open treatment study which used amisulpride use in older people with AD psychosis and very late-onset (>60 years) schizophrenia-like psychosis (VLOSLP), treatment response and parkinsonism occurred at very low doses (25-75mg/day AD, 50-100mg/day VLOSLP), and at correspondingly low blood drug concentrations (40-100ng/ml AD, 40-169ng/ml VLOSLP) due to higher than anticipated striatal dopamine D2/3 receptor occupancies (caudate occupancy, steady state treatment, 50mg/day amisulpride; 41-83% AD, 41-59% VLOSLP) (Clark-Papasavas *et al.* 2014)(Reeves *et al.* 2016)(Reeves *et al.* 2017)(Clark-Papasavas *et al.* 2014)(Reeves *et al.* 2018). These findings strongly implicate age and AD-specific changes in central pharmacokinetics in antipsychotic drug sensitivity, particularly at the BBB, which controls drug entry through the expression of transporters (Saidijam *et al.* 2017). Furthermore, they suggest that amisulpride (Dos Santos Pereira *et al.* 2014)(Natesan *et al.* 2008) is a sufficiently sensitive tool with which to probe BBB functionality.

The majority of research into BBB transporters has been directed towards the ABC superfamily, which are ATP-dependent efflux transporters such as P-glycoprotein (P-gp) (Kania *et al.* 2011), whose action is compromised in age (A *et al.* 2012), and more markedly so in AD(Vogelgesang *et al.* 2002) (Deo *et al.* 2014)(Wijesuriya *et al.* 2010)(Park *et al.* 2014). It has been suggested that amisulpride is a P-gp substrate(Härtter *et al.* 2003) (Schmitt *et al.* 2012), but this has not been tested, and the importance of P-gp relative to other transporters, especially members of the SLC superfamily, remains unclear. Amisulpride is predominately positively charged (98.9%) at physiological pH (pKa 9.37), and is likely a substrate for the organic cation transporters (OCT) and organic cation transporters novel (OCTN); as observed using the immortalized human cerebral microvessel endothelial cell line (hCMEC/D3) (Dos Santos Pereira *et al.* 2014). However, it is also possible that other SLC transporters of organic cations, such as plasma membrane monoamine transporter (PMAT) and multi-drug and toxic compound extrusion proteins (MATEs), are involved. Haloperidol is a first generation anti-psychotic, one of the 20 drugs on the WHO list of essential medications and an OCT1 substrate and inhibitor(Ahlin *et al.* 2008)(Bourdet 2005).

This study tested the hypothesis that there was an interaction between amisulpride and influx and/or efflux transporters which was relevant from a pharmacodynamic perspective by:

1. identifying the transporter involved in the CNS distribution of amisulpride and haloperidol, by examining their kinetic characteristics and inhibitor sensitivity at the human and mouse BBB *in vitro*.
2. confirming the molecular level interactions of amisulpride and haloperidol with the selected BBB transporters using an *in silico* computational approach.
3. establishing whether amisulpride access to the CNS is increased in transgenic AD which harbour the human APP-Swedish mutation (KM670/671NL), tau mutation (P301L), and presenilin-1 mutation (M146V) compared to wildtype mice.
4. investigating transporter expression in human (and mouse) brain endothelium from age-matched post-mortem AD and healthy aged controls.
5. Examining the type of medications prescribed to patients with AD and age matched controls.

Overall BBB dysfunction in the AD process and its potential impact on drug delivery in particular on antipsychotic medication will be explored (Figure 1). The results can also be used to inform further studies. Abstracts of this work have been presented (Sekhar, G., Reeves, S. and Thomas 2015)(Boyanova, S., Wang, H., Reeves, S. and Thomas 2018).

**Figure 1:**
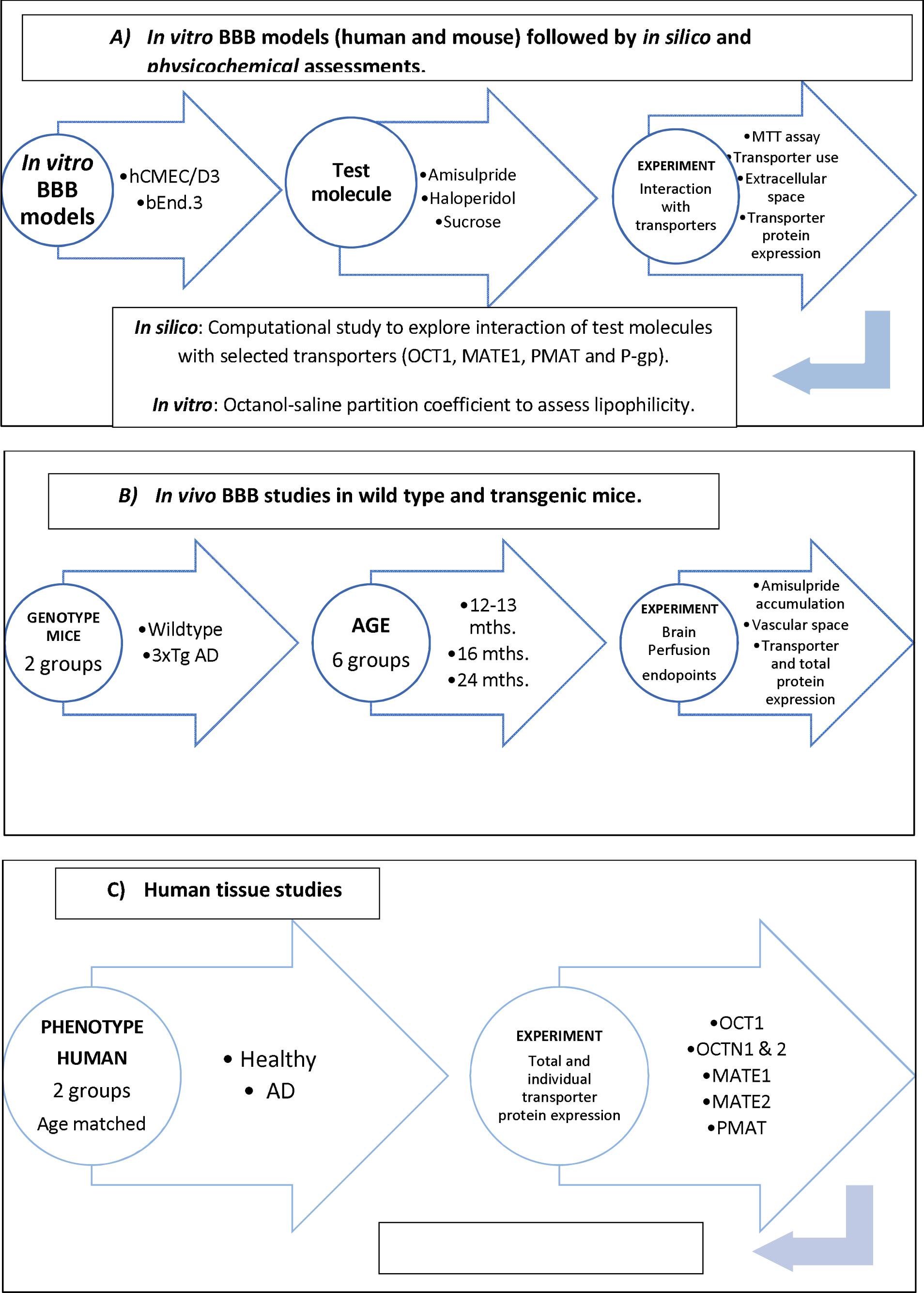
Flow charts to provide an overview of the experimental design for the *in silico, in vitro* and *in vivo* approaches. Experiments from the three approaches were performed in parallel.

## 2. Results

### 2.1.1 Amisulpride accumulation and saturable transport

[^3^H]Amisulpride was able to accumulate in both hCMEC/D3 and bEnd3 cell lines to a greater extent than the baseline marker, [^14^C]sucrose (Figures 2 and S1). Incubation of hCMEC/D3 and bEnd3 cell lines with 20 µM amisulpride significantly decreased the accumulation of [^3^H]amisulpride (6.5nM) at 5 minutes – with a significant decrease of 37% in hCMEC/D3 cells and a decrease of 50% in bEnd.3 cells observed after 2 hours (Figure 2). No significant differences were observed for [^14^C]sucrose between the treatments, except in the bEnd3 cells at 120 minutes where the presence of 20μM amisulpride decreased the accumulation of [^14^C]sucrose (Figure S1). However, all [^14^C]sucrose values were within the expected range for these *in vitro* models. Studies also revealed that lower concentrations of unlabelled amisulpride (0.1 μM) did not affect accumulation of [^3^H]amisulpride in hCMEC/D3 (n=5 passages) or bEnd.3 (n=3 passages) cells at all time points (data not shown).

**Figure 2:**
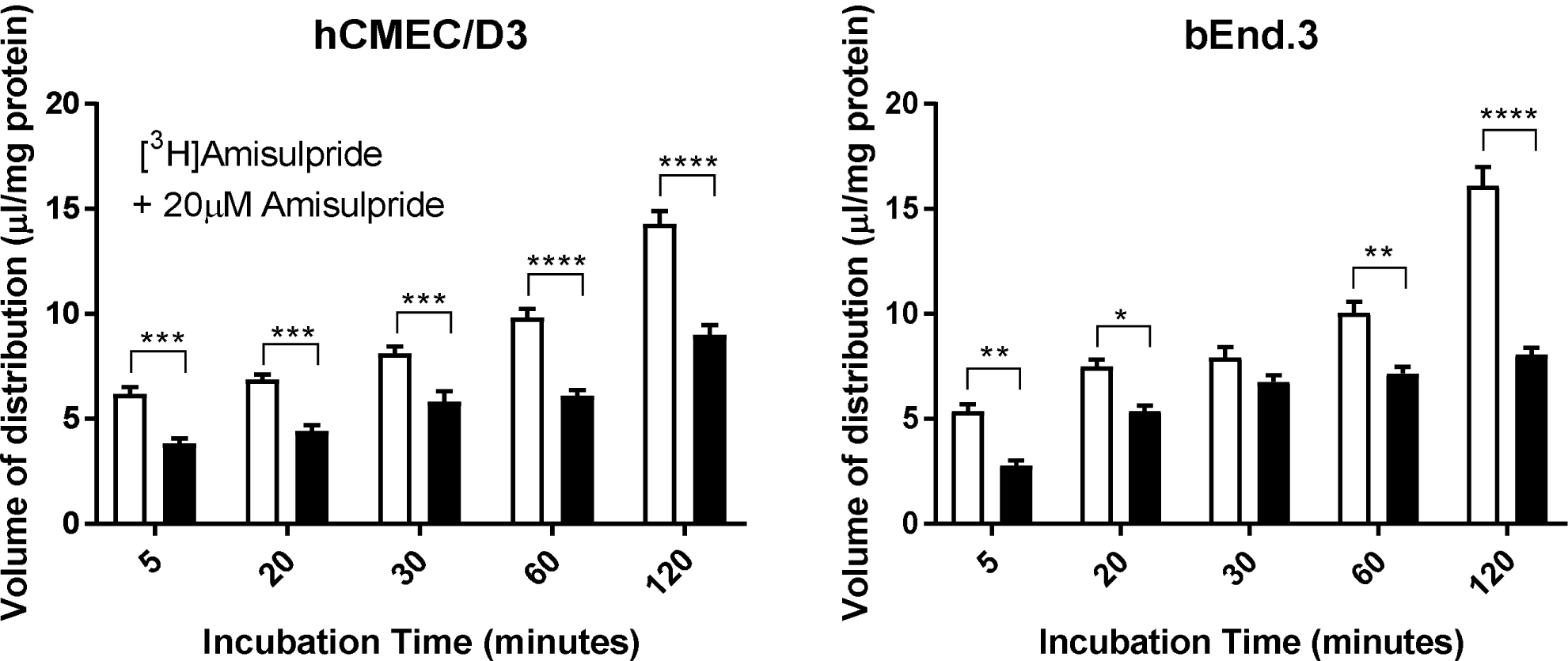
The effect of self-inhibition (20 μM) on the accumulation of [^3^H]amisulpride (6.5nM) was determined in hCMEC/D3 (A) and bEnd.3 (B) cell lines. Significant differences compared to control were observed -*p≤0.05, **p≤0.01, ***p≤0.001, ****p≤ 0.0001. All data have been corrected for [^14^C]sucrose and are expressed as mean ± S.E.M, n = 3 to 7 plates with 6 replicates (wells) per timepoint per plate (5 time-points).

### 2.1.2 ABC transporter involvement

ATP depletion did not affect the accumulation of [^3^H]amisulpride or [^14^C]sucrose in either cell line (Figure S2). The P-gp substrate, dexamethasone, BCRP substrate, ko143, and inhibitor, pheophorbide A, and MRP family inhibitor, MK571, did not affect the accumulation of [^3^H]amisulpride or [^14^C]sucrose in either cell line (Figure S3).

### 2.1.3 OCT, OCTN, PMAT and MATE involvement

Involvement of OCTs in [^3^H]amisulpride uptake was investigated by incubating hCMEC/D3 and bEnd.3 cells with the, OCT1 and 2 substrate, amantadine, the OCT1 and 3 substrate, prazosin, and the OCT3 substrate, corticosterone (Figure S4). [^3^H]amisulpride accumulation did not change in the presence of amantadine in hCMEC/D3 cells, but significantly increased by 84% in bEnd.3 cells. In the presence of prazosin, there was a significantly reduced accumulation of [^3^H]amisulpride in hCMEC/D3 cells, but not in bEnd.3 cells. Corticosterone did not affect the accumulation of [^3^H]amisulpride in either cell line. No differences were found for [^14^C]sucrose between the treatments (Figure S4).

[^3^H]amisulpride accumulation in hCMEC/D3 and b.End3 was unaffected by the presence of ergothioneine (OCTN1) and *L*-carnitine (OCTN2) respectively (Figures S5 and S6). [^14^C]Sucrose V_d_ was not significantly different between the treatments, except significant differences were observed between [^14^C]sucrose and *L*-carnitine at 2 hours suggestive that this time point for [^3^H]amisulpride should be ignored.

Incubation with the PMAT inhibitor, lopinavir, resulted in a significant increase in the V_d_ of [^3^H]amisulpride by 63.9% at 20 minutes, 83.2% at 30 minutes, 85.1% at 60 minutes and by 68.6% at 120 minutes (Figure 3). The V_d_ of [^14^C]sucrose was not significantly different between control and test groups (Figure 3). No affect was observed with this inhibitor in the b.End3 cells (Figure S7).

**Figure 3:**
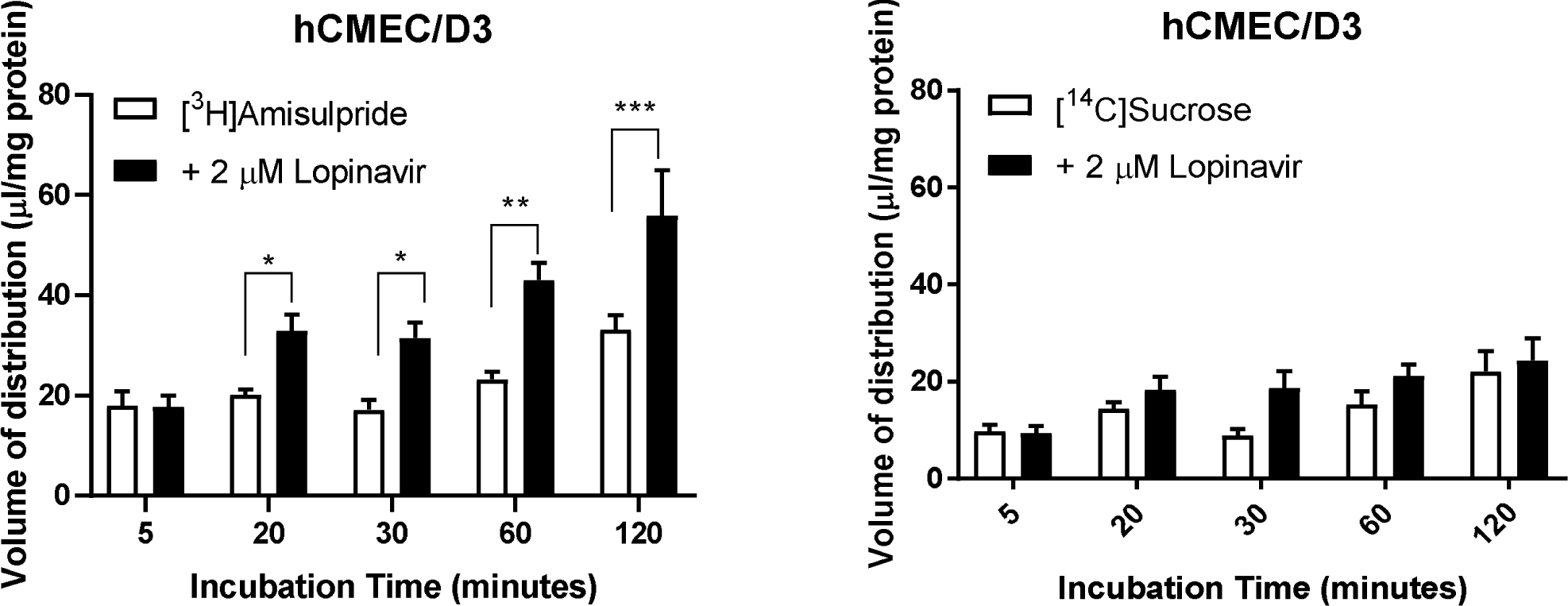
The effect of PMAT inhibition on the accumulation of [^3^H]amisulpride (3.7-7.7nM) was determined in hCMEC/D3 cell lines. Significant increases were observed compared to control ****p≤0.0001, ***p≤ 0.001, **p=0.01 and *p=0.05. [^3^H]amisulpride data has been corrected for [^14^C]sucrose and are expressed as mean ± S.E.M, n =5 passages (p30, 2 x p31, p32 and p34 for PMAT) with 6 replicates (wells) per timepoint per plate (5 time-points).

Incubation with the MATE1 inhibitor, famotidine (1 μM) resulted in no significant effect on the V_d_ of [^3^H]amisulpride in hCMEC/D3 cells (Figure 4). The V_d_ of [^14^C]sucrose was not significantly different between control and test groups with (1 μM) famotidine during the standard 2 hour incubation period. An assessment of the effect of 2 μM famotidine on [^14^C]sucrose alone revealed a loss of hCMEC/D3 integrity at 2 hours. Further assessment of famotidine (2 μM) did not affect either [^3^H]amisulpride or [^14^C]sucrose accumulation in hCMEC/D3 cells over a one hour period (Figure 4). No affect was observed with famotidine (1 μM) in b.End3 with either [^3^H]amisulpride or [^14^C]sucrose (Figure S7).

**Figure 4:**
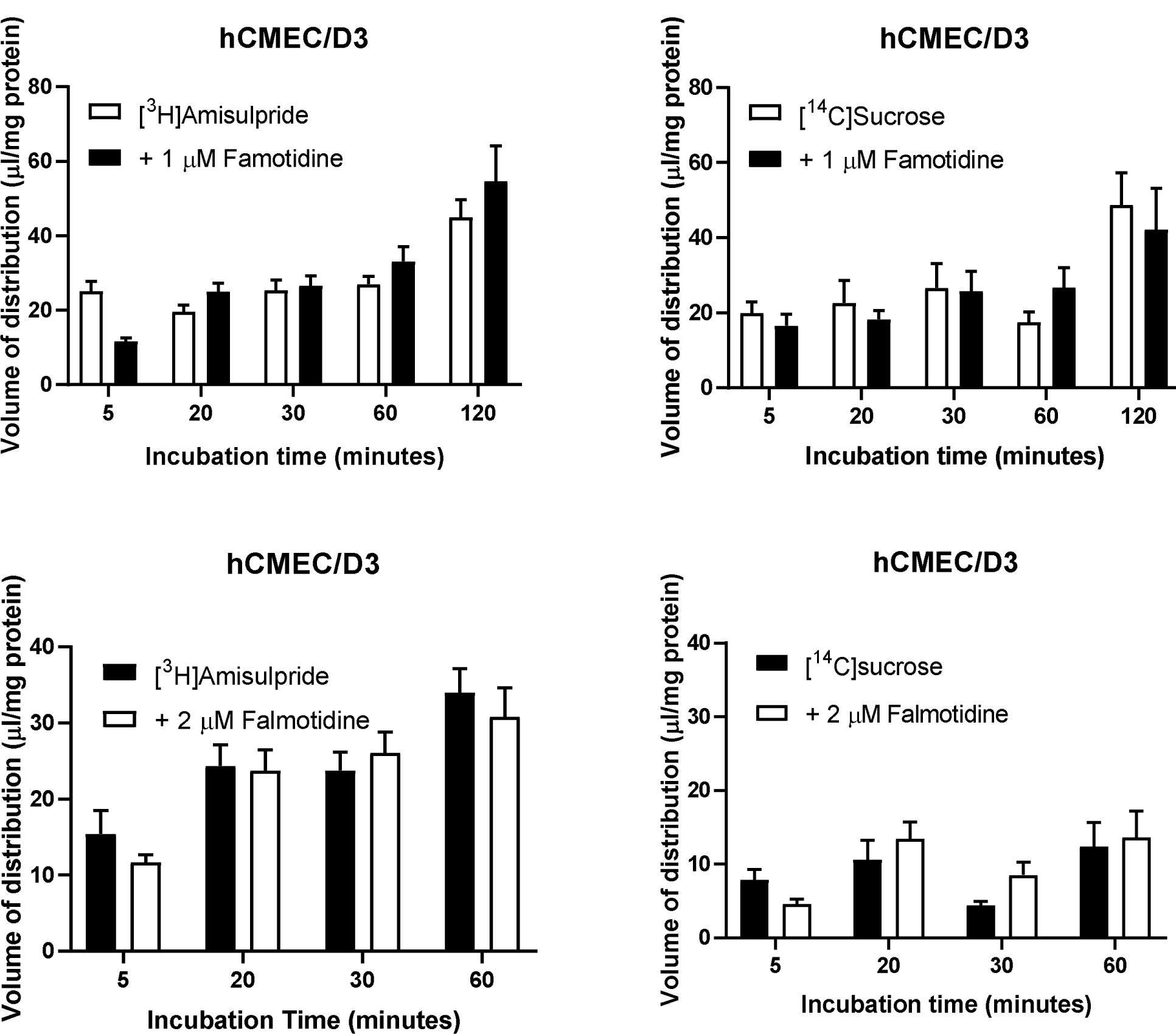
The effect of MATE1 inhibition on the accumulation of [^3^H]amisulpride (3.7-7.7nM) was determined in hCMEC/D3 cell lines. Significant increases were observed compared to control ****p≤0.0001, ***p≤ 0.001, **p=0.01 and *p=0.05. [^3^H]amisulpride data has been corrected for [^14^C]sucrose and are expressed as mean ± S.E.M, n = 4 passages (p30 x 2, p31 and p34 for MATE1 inhibitor falmotidine at 1 μM and n =3 passages for MATE1 inhibitor falmotidine at 2 μM) with 6 replicates (wells) per timepoint per plate (5 time-points).

Incubation with the MATE2 inhibitor, nifekalant, resulted in no significant affect on the V_d_ of [^3^H]amisulpride in hCMEC/D3 cells (Figure S8). The V_d_ of [^14^C]sucrose was not significantly different between control and test groups up to 120 minutes with nifekalant suggesting loss of membrane integrity at this time point. No affect was observed with this inhibitor in b.End3 with either [^3^H]amisulpride or [^14^C]sucrose (Figure S7).

### 2.1.4 Interaction with the positively charged anti-psychotic drug, haloperidol

The effect of the cationic drug, haloperidol (40μM), on radiolabelled amisulpride accumulation was also investigated. Incubation of unlabelled haloperidol with [^3^H]amisulpride did not yield any significant effects in either cell line (Figure S9). No significant differences were found for [^14^C]sucrose between the treatments (Figure S9).

### 2.1.5 Characteristics of haloperidol accumulation in hCMEC/D3 and b.End3 cell lines

hCMEC/D3 and b.End3 cells were incubated with unlabelled haloperidol (40μM) along with [^3^H]haloperidol (10nM). Incubation with unlabeled haloperidol significantly decreased the accumulation of radiolabelled haloperidol by approximately 93% in hCMEC/D3 cell line and by 94% in bEnd.3 cell line at all times (***p<0.001) (Figure S10). No significant differences were found for [^14^C]sucrose between the treatments (Figure S10).

ATP was depleted from both cell lines to determine the role of ABC transporters in the efflux of haloperidol. ATP depletion did not affect the accumulation of haloperidol in either cell line (Figure S11). No significant differences were observed for [^14^C]sucrose between the treatments (Figure S11). The hypothesis that haloperidol uptake is by OCT transporters was investigated by incubating the cells with OCT1 and 2 substrate amantadine (500μM) and OCT1 and 3 substrate prazosin (100μM). [^3^H]haloperidol accumulation significantly decreased in the presence of amantadine in both cell lines compared to control - by 89% in hCMEC/D3 cells and by 82% in bEnd.3 cells and in the presence of prazosin - by 85% in hCMEC/D3 cells and by 82% in bEnd.3 cells (***p<0.001) (Figure S12). No significant differences were found for [^14^C]sucrose between the treatments (Figure S12).

The effects of other cationic drugs – unlabeled pentamidine (100μM), unlabeled efornithine (250 μM) and unlabelled amisulpride (20 μM) on radiolabelled haloperidol accumulation in hCMEC/D3 was investigated. Unlabelled pentamidine significantly reduced the accumulation of radiolabeled haloperidol in the cell lines after 2 hours - by 31% in hCMEC/D3 cells (***p<0.001, **p<0.01, and *p<0.05). Unlabelled eflornithine significantly decreased the accumulation of radiolabelled haloperidol by 11% in hCMEC/D3 cells (***p<0.001). Unlabelled amisulpride (20 μM) significantly decreased the accumulation of radiolabelled haloperidol in hCMEC/D3 cells - by 27% after 2 hours (***p<0.001) (Figure S13). No significant differences were found for [^14^C]sucrose between the treatments (Figure S13).

### 2.1.5 Cytotoxicity

No cytotoxic effects of amisulpride (0.1-20μM), 1μM famotidine, 2μM famotidine, 2μM lopinavir, 20μM ergothioeine, 5μM L-carnitine, 3μM nifekalant hydrochloride and 1.5μM [^14^C]sucrose (Figures S14A and S14B) and eflornithine (250-500μM) were detected using the MTT assay (data not shown). A marker molecule ([^14^C]sucrose) of extracellular/vascular space was included in all [^3^H]amisulpride accumulation experiments and ensured that any measured effect on [^3^H]amisulpride values could be interpreted correctly and was not simply due to loss of membrane integrity caused by the cytotoxic nature of the drugs/inhibitors utilized.

### 2.2 Lipophilicity

The octanol-saline partition coefficient for [^3^H]amisulpride was determined to be 0.0422±0.0045 and for [^3^H]haloperidol was determined to be 0.6678±0.1278.

### 2.3.1 Molecular docking studies with the SLC transporters-OCT1, MATE1 and PMAT

Amisulpride showed molecular interactions inside the binding site of OCT1 in the form of hydrogen bonds with amino acids Gln283, Asn288, Glu380 and Asn415 as well as hydrophobic interactions with amino acid residues Phe26 and Pro385 (Figure 5A), while haloperidol showed hydrogen bonds with amino acids Gln 283, Gly384, Trp388 and hydrophobic interactions with Gly384 and Pro385 while prazosin interacted with Asn288, Thr321, Ser324, Glu380, Asn411 through hydrogen bonds and hydrophobic interactions with Leu325, Val328, Gly384 and Pro385 (Figure S15A, ESI). The best pose of amisulpride interacted with the binding pocket of OCT1 with a free energy of binding of −14.28 kcal/mole while the free energy of binding for haloperidol and prazosin were −29.97 kcal/mole and −27.57 kcal/mole.

**Figure 5:**
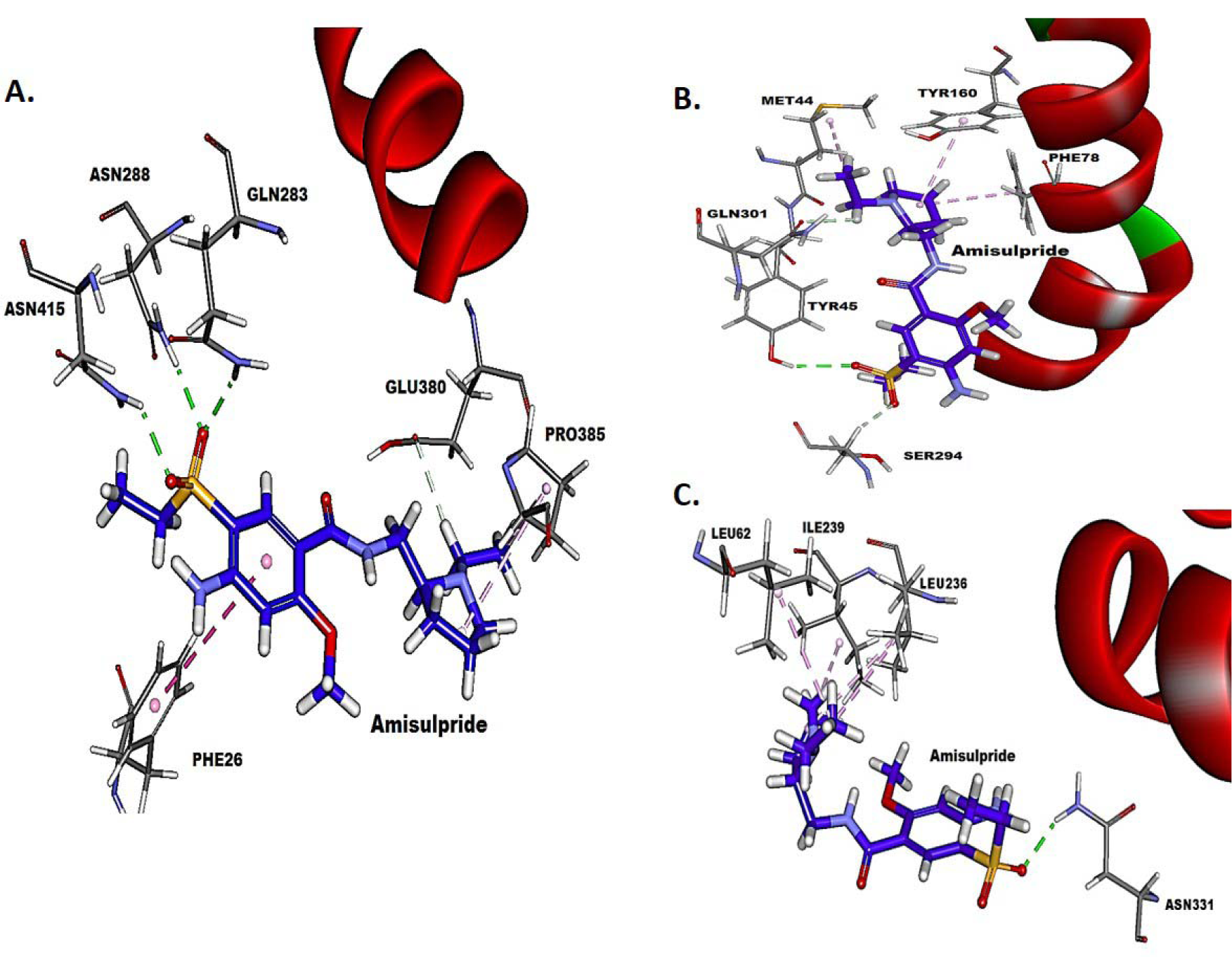
Molecular-level interactions of amisulpride within the binding site of OCT1 (A), MATE1 (B) and PMAT (C). Amisulpride is represented in stick-representation and amino acid residues in line-representations. Hydrogen bonds are represented in green dotted lines, and hydrophobic interactions are represented in pink dotted lines.

Amisulpride showed a similar level of interaction with MATE1 transporter with a free energy of binding of −14.32 kcal/mole. It fit snugly within the binding pocket (Figure 5B) and formed hydrogen bonds with amino acids Tyr45, Ser294 and Gln301 as well as hydrophobic interactions with amino acid residues Met44, Phe78 and Tyr160. The hydrophobic interactions appeared to play an important role in its interaction with MATE1 compared to its interaction with OCT1. The interaction of amisulpride was relatively weaker with PMAT compared to both OCT1 and MATE1 with free energy of binding −11.4 kcal/mole. It formed a single hydrogen bond with Asn331 and interacted with hydrophobic interactions with amino acid residues Leu62, Leu236 and Ile239 through hydrophobic interactions (Figure 5C).

A similar molecular docking study suggested haloperidol is a better substrate of both MATE1 and PMAT compared to amisulpride as it showed binding affinity of binding −22.27 kcal/mole and −21.72 kcal/mole for MATE1 and PMAT, respectively, which are notably higher than amisulpride (Figures 5 and S16, ESI). It showed good interaction with MATE1 with hydrogen bonds with Tyr45 and Ser74 and hydrophobic interaction with Phe82 and Ala67 (Figure S16A, ESI). However, the interaction of haloperidol with PMAT was limited to single hydrogen bond with Asp34 and hydrophobic interaction with Leu62 (Figure S16B, ESI).

### 2.3.2 Molecular docking studies with the ABC transporter-P-gp

The molecular docking study revealed that amisulpride was not a substrate for P-gp with a free energy binding of −1.81 kcal/mol and the molecule was not able to interact favourably with the binding pocket of P-gp. P-gp substrates, dexamethasone and colchicine, showed notably superior interaction with P-gp with free energy of binding values of −31.83kcal/mol and −16.07 kcal/mol. Both dexamethasone and colchicine interacted with the binding pocket employing hydrogen bonds and hydrophobic interactions (Figure S17 and Table S1).

### 2.4 CNS amisulpride delivery *in vivo*

The R_TISSUE_ values for [^3^H]amisulpride did not differ from the R_TISSUE_ values for [^14^C]sucrose in wildtype mice in all the age groups tested (Table S2). There was also no effect of ageing on the brain distribution of [^3^H]amisulpride or [^14^C]sucrose. No differences were observed in the [^14^C]sucrose R_TISSUE_ values in the frontal and occipital cortex between the wildtype and transgenic mice (Figure 6). However, in transgenic mice the R_TISSUE_ value for [^3^H]amisulpride was significantly higher than [^14^C]sucrose in the frontal cortex, but not the occipital cortex. Importantly, in transgenic mice the sucrose-corrected R_TISSUE_ value for [^3^H]amisulpride was significantly higher than that of wildtype mice in the frontal cortex, but not the occipital cortex (Figure 6).

**Figure 6:**
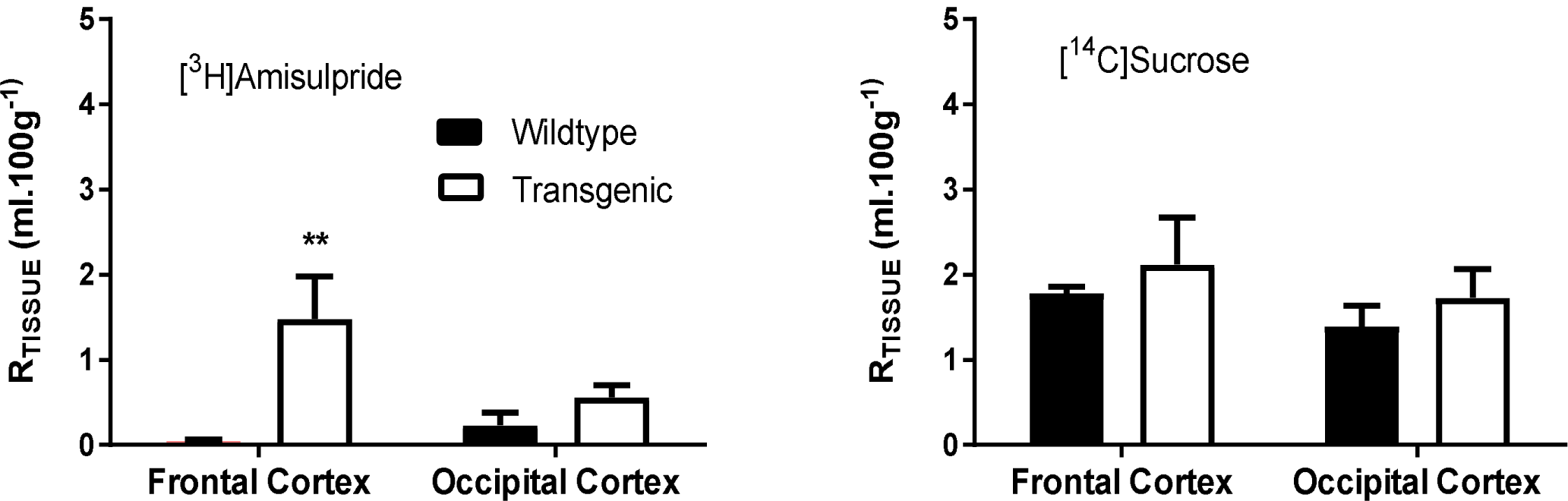
The uptake of [^3^H]amisulpride was determined in wildtype and 3xtransgenic AD mice. Significant differences were observed for [^3^H]amisulpride between wildtype (n=5 frontal cortex and n=6 occipital cortex) and transgenic mice (n=4 each region) - **p<0.005. [^3^H]Amisulpride data have been corrected for [^14^C]sucrose. [^14^C]Sucrose uptake is shown. No differences in paracellular permeability and membrane integrity were observed. All data are expressed as mean ± S.E.M, n = 4-6 mice, 2 years old. Perfusion time was 10 minutes. 6 C57BL6/129 mice (3 males and 3 females: weight 37.0±1.8g) and 4 transgenic (2 males, 2 females: weight 29.1±1.0g) were used. Also see Table S2.

### 2.5 Endothelial Transporter expression

#### 2.5.1 Cell lines

OCTN1, OCTN2, MATE1 and MATE2 expression was confirmed in hCMEC/D3 (passages 28 and 33) and bEnd.3 (passages 18, 19 and 23) cells (Figures S18 and S19A). PMAT was expressed in hCMEC/D3 cells (passages 28, 31 and 32) and bEnd.3 (passages 17, 18, 20 and 24) (Figure S19B and C).

#### 2.5.2 Wildtype and Transgenic AD mice

The total protein concentration measured in wildtype mice (188.2±12.8 μg/100μl) was not significantly different to that measured in 3xTg AD mice (195.6±13.3 μg/100μl) brains, but this may be attributed to the fact that it was not possible to assess regional differences in these small samples. Although slight variability was observed, there was no significant differences in individual transporter (P-gp, OCT1, OCT2, OCT3, OCTN1, OCTN2, MATE1 and MATE2) expression between the wildtype and 3xTgAD mice (Figures S20 and S21). Data not shown for P-gp, OCT1, OCT2 and OCT3. The expression of all these BBB transporters suggests that a capillary enriched sample had been assessed.

#### 2.5.3 Human brain

Human brain capillaries were isolated from the frontal cortex (for comparison with *in situ* perfusion experiments), caudate nucleus and the putamen (forming the striatum where high D2 and D3 receptor occupancy is observed in AD patients with amisulpride usage) of healthy controls and age-matched AD affected individuals (Table S3 and S4). The total protein concentration in the capillaries was found to be significantly lower in the caudate nucleus (by 37.4%) and putamen (by 32.5%), but not the frontal cortex samples, from AD patients compared to healthy controls (Table S5).

Individual transporter expression in each brain region between AD and healthy cases is comparable as the same amount of protein has been loaded into each well. Note this amount was dependent on the antibody utilized so was variable. Transporter expression in the frontal cortex was less variable between healthy and AD cases than in the other regions studied with no significant differences in transporter expression being observed (Figures S22-S28; Figure 7). Expression of OCT1 did not change between control and AD patients in all the regions tested (Figures S22-S23). PMAT and MATE1 expression was significantly lower in AD patients in the caudate nucleus (56.2%; P<0.05) and putamen (74.8%; P<0.05) samples, respectively, compared to control (Figure 7; Student’s t-test). No other significant differences were observed, however, further cases are required to explore this more fully. Please note it is also likely that transporter expression in caudate nucleus and putamen AD samples is even lower than the heathy controls shown here (Figure 7; Figures S22-S28) as total protein expression is significantly reduced (Table S5). The expression of all these BBB transporters would also indicate that a capillary enriched sample had been assessed.

**Figure 7:**
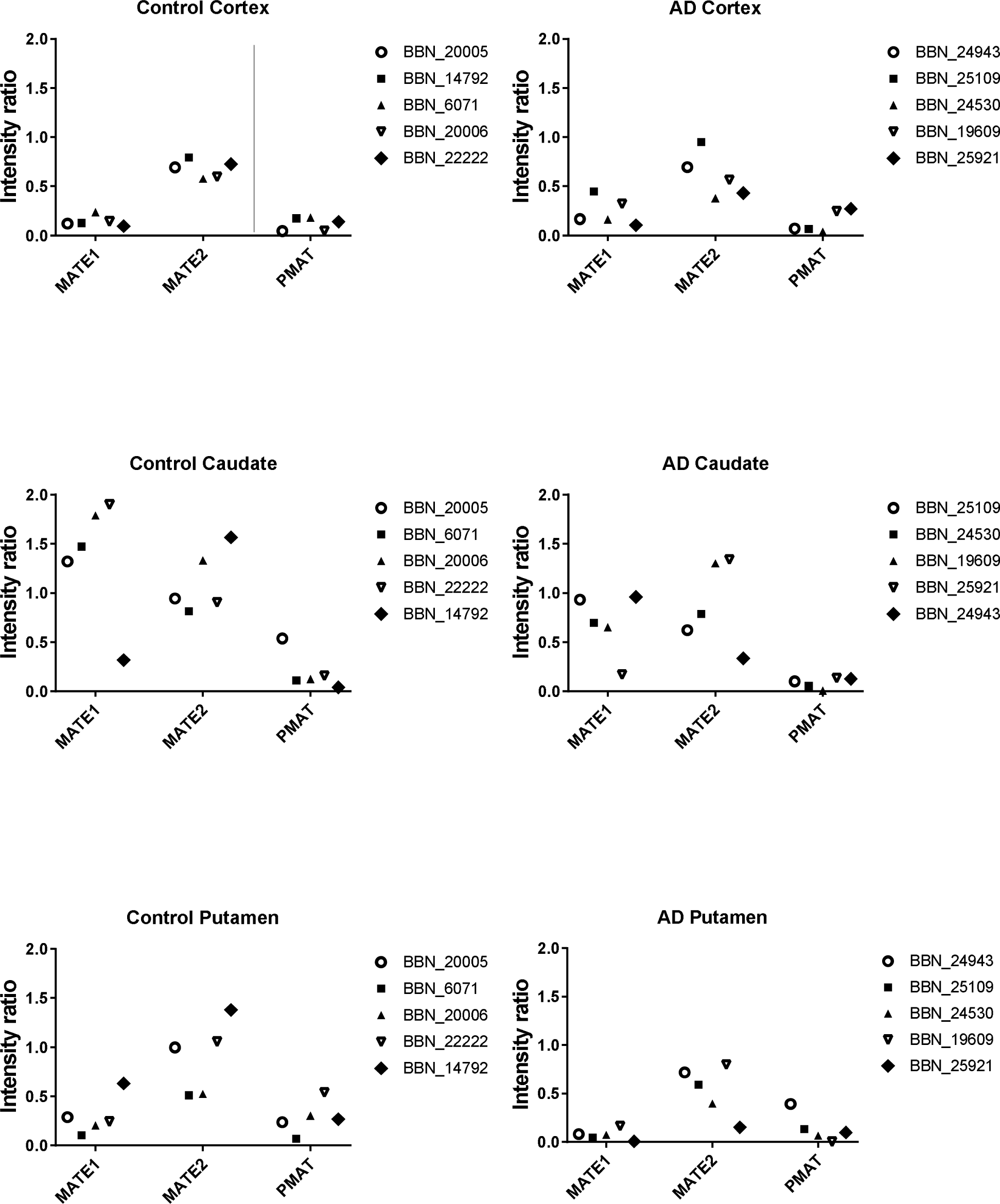
Individual values have been plotted for the transporter expression in the capillaries of frontal cortex, caudate nucleus and caudate putamen samples from healthy and AD affected individuals. The numbers in the key indicate the MRC ID designated to each sample. Details of the samples can be found in the supplementary information Tables S3 and S5.

### 2.6 Medication history of the cases

Table S4 shows the medication history of the cases that have been used in this study. Sedatives, antidepressants and antipsychotic drugs (including haloperidol) were identified and listed together. A separate column lists all other medications.

## 3. Discussion

This study aimed to answer an important clinical question using an integrative approach to investigate the interaction between two drugs and BBB transporters and their potential pharmacodynamic relevance in AD. The *in vitro* cell culture models and *in silico* computational model allowed us to (1) identify the transporters, (2) assess the potential mechanisms of amisulpride and haloperidol transport (3) perform studies on human and mouse brain endothelium and (4) minimize animal studies in line with the 3Rs (replacement, refinement and reduction principles). The *in situ* brain perfusion technique allowed us to (1) study the whole animal (2) and utilize a mouse model of AD. Importantly this is the only model to exhibit both amyloid-β_40_ and _42_ and tau pathology, mimicking human AD (Oddo *et al.* 2003)(Hirata-Fukae *et al.* 2008). Both plaque and tangle pathology are mainly restricted to the hippocampus, amygdala and cerebral cortex. Transporter expression in capillaries isolated from AD and age-matched human cases and mouse brain were also assessed. Medication history of the cases was evaluated.

Cell culture studies revealed a slow accumulation of [^3^H]amisulpride indicating a low BBB permeability. This is linked to its low lipophilicity, as measured by the octanol-saline partition coefficient, and its inability to interact strongly with neutral and negatively charged lipid model systems (Härtter *et al.* 2003)(Skrobecki *et al.* 2017). Please note the plasma half-life of a single oral dose of amisulpride (50 or 200mg) is ∽12 hours, which suggests that amisulpride will not have significantly degraded within the 2 hour incubation period in this study(Coukell *et al.* 1996). Note the accumulation buffer does not contain plasma enzymes. Unlabelled amisulpride (20µM) significantly reduced [^3^H]amisulpride accumulation in both cell lines, suggesting that there is a relatively low affinity influx transporter at the plasma membrane or an organelle membrane (e.g. lysozyme) (Schwake *et al.* 2013). OCT1, OCT2 and 3 have been shown to be expressed in hCMEC/D3 and b.End3 cells in an earlier study by our group(Sekhar *et al.* 2017) and OCTN1 and OCTN2 were shown to be expressed in both cell lines in this present study.

In line with the findings of (Dos Santos Pereira *et al.* 2014), our accumulation studies hinted that OCT1 may be involved, as prazosin (substrate for OCT1 and 3) reduced [^3^H]amisulpride uptake in hCMEC/D3 cells after 2 hours, whereas corticosterone (OCT3 substrate) had no impact on [^3^H]amisulpride accumulation in either cell line. In addition, OCTN1 and 2 inhibitors did not affect accumulation of [^3^H]amisulpride into hCMEC/D3 or bEnd.3 cells. However, in the presence of amantadine, a substrate for several transporters of organic cations (OCT1, OCT2 (Dickens *et al.* 2012)), MATE 1, MATE 2 (Nies *et al.* 2011)(Tsuda *et al.* 2009) and PMAT (Itagaki *et al.* 2012)) there was no effect on hCMEC/D3 cells, and an increase in [^3^H]amisulpride accumulation in bEnd.3 cells after 2 hours incubation, which suggests the involvement of an efflux transporter. This is consistent with previous observations (Härtter *et al.* 2003), which found higher amisulpride transport in the basolateral to apical direction (Pe 5.2±3.6 x 10^-6^cm/s) compared to the apical to basolateral direction (Pe<10^-7^cm/s) in porcine brain microvessel endothelium. The differences in the effect of inhibitors on the two cell lines may be explained by amisulpride being a substrate for multiple transporters and variations in the function/expression of OCT1, MATE and PMAT transporters possibly related to species differences (Shimomura *et al.* 2013)(Wu *et al.* 2015). The absence of any effect of prazosin on [^3^H]amisulpride uptake in bEnd.3 cells can be explained by prazosin-associated toxicity in this cell line, which causes protein values to decrease over the course of the experiment, resulting in no net effect on V_D_ values(Sekhar *et al.* 2017). It is unlikely that this efflux transporter is an ABC transporter, as neither ATP-depletion or substrates for P-gp, BCRP, or the MRP family had an effect on [^3^H]amisulpride accumulation in either cell line. Importantly our molecular docking studies also revealed that amisulpride was not a P-gp substrate unlike dexamethasone and colchicine. However, *in vitro* inhibitor studies with lopinavir, suggest that amisulpride could be effluxed by PMAT. This transporter is expressed on brain capillaries (Figures 7, S19B, S19C and S28) (Shimomura *et al.* 2013)(Hiasa *et al.* 2006)(Wu *et al.* 2015)(Kurosawa *et al.* 2018). PMAT mRNA and protein has also been identified on the luminal and abluminal membrane of human, mouse and rat brain endothelial cells(Wu *et al.* 2015).

It is important to highlight that the lack of inhibitor effect may not be conclusive proof of a lack of substrate interaction with the transporter. It may be that the transporter was not sufficiently expressed. For example in a recent study, MATE1 mRNA was below the limit of quantification in hCMEC/D3 cells (Kurosawa *et al.* 2018), but had been detected in an earlier study albeit at low levels(Shimomura *et al.* 2013). Other considerations are that the inhibitor may need to reach a therapeutic concentration within the cell to elicit a response as has been observed with MATE1 inhibitors (Tsuda *et al.* 2009), the substrate and the inhibitor may bind to different binding sites, the non-specificity of the inhibitor, and that amisulpride interacts with both influx (OCT) and efflux (PMAT) transporters(Wittwer *et al.* 2013)(Wu *et al.* 2015)(Bourdet 2005).

Haloperidol has been observed to have a high degree of dopamine receptor (D2) occupancy within the brain at very low doses suggesting that haloperidol is very efficient at crossing the BBB. This is in agreement with our *in vitro* BBB data. Please note that the half-life of haloperidol has been reported to range 14.5-36.7 hours (or up to 1.5 days) after a single oral dose so will not have been metabolized significantly over our 2 hour incubation period(de Leon *et al.* 2004). Haloperidol may cross by passive diffusion, as supported by the relatively high octanol-saline partition coefficient of haloperidol (0.6678 ± 0.1278) compared to amisulpride (0.0422 ± 0.0045), or may involve transporters. The use of transporters by haloperidol is confirmed by the self-inhibition studies in both hCMEC/D3 and b.End3 cell lines. As haloperidol exists predominately (94.8%) as a positively charged drug at physiological pH (pKa is 8.66) the transporter is likely to be OCT, which is expressed at the BBB. [^3^H]Haloperidol was incubated with the OCT substrates, amantadine (OCT1 and 2) and prazosin (OCT1 and 3), which confirmed the involvement of OCTs at the BBB in haloperidol transport. The transport of haloperidol from the BBB into the brain is also likely to be carried out by OCTs since they are expressed at the luminal and abluminal membrane of the BBB. We also investigated the involvement of ABC transporters in the transport of haloperidol. For this, ATP was depleted from the cells by incubating them with 10 mM 2-deoxy-d-glucose. No effects of ATP depletion were observed compared to control in either cell line suggesting that haloperidol is not a *substrate* for ABC transporters P-gp, BCRP, or the MRP family at the BBB as previously observed (Iwaki *et al.* 2006)(Schinkel *et al.* 1996).

Radiolabelled amisulpride (6.5nM) was also incubated with haloperidol (OCT1 substrate and P-gp *inhibitor*)(Ahlin *et al.* 2008)(Matsson *et al.* 2009). Radiolabeled amisulpride accumulation was not affected by haloperidol (40 μM) in either cell line. This may be the result of interactions with both OCT1 and P-gp, although our inhibitor and *in silico* studies do not suggest amisulpride is a substrate for P-gp. Conversely when radiolabeled haloperidol (10nM) was incubated with unlabeled amisulpride (20μM) there was a significant decrease in accumulation. Overall these results may reflect differences in the interaction of amisulpride, haloperidol and prasozin with specific OCT1 transporter binding sites.

Further insight into the interaction of amisulpride, haloperidol and prasozin with the transporters was obtained through the *in silico* computational studies, where amisulpride showed binding affinities towards the binding sites of the influx transporter OCT1 as well as the efflux transporters MATE1 and PMAT. The binding affinities of amisulpride towards the binding sites of these transporters showed comparable energies for OCT1 and MATE1 with molecular level interactions through hydrogen bonds and hydrophobic interactions with a number of amino acids within the binding pocket. The nature of the interaction of amisulpride with OCT1 is similar to that observed for haloperidol and prazosin which are known substrates for this transporter. The binding affinity and level of interaction were slightly weaker with PMAT compared to OCT1 and MATE1, but still considerable level of interactions were observed suggesting amisulpride is potentially a weak substrate of PMAT. Haloperidol showed comparable affinity for both MATE1 and PMAT suggesting it is a substrate for both transporters.

The *in silico* study supports the experimental observations and provides further evidence that amisulpride might be influxed through OCT1 and effluxed through PMAT and MATE1 but not P-gp. When amisulpride transport was investigated *in vivo*, a low BBB permeability to [^3^H]amisulpride was also observed in the wild-type mice and this did not change with age. Further studies in the 3xTG AD mice revealed an increased CNS uptake which was not accounted for by altered BBB integrity or changes in vascular space in the AD model mice, as there were no differences in [^14^C]sucrose uptake between the two groups; and neither was it explained by non-expression of the transporters studied (P-gp, OCT1, OCT2, OCT3, OCTN1, OCTN2, MATE1 and MATE2). However, in post-mortem human brains expression of the efflux transporter MATE1 was lower in AD patients compared to age-matched healthy controls in the putamen; and PMAT showed a similar trend in the caudate nucleus, but there was no change in expression levels of these transporters in the frontal cortex. Importantly it has been reported that BBB impairments stem from AD abnormality instead of from vascular comorbidities (van de Haar *et al.* 2016). Further evidence for regional tissue changes with AD came from our total protein measures (Table S5). This may be linked to changes in the expression of other transporters such as P-gp (Vogelgesang *et al.* 2002) and GLUT1 (Horwood and Davies 1994) as well as the SLC transporters measured in this present study. Interestingly regional differences are known to exist in the BBB transport and the intracellular distribution of antipsychotics (Loryan *et al.* 2016). MATE1 is an H^+^/organic cation antiporter, which is expressed at the luminal membrane of renal tubule cells where it takes up protons from the filtrate, in exchange for the efflux of organic cations (André *et al.* 2012)(Ito *et al.* 2012)(Hiasa *et al.* 2006). At this site MATE1 is thought to work in cooperation with P-gp (Hiasa et al., 2006). Co-localization of MATE1 with members of the ABC transporter family has previously been reported(Staud *et al.* 2013). This suggests that MATE1, like P-gp, is expressed on the luminal membrane of the BBB, although this remains to be confirmed. PMAT transport activity is pH-dependent and it may also use a proton gradient to drive substrate efflux(Wu *et al.* 2015). The reduced MATE1 and PMAT expression observed in AD may therefore underpin the heightened sensitivity to amisulpride observed in the clinical population especially as our *in silico* and *in vitro* studies suggested that amisulpride was a substrate for MATE 1 and PMAT. Several medications listed in Table S4 (e.g. citalopram, metoclopramide and loratadine) have previously been identified as OCT1 inhibitors (Ahlin *et al.* 2008)(Bourdet 2005). It is not yet known if they also interact with MATE1 and PMAT. One of the medications (rantidine) is an inhibitor of both OCT1 and MATE(Staud *et al.* 2013).

### Conclusion

This study included a detailed evaluation of transporter expression and usage at the BBB using *in silico* computational approaches, *in vitr*o models and an *in vivo* animal model of AD as well as patient material. The datasets have provided evidence of an interaction of amisulpride and haloperidol with both influx (OCT1) and efflux (MATE1 and PMAT) transporters, which may be expressed at the luminal or abluminal membranes of the BBB and/or at an intracellular membrane. Furthermore, the study is of key importance as the results suggest that the heightened sensitivity to amisulpride observed in older people with AD is due to previously unreported changes in SLC transporter expression, which increase amisulpride entry into, or possibly reduce clearance from the brain. This study is also the first step in the process of characterising age and AD-specific changes in SLC transporters of organic cations. Overall our study has implications beyond antipsychotic prescribing, as it suggests that dose adjustments may be required for other drugs (e.g. haloperidol) which are substrates for SLC transporters in particular MATE1 and PMAT.

## 4. Materials and Methods

### 4.1 Materials

[O-methyl-^3^H]amisulpride (MW374.8; specific activity 77Ci/mmol; 97% radiochemical purity) was custom tritiated (TRQ41291 Quotient, UK). [^3^H(G)]haloperidol (MW375.9; specific activity, 20 Ci/mmol; 99% radiochemical purity: catalogue number ART1729) was purchased from American Radiolabelled Chemicals Inc, St. Louis, Missouri, USA. [^14^C(U)]sucrose (MW359.48; specific activity 536mCi/mmol; 99% radiochemical purity: cat# MC266) was purchased from Moravek Biochemicals, USA. Amisulpride (MW369.5, >98% purity) was purchased from Cayman Chemicals, UK (cat#71675-85-9). Haloperidol (MW375.9; >98% purity) was purchased from Sigma-Aldrich, Dorset, UK (cat#H1512). Anti-SLC22A1 antibody (Cat#ab55916; RRID:AB_882579), Anti-SLC22A2 antibody (Cat#ab170871: RRID:AB_2751021, Anti-SLC22A3 antibody (Cat#ab183071; RRID:AB_2751016), Anti-SLC22A4 antibody (Cat#ab200641; RRID:AB_2751017), Anti-SLC22A5 antibody (Cat#ab180757; RRID:AB_2751018), Anti-SLC47A1 antibody (Cat#ab104016: RRID:AB_10711136), Anti-SLC47A2 (Cat#ab174344: RRID:AB_2751019), SLC29A4 antibody (Cat#ab56554: RRID:AB_2190909), Goat Anti-Rabbit IgG H&L (cat#ab6721: RRID:AB_955447) and Rabbit anti-mouse HRP (Cat#ab6728: RRID:AB_955440) were purchased from Abcam, UK. Anti-SLC47A1 (Cat#ab174344: RRID:AB_2751019) was purchased from Alomone Laboratories, Israel. Anti-SLC29A4 (Cat#: bs-4176R: RRID:AB_11108960) was purchased from Bioss antibodies, USA. Goat anti-rabbit (IgG)-HRP was purchased from (Cell Signalling. Cat#7074S:AB_2099233). Table S6 details the dilutions used.

### 4.2 *In vitro* Model of the BBB

#### 4.2.1 Cell culture

The hCMEC/D3 (human) and bEnd.3 (mouse) are well-established cell lines that model the BBB *in vivo* (Weksler *et al.* 2013)(Brown *et al.* 2007)(Sekhar *et al.* 2017)(Park *et al.* 2014). They are not listed on the misidentified cell line register (version 9 released 14^th^ October 2018). Both cell lines require different cell mediums to grow to confluence and their BBB phenotype has been confirmed using Western blots, confocal and transmission electron microscopy (Sekhar *et al.* 2017)(section 4.10). Functional expression of several transporters has also been demonstrated by our group(Sekhar *et al.* 2017)(Watson *et al.* 2016)(Watson *et al.* 2012).

a. hCMEC/D3 cells (passages 27-35) were provided under a MTA and maintained in Clonetics® endothelial cell growth medium-2 MV Bullet Kit (Cat# CC-3162 Lonza, UK) containing the endothelial basal medium, the SingleQuotsTM growth factor kit, foetal bovine serum (FBS), penicillin-streptomycin and HEPES (Sigma-Aldrich, UK) (Poller *et al.* 2008)(Watson *et al.* 2012)(Sekhar *et al.* 2017).
b. bEnd.3 cells (passages 17-25), isolated from the SV129 strain of mice and transformed with the Polyoma virus middle T-antigen, were purchased from ATCC® (CRL-2299: RRID:CVCL0170). They were grown in T-75 flasks (Fisher Scientific, catalogue number 15350591) using high glucose Dulbeccos Modified Eagles Medium (Sigma-Aldrich, UK, catalogue number D6429) supplemented with 10% FBS (vol/vol) and 1% penicillin-streptomycin (vol/vol: Fisher Scientific cat#10003927).

Both lines were maintained at 37°C/ 5% CO_2_ in an incubator with saturated humidity. Medium was changed every 2-3 days. Cells were split when they reached 80-90% confluency and were seeded onto 96-well plates (ThermoScientific, UK) at 20,000 cells/cm^2^for b.End3 cells and 25,000cells/cm^2^ for hCMEC/D3 cells. The hCMEC/D3 96-well plates had been pre-coated with 0.1mg/ml rat tail collagen type1 (Gibco cat#A1048301).

Both cell lines were confluent in 4-5 days and they were left for a further 4 days to allow for them to further di□erentiate before experimentation. The medium was changed every 2-3 days.

#### 4.2.2 Drug accumulation assay

Accumulation assays were performed on confluent cell monolayers grown in the centre 60 wells of 96-well plates. Each passage was regarded as one ‘n’. No sample calculation was performed. The accumulation bu□er (pH7.4) composition was 135mM NaCl, 10mM HEPES, 5.4mM KCl, 1.5mM CaCl_2_, 1.2mM MgCl_2_, and 1.1mM D-glucose and water. It also contained 3.7-7.7nM [Omethyl-^3^H]amisulpride and 9.4µM [^14^C(U)]sucrose or 10nM [^3^H]haloperidol and 3.8μM [^14^C]sucrose. [^14^C]sucrose is a similar size to the test molecules and was used as an inert marker of extracellular space and membrane integrity. After the exposure period, bu□er was aspirated and the wells washed with ice-cold PBS^+^ (Sigma-Aldrich, UK) to remove drug that was not taken up by cells and to stop further transport. 1% Triton X-100 (Sigma-Aldrich) was added and the plate was incubated for an hour at 37°C to lyse the cells and to release accumulated [^3^H]drug. 100µl from each of the wells was transferred to a vial and scintillation fluid (4ml) added (Optiphase Hisafe 2, PerkinElmer, UK). Radioactivity was measured using a Packard Tri-Carb 2900TR liquid scintillation counter (PerkinElmer, UK) and corrected for background. The remaining 100µl in each well were used to perform a bicinchoninic acid (BCA) protein assay. A range of 2-30 μl.mg^-1^ of protein was acceptable. All data for [^3^H]amisulpride or [^3^H]haloperidol were expressed as a volume of distribution (V_d_) after correction for [^14^C]sucrose. The V_d_ was calculated from the sum of accumulated radioactivity (a sum of e□ux and influx of the molecule) (disintegrations per minute (dpm)/mg protein) over the ratio of dpm/μl of accumulation buffer. Outliers were identified by examining the [^14^C]sucrose values. [^14^C]sucrose values are presented in the figures and tables.

#### 4.2.3 Transporter inhibition assay

Transporter interaction was examined by incubating [^3^H]amisulpride and [^14^C]sucrose with potential inhibitors. These included unlabelled amisulpride and inhibitors/substrates of OCT, OCTN, MATE, PMAT, P-gp, BCRP, and MRP at established concentrations (Table S7). The MATE1 inhibitor, famotidine, was also utilized at a higher concentration of 2μM. ATP depletion (Promega Enliten assay) was carried out by pre-incubation with the glycolysis inhibitor, 10mM 2-deoxy-d-glucose, for an hour before incubating the cells with [^3^H]amisulpride and [^14^C]sucrose. This assay had previously been shown to inhibit drug efflux by our group (Watson *et al.* 2012)(Sekhar *et al.* 2017).

#### 4.2.4 Cytotoxicity assay

Cytotoxicity of unlabelled amisulpride (0.1-20μM) and eflornithine (250-500μM) on both cell lines was assessed using 3-(4,5-dimethylthiazol-2-yl)-2,5-diphenyltetrazolium bromide (MTT) assay (Sekhar *et al.* 2017). This assay was also utilized on hCMEC/D3 cells (passage 32 and/ or 33) for 2 hours to assess toxicity of 1μM famotidine, 2μM famotidine, 2μM lopinavir, 20μM ergothioeine, 5μM L-carnitine, 3μM nifekalant hydrochloride and 1.5μM [^14^C]sucrose. We have already published MTT assay results for eflornithine (250μM) in hCMEC/D3 cells and haloperidol (40μM), dexamethasone (200μM), pentamidine (10μM), ko143 (1μM), MK571 (10μM), amantadine (500μM), corticosterone (50μM), pheophorbide A (1μM) and prazosin (100μM) in both cell lines (Watson *et al.* 2012)(Sekhar *et al.* 2017). No significant effect was observed except with prazosin on bEnd.3 cells.

### 4.3 Lipophilicity

An octanol-saline partition coefficient for [^3^H]amisulpride and [^3^H]haloperidol was determined (Anthonypillai *et al.* 2006).

### 4.4 *In silico* computational study

Using an *in silico* molecular docking study, we tested the molecular level interactions of amisulpride, haloperidol and prazosin (OCT1 and OCT3 substrate) with the influx and efflux BBB transporters OCT1, PMAT and MATE1. Due to the unavailability of the crystal structures of these transporters, the molecular models of OCT1, PMAT and MATE1 were developed using homology modelling with Swiss-model webserver using PDB codes 4PYP, 5Y50 and 4ZOW, respectively, as templates (ESI). A further molecular docking study was carried out to explore the interaction of amisulpride, dexamethasone (P-gp substrate) and colchicine (P-gp substrate) in the binding site of the multidrug transporter ABCB1 (P-glycoprotein), and the pbd code 6FN1 (Alam *et al.* 2018) was used as the template for this study. Molecular docking was performed using Dock Ligands (CDOCKER) protocol from Discovery studio version 4.0. CDOCKER is an implementation of a CHARMm based docking tool where each orientation is subjected to simulated annealing molecular dynamics. The binding sites were chosen after comparing the docking results of the ligands in all possible binding cavities within the transporters.

### 4.5 Animal model of AD

All experiments were carried out in accordance with the Animal Scientific Procedures Act (1986) and Amendment Regulations 2012 and with consideration to the ARRIVE guidelines. The study was approved by the King’s College London Animal Welfare and Ethical Review Body and performed under a home office project license (70/7755). All mice used in this study were housed at King’s College London. C57BL6/129 mice (wild-type) and the triple transgenic AD model (3xTgAD) of C57 mice which harbour the human APP-Swedish mutation (KM670/671NL), tau mutation (P301L), and presenilin-1 mutation (M146V) were utilized at 12-13 (mid-age), 16 (old) or 24 (elderly) months old. The 3xTgAD mice is an established AD model since it represents the temporal and spatial progression of AD and mirrors the neuropathological development seen in AD(Oddo *et al.* 2003). These mice were bred and genotyped at King’s College London and were a gift. They had originally been supplied by the Jackson Laboratory (RRID:IMSR_JAX:008880). No sample size calculation was performed. Previous experience suggested that even with the limited number of animals available there would be sufficient power (95%) to detect significant differences (p<0.05)(Sanderson *et al.* 2008). Welfare was assessed by animal technologists on a daily basis. Animals were identified by earmarks and housed together by age and genotype in guideline compliant cages. All animals were maintained under standard temperature/lighting conditions and given food and water *ad libitum*. The experiment on each animal had to be performed within set time frames to allow the three age groups to be achieved. Experiments were performed between 9am and 5pm. The experimenter was not blinded.

The median life span of 3xTgAD mice has been previously reported to be 673 days (22 months) which is shorter than the 907 day (30 months) lifespan of C57BL/6J (Rae and Brown 2015). Adult male BALB/c mice (inbred strain) were purchased from Harlan UK Limited (Oxon, UK). All mice were anaesthetised (2mg/kg i.p. medetomidine hydrochloride and 150mg/kg i.p. ketamine) and heparinized (100U i.p.) in a procedure room separate to the main laboratory where the brain perfusion method was performed. Advice was sought from the named veterinary surgeon regarding the type and route of anaesthetic.

#### 4.5.1 *In situ* brain perfusion

To assess differences in [^3^H]amisulpride and [^14^C]sucrose transport into the brain in ageing and in AD, wildtype (C57BL6/129) and transgenic AD (3xTg) mice were used. Perfusion (10 minutes; 5ml/min) with a warmed (37°C) and oxygenated (95%O_2_; 5% CO_2_) artificial plasma was via a cannula in the left ventricle of the heart as previously described (Sanderson *et al.* 2007). The artificial plasma consisted of a modified Krebs-Henseleit mammalian Ringer solution with the following constituents: 117mM NaCl, 4.7mM KCl, 2.5mM CaCl_2_, 1.2mM MgSO_4_, 24.8mM NaHCO_3_, 1.2mM KH_2_PO_4_, 10mM glucose, and 1g/liter bovine serum albumin. With the start of perfusion the right atrium of the heart was sectioned to prevent the recirculation of the artificial plasma. At the end of perfusion the animal was decapitated and the brain removed. To determine the brain concentration of [^3^H]amisulpride, frontal and occipital cortex samples were taken using a Leica S4E stereozoom microscope and weighed. Brain samples were then solubilized with 0.5ml of Solvable (PerkinElmer, USA). Liquid scintillation fluid (3.5ml; Lumasafe; PerkinElmer) was then added. Radioactivity concentrations in the samples were counted using the Tri-Carb2900TR scintillation counter.

#### 4.5.2 Expression of results

Brain samples (dpm per gram) were expressed as a percentage of the drug concentration that was present in the artificial plasma (dpm per millilitre) and termed R_TISSUE_ (millilitres per 100g). All R_TISSUE_ values for [^3^H]amisulpride were corrected for vascular/extracellular space by subtracting the [^14^C]sucrose R_TISSUE_ value. Importantly examination of the [^14^C]sucrose values (i.e. vascular space) and comparison to previously published values determines if the result is an outlier. This is acceptable for the WT animals, however, it is noted that [^14^C]sucrose (vascular space) may be affected in AD. It is noted that no outliers were detected in this study.

### 4.6 hCMEC/D3 and b.End3 monolayers isolation

In order to perform Western blots, cell lines were grown to confluence in T-75 flasks (Thermo Scientific, UK) and left for 3-4 days, as previously described. The flask was then transferred to ice and the medium removed, before the cells were washed twice using ice-cold PBS+. Then, 1ml of ice-cold Radio-Immunoprecipitation Assay (RIPA) buffer (Sigma-Aldrich, Dorset, UK) with added protease inhibitors (10%v/v)(Thermo Scientific, Loughborough, UK) was added to the flask to lyse the cells. A plastic cell scraper (Greiner Bio-One Ltd, Gloucestershire, UK) was used to scrape the cells off the bottom of the flask and the cell lysate was transferred to a pre-cooled 1.5 ml Eppendorf tube which was left on ice for 20 minutes. The tubes were then centrifuged at 10,000 rpm for 10 minutes at 4°C using a Thermo Electron Corporation Heraeus Fresco17 bench-top micro-centrifuge. After centrifugation, the supernatant was transferred to another pre-cooled 1.5 ml Eppendorf and the pellet discarded. The resulting supernatant was taken for Western blot analysis.

### 4.7 Mouse brain capillary isolation

Brain capillaries from old-age (16 months) wild-type (3 females) and 3xTgAD (3 males, 1 female) mice were used to explore MATE1 expression. Brain capillaries were also isolated from elderly (24 month) age-matched wild-type C57BL6/129 mice (3 males, 2 females) and 3xTgAD mice (3 males, 2 females) for all other transporter studies. The left ventricle of the heart was cannulated and perfused (5ml/min) with an oxygenated artificial plasma (modified Krebs-Henseleit mammalian Ringer) for up to 2 minutes. The right atrium was sectioned before perfusion was started. The mice were then decapitated and the perfused brain removed. The brain was homogenized in physiological buffer (brain weightx3) and 26% dextran (brain weightx4). The homogenate was subjected to density gradient centrifugation (5,400xg for 15 min at 4°C) to give an endothelial cell-enriched pellet and the supernatant was discarded (Sanderson *et al.* 2007). 300µl of ice-cold RIPA: ThermoFisher Scientific cat#89900) bu[er with added protease inhibitors was added to the pellet at 4°C to lyse the tissue and then centrifuged at 8,000xg for 15minutes at 4°C. The resulting supernatant was taken for Western blot analysis.

### 4.8 Human tissue

Human tissue was provided via the brains for dementia research (BDR) and were anonymized. BDR has ethical approval granted by the national health service (NHS) health research authority (NRES Committee London-City & East, UK: REC reference:08/H0704/128+5. IRAS project ID:120436). Tissue was received on the basis that it will be handled, stored, used and disposed of within the terms of the Human Tissue Act 2004. Post-mortem brain capillaries from healthy individuals (Braak stage 0-II; 86.8±1.5 years; 2 females, 3 males) and AD cases (Braak stage V-VI; 79.4±3.7 years; 2 females, 3 males) were used to investigate the expression of transporters (Case details –Table S3). Medication history of the cases was supplied by the Manchester Brain Bank (Table S4). In this study we identified those drugs prescribed as sedatives, antidepressants and antipsychotics.

### 4.9 Human brain microvasculature isolation

Brain capillaries from frontal cortex, caudate nucleus, and putamen samples were isolated after homogenising 300mg tissue and carrying out a dextran based density-gradient centrifugation to produce a capillary-enriched pellet.

The pellet was further lysed with 500µl of ice-cold RIPA buffer with added protease inhibitors at 4°C and then centrifuged at 8,000xg for 15 minutes at 4°C. The resulting supernatant was taken for Western blot analysis to examine transporter expression.

### 4.10 Western Blot Procedure

The supernatant protein concentration was determined using a BCA assay (Albumin standard, ThermoScientific). The supernatants were diluted and boiled for 5 minutes at 95°C in 5x Laemmli sample buffer. Cell lines (30μg except for MATE 1 antibody in Bend.3 cells where 15μg was utilized and PMAT antibody in hCMEC/D3 and bEnd.3 cells where 20μg and 10μg was utilized respectively), mouse samples (15μg for MATE1, OCTN1 and 2) and (30μg for MATE2, PMAT and OCT1), human samples (10μg for OCNT1 and 2) or 15 μg (for MATE1, MATE2, PMAT and OCT1) were loaded equally on 4–20% Mini-PROTEAN® TGX™ gels (Bio-Rad) alongside a molecular weight marker (Precision plus protein, Bio-Rad). Samples underwent SDS-PAGE at 160V for 1 hour. Proteins were transferred onto 0.45μm polyvinylidene fluoride membranes (GE Healthcare, UK) after methanol activation at 100V for 1 hour. Membranes were blocked to reduce nonspecific binding using 5% milk with PBS-TWEEN® tablets (PBS-T) (Calbiochem, USA) at room temperature (RT) for 1 hour. Membranes were incubated overnight at 4°C with primary antibodies in PBS-T (Table S6). Membranes were washed in PBS-T (3×10 min) and incubated with the secondary antibody in PBS-T at RT for 1 hour. Further washing in PBS-T (3×10 min), membranes were then incubated with enhanced chemiluminescent reagent (ThermoScientific) for 30 seconds at RT.

Quantification of protein expression was determined by calculating the intensity ratio of the band of interest and the band of the loading control (tubulin or GAPDH). Band intensity ratio analysis was conducted using ImageJ software (NIH). Our group have previously published results from Pgp, BCRP, OCT1, OCT2 and OCT3 protein expression studies of bEnd.3 and hCMEC/D3 cells (Watson *et al.* 2012)(Sekhar *et al.* 2017).

### 4.11 Data analysis

Data are expressed as mean±SEM. The data was analysed by two-way ANOVA with Holm-Sidak post-hoc test for the accumulation studies and *in situ* perfusion studies, one-way ANOVA with Tukey’s post-hoc test for MTT assay and Student’s t-test or two-way ANOVA for Western blot data using Sigmaplot version 13 (Systat, USA) or GraphPad Prism 7.03. p <0.05 were considered as statistically significant. Exact p-values are provided in the figure legends/results section.

## Acknowledgements

This research was supported by the Medical Research Council (PhD studentship for Ms Sekhar PhD studies MR/K500811/1), the King’s College London Alzheimer’s Research UK (ARUK) network centre seed fund (to Drs Suzanne Reeves, Dr Sarah Thomas and Ms Gayathri Sekhar), The Edmond and Lily Safra Research Foundation (Dr Broadstock) and a multi-user equipment grant from The Wellcome Trust (Dr Thomas (PI), Professors Francis, McMahon, Malcangio, and Rattray 080268). Ms Fleckney is a BBSRC-CASE funded PhD student (BB/L01534X/1). Ms Sevda Boyanova is a Guy’s and St Thomas’ Charity funded MRes-PhD studentship. Mr Huzefa Rupawala is on a MRC DTP PhD studentship (MR/N013700/1). Dr Suzanne Reeves is funded by the UCLH NIHR BRC. The human brain tissue samples were supplied by The Manchester Brain Bank, which is part of the Brains for Dementia Research programme jointly funded by Alzheimer’s Research UK and Alzheimer’s Society (BDR Reference: Amendment to study-BDR14 TRID 170 recipient PI: Dr Sarah Thomas). We would like to thank Professor I. Romero, Professor B. Weksler, and Professor P. Couraud for the hCMEC/D3 cell line provided under MTA and Dr Manasi Nandi for advice regarding the Western blots.

## ABBREVIATIONS

(OCT): organic cation transporters
(OCTN): organic cation transporters novel
(hCMEC/D3): immortalized human cerebral microvessel endothelial cell line
(MATEs): multi-drug and toxic compound extrusion proteins
(PMAT): plasma membrane monoamine transporter, 3xTgAD
(FBS): foetal bovine serum
(BBB): blood-brain barrier
(AD): Alzheimer’s disease
(P-gp): P-glycoprotein
(BCRP): breast cancer resistance protein
(MRP): multi-drug resistance protein.

## Author contributions

**GNS** co-designed the study. Performed the brain perfusions (with AF). Performed MTT assay for amisulpride and eflornithine, Western blot analysis on human tissue and the bEnd.3 and D3 cell culture studies with unlabelled amisulpride, ATP depletion, ATP-transporter and OCT inhibitors. Analysed the resulting data. Contributed to the writing of the manuscript.

**ALF** performed the brain perfusions (with GNS). Performed Western blots for MATE1 in bEnd.3, hCMEC/D3, wild-type and 3xTg AD mice. Analysed the resulting data.

**STB** and **RL** performed the cell culture studies with MATE1, MATE2, PMAT, OCTN1 and OCTN2 inhibitors with hCMECD3 and bEnd.3 respectively. Analysed the resulting data.

**STB** also performed MTT assay with ergothionine, *L*-carnitine, famotidine, lopinavir, nifekalant hydrochloride and [^14^C]sucrose. PMAT Western blot on hCMEC/D3. Analysed the resulting data.

**HR** performed the Western blots for OCTN1, OCTN2 and MATE2 on bEnd.3, hCMEC/D3, wild-type and 3xTg AD mice. Also Western blot on OCTN1 and OCTN2 on human control and AD cases. Analysed the resulting data.

**HW** provided cell culture guidance to STB and RL.

**DBF and KMR** performed the *in silico* simulations, analysed the data and contributed to the writing of the manuscript.

**MB** provided the WT and 3xAD mice.

**SR** provided clinical insight into the project, separated the medications into the two groups and contributed to the writing of the manuscript.

**SAT** co-designed, directed and co-ordinated the complete study, developed the concept and wrote the majority of the manuscript. Applied for human tissue through the brains for dementia research fund. Contributed to analysing the data.

## Supplementary information

See separate document

## References

A A. Daniëlle M E van B. B. van van A., Mark L., Ronald B., Windhorst Albert D S. R. C., Lammertsma S. P. (2012) P-Glycoprotein Function at the Blood–Brain Barrier: Effects of Age and Gender. Mol. Imaging Biol. 14, 771–776.

Ahlin G., Karlsson J., Pedersen J. M., Gustavsson L., Larsson R., Matsson P., Norinder U., Bergström C. A. S., Artursson P. (2008) Structural requirements for drug inhibition of the liver specific human organic cation transport protein 1. J. Med. Chem. 51, 5932–42.

Alam A., Küng R., Kowal J., McLeod R. A., Tremp N., Broude E. V., Roninson I. B., Stahlberg H., Locher K. P. (2018) Structure of a zosuquidar and UIC2-bound human-mouse chimeric ABCB1. Proc. Natl. Acad. Sci. 115, E1973–E1982.

André P., Saubaméa B., Cochois-Guégan V., Marie-Claire C., Cattelotte J., Smirnova M., Schinkel A. H., Scherrmann J.-M., Cisternino S. (2012) Transport of biogenic amine neurotransmitters at the mouse blood-retina and blood-brain barriers by uptake1 and uptake2. J. Cereb. Blood Flow Metab. 32, 1989–2001.

Anthonypillai C., Gibbs J. E., Thomas S. A. (2006) The distribution of the anti-HIV drug, tenofovir (PMPA), into the brain, CSF and choroid plexuses. Cerebrospinal Fluid Res. 3, 1.

Ballard C., Howard R. (2006) Neuroleptic drugs in dementia: benefits and harm. Nat. Rev. Neurosci. 7, 492–500.

Bourdet D. L. (2005) Differential Substrate and Inhibitory Activities of Ranitidine and Famotidine toward Human Organic Cation Transporter 1 (hOCT1; SLC22A1), hOCT2 (SLC22A2), and hOCT3 (SLC22A3). J. Pharmacol. Exp. Ther. 315, 1288–1297.

Boyanova, S., Wang, H., Reeves, S. and Thomas S. A. (2018) The role of solute carrier transporters in the translocation of the antipsychotic amisulpride at the blood-brain barrier in Alzheimer’s disease and in normal ageing. Europhysiology, 248P.

Brown R. C., Morris A. P., O’Neil R. G. (2007) TIGHT JUNCTION PROTEIN EXPRESSION AND BARRIER PROPERTIES OF IMMORTALIZED MOUSE BRAIN MICROVESSEL ENDOTHELIAL CELLS. Brain Res. 1130, 17–30.

Caravaggio Fernando; Graff-Guerrero A. (2017) Is antipsychotic sensitivity in Alzheimer’s disease secondary to abnormal blood-brain barrier integrity? Brain 140, 865–867.

Clark-Papasavas C., Dunn J. T., Greaves S., Mogg A., Gomes R., Brownings S., Liu K., et al. (2014) Towards a therapeutic window of D2/3 occupancy for treatment of psychosis in Alzheimer’s disease, with [18F]fallypride positron emission tomography. Int. J. Geriatr. Psychiatry 29, 1001–9.

Coukell A. J., Spencer C. M., Benfield P. (1996) Amisulpride. CNS Drugs 6, 237–256.

Deo A. K., Borson S., Link J. M., Domino K., Eary J. F., Ke B., Richards T. L., et al. (2014) Activity of P-Glycoprotein, a β-Amyloid Transporter at the Blood-Brain Barrier, Is Compromised in Patients with Mild Alzheimer Disease. J. Nucl. Med. 55, 1106–11.

Dickens D., Owen A., Alfirevic A., Giannoudis A., Davies A., Weksler B., Romero I. A., Couraud P.-O., Pirmohamed M. (2012) Lamotrigine is a substrate for OCT1 in brain endothelial cells. Biochem. Pharmacol. 83, 805–814.

Haar H. J. van de, Burgmans S., Jansen J. F. A., Osch M. J. P. van, Buchem M. A. van, Muller M., Hofman P. A. M., Verhey F. R. J., Backes W. H. (2016) Blood-Brain Barrier Leakage in Patients with Early Alzheimer Disease. Radiology 281, 527–535.

Härtter S., Hüwel S., Lohmann T., Abou el ela A., Langguth P., Hiemke C., Galla H.-J. (2003) How Does the Benzamide Antipsychotic Amisulpride get into the Brain?—An In Vitro Approach Comparing Amisulpride with Clozapine. Neuropsychopharmacology 28, 1916–1922.

Hiasa M., Matsumoto T., Komatsu T., Moriyama Y. (2006) Wide variety of locations for rodent MATE1, a transporter protein that mediates the final excretion step for toxic organic cations. Am. J. Physiol. - Cell Physiol. 291, C678–C686.

Hiemke C., Baumann P., Bergemann N., Conca A., Dietmaier O., Egberts K., Fric M., et al. (2011) AGNP consensus guidelines for therapeutic drug monitoring in psychiatry: update 2011. Pharmacopsychiatry 44, 195–235.

Hirata-Fukae C., Li H.-F., Hoe H.-S., Gray A. J., Minami S. S., Hamada K., Niikura T., et al. (2008) Females exhibit more extensive amyloid, but not tau, pathology in an Alzheimer transgenic model. Brain Res. 1216, 92–103.

Horwood N., Davies D. C. (1994) Immunolabelling of hippocampal microvessel glucose transporter protein is reduced in Alzheimer’s disease. Virchows Arch. 425, 69–72.

Itagaki S., Ganapathy V., Ho H. T. B., Zhou M., Babu E., Wang J. (2012) Electrophysiological characterization of the polyspecific organic cation transporter plasma membrane monoamine transporter. Drug Metab. Dispos. 40, 1138–43.

Ito S., Kusuhara H., Yokochi M., Toyoshima J., Inoue K., Yuasa H., Sugiyama Y. (2012) Competitive inhibition of the luminal efflux by multidrug and toxin extrusions, but not basolateral uptake by organic cation transporter 2, is the likely mechanism underlying the pharmacokinetic drug-drug interactions caused by cimetidine in the kidney. J. Pharmacol. Exp. Ther. 340, 393–403.

Iwaki K., Sakaeda T., Kakumoto M., Nakamura T., Komoto C., Okamura N., Nishiguchi K., Shiraki T., Horinouchi M., Okumura K. (2006) Haloperidol is an inhibitor but not substrate for MDR1/P-glycoprotein. J. Pharm. Pharmacol. 58, 1617–22.

Kania K. D., Wijesuriya H. C., Hladky S. B., Barrand M. A. (2011) Beta amyloid effects on expression of multidrug efflux transporters in brain endothelial cells. Brain Res. 1418, 1–11.

Kurosawa T., Tega Y., Higuchi K., Yamaguchi T., Nakakura T., Mochizuki T., Kusuhara H., Kawabata K., Deguchi Y. (2018) Expression and Functional Characterization of Drug Transporters in Brain Microvascular Endothelial Cells Derived from Human Induced Pluripotent Stem Cells. Mol. Pharm. 15, 5546–5555.

Lako I. M., Heuvel E. R. van den, Knegtering H., Bruggeman R., Taxis K. (2013) Estimating Dopamine D2 Receptor Occupancy for Doses of 8 Antipsychotics. J. Clin. Psychopharmacol. 33, 675–681.

Leon J. de, Diaz F. J., Wedlund P., Josiassen R. C., Cooper T. B., Simpson G. M. (2004) Haloperidol half-life after chronic dosing. J. Clin. Psychopharmacol. 24, 656–60.

Loryan I., Melander E., Svensson M., Payan M., K?nig F., Jansson B., Hammarlund-Udenaes M. (2016) In-depth neuropharmacokinetic analysis of antipsychotics based on a novel approach to estimate unbound target-site concentration in CNS regions: link to spatial receptor occupancy. Mol. Psychiatry 21, 1527–1536.

Matsson P., Pedersen J. M., Norinder U., Bergström C. A. S., Artursson P. (2009) Identification of novel specific and general inhibitors of the three major human ATP-binding cassette transporters P-gp, BCRP and MRP2 among registered drugs. Pharm. Res. 26, 1816–31.

Mauri M. C., Paletta S., Maffini M., Colasanti A., Dragogna F., Pace C. Di, Altamura A. C. (2014) Clinical pharmacology of atypical antipsychotics. EXCLI J. 13.

Natesan S., Reckless G. E., Barlow K. B. L., Nobrega J. N., Kapur S. (2008) Amisulpride the ‘atypical’ atypical antipsychotic — Comparison to haloperidol, risperidone and clozapine. Schizophr. Res. 105, 224–235.

Nies A. T., Koepsell H., Damme K., Schwab M. (2011) Organic cation transporters (OCTs, MATEs), in vitro and in vivo evidence for the importance in drug therapy. Handb. Exp. Pharmacol. 201, 105–67.

Oddo S., Caccamo A., Shepherd J. D., Murphy M. P., Golde T. E., Kayed R., Metherate R., Mattson M. P., Akbari Y., LaFerla F. M. (2003) Triple-transgenic model of Alzheimer’s disease with plaques and tangles: intracellular Abeta and synaptic dysfunction. Neuron 39, 409–21.

Park R., Kook S.-Y., Park J.-C., Mook-Jung I. (2014) Aβ1-42 reduces P-glycoprotein in the blood-brain barrier through RAGE-NF-κB signaling. Cell Death Dis. 5, e1299.

Poller B., Gutmann H., Krähenbühl S., Weksler B., Romero I., Couraud P.-O., Tuffin G., Drewe J., Huwyler J. (2008) The human brain endothelial cell line hCMEC/D3 as a human blood-brain barrier model for drug transport studies. J. Neurochem. 107, 1358–1368.

Rae E. A., Brown R. E. (2015) The problem of genotype and sex differences in life expectancy in transgenic AD mice.

Reeves S., Bertrand J., D’Antonio F., McLachlan E., Nair A., Brownings S., Greaves S., Smith A., Taylor D., Howard R. (2016) A population approach to characterise amisulpride pharmacokinetics in older people and Alzheimer’s disease. Psychopharmacology (Berl). 233, 3371–3381.

Reeves S., Eggleston K., Cort E., McLachlan E., Brownings S., Nair A., Greaves S., et al. (2018) Therapeutic D2/3 receptor occupancies and response with low amisulpride blood concentrations in very late-onset schizophrenia-like psychosis (VLOSLP). Int. J. Geriatr. Psychiatry 33, 396–404.

Reeves S., McLachlan E., Bertrand J., Antonio F. D., Brownings S., Nair A., Greaves S., et al. (2017) Therapeutic window of dopamine D2/3 receptor occupancy to treat psychosis in Alzheimer’s disease. Brain 140, 1117–1127.

Saidijam M., Karimi Dermani F., Sohrabi S., Patching S. G. (2017) Efflux proteins at the blood–brain barrier: review and bioinformatics analysis. Xenobiotica, 1–27.

Sanderson L., Dogruel M., Rodgers J., Bradley B., Thomas S. A. (2008) The blood-brain barrier significantly limits eflornithine entry into Trypanosoma brucei brucei infected mouse brain. J. Neurochem. 107, 1136–1146.

Sanderson L., Khan A., Thomas S. (2007) Distribution of suramin, an antitrypanosomal drug, across the blood-brain and blood-cerebrospinal fluid interfaces in wild-type and Pglycoprotein transporter-deficient mice. Antimicrob. Agents Chemother. 51, 3136–3146.

Santos Pereira J. N. Dos, Tadjerpisheh S., Abu Abed M., Saadatmand A. R., Weksler B., Romero I. A., Couraud P.-O., Brockmöller J., Tzvetkov M. V (2014) The poorly membrane permeable antipsychotic drugs amisulpride and sulpiride are substrates of the organic cation transporters from the SLC22 family. AAPS J. 16, 1247–58.

Schinkel A. H., Wagenaar E., Mol C. A., Deemter L. van (1996) P-glycoprotein in the blood-brain barrier of mice influences the brain penetration and pharmacological activity of many drugs. J. Clin. Invest. 97, 2517–24.

Schmitt U., Kirschbaum K. M., Poller B., Kusch-Poddar M., Drewe J., Hiemke C., Gutmann H. (2012) In vitro P-glycoprotein efflux inhibition by atypical antipsychotics is in vivo nicely reflected by pharmacodynamic but less by pharmacokinetic changes. Pharmacol. Biochem. Behav. 102, 312–20.

Schneider L. S., Tariot P. N., Dagerman K. S., Davis S. M., Hsiao J. K., Ismail M. S., Lebowitz B. D., et al. (2006) Effectiveness of Atypical Antipsychotic Drugs in Patients with Alzheimer’s Disease. N. Engl. J. Med. 355, 1525–1538.

Schoemaker H., Claustre Y., Fage D., Rouquier L., Chergui K., Curet O., Oblin A., et al. (1997) Neurochemical characteristics of amisulpride, an atypical dopamine D2/D3 receptor antagonist with both presynaptic and limbic selectivity. J. Pharmacol. Exp. Ther. 280, 83–97.

Schwake M., Schröder B., Saftig P. (2013) Lysosomal membrane proteins and their central role in physiology. Traffic 14, 739–48.

Sekhar, G., Reeves, S. and Thomas S. A. (2015) Exploring the interaction of amisulpride with influx and efflux transporters expressed at the blood-brain barrier. Pharmacology 12, 173P.

Sekhar G. N., Georgian A. R., Sanderson L., Vizcay-Barrena G., Brown R. C., Muresan P., Fleck R. A., Thomas S. A. (2017) Organic cation transporter 1 (OCT1) is involved in pentamidine transport at the human and mouse blood-brain barrier (BBB). PLoS One 12.

Shimomura K., Okura T., Kato S., Couraud P.-O., Schermann J.-M., Terasaki T., Deguchi Y. (2013) Functional expression of a proton-coupled organic cation (H+/OC) antiporter in human brain capillary endothelial cell line hCMEC/D3, a human blood-brain barrier model. Fluids Barriers CNS 10.

Skrobecki P., Chmieli?ska A., Bonarek P., Stepien P., Wisniewska-Becker A., Dziedzicka-Wasylewska M., Polit A. (2017) Sulpiride, Amisulpride, Thioridazine, and Olanzapine: Interaction with Model Membranes. Thermodynamic and Structural Aspects. ACS Chem. Neurosci., acschemneuro.7b00057.

Sparshatt A., Taylor D., Patel M. X., Kapur S. (2009) Amisulpride - dose, plasma concentration, occupancy and response: implications for therapeutic drug monitoring. Acta Psychiatr. Scand. 120, 416–428.

Staud F., Cerveny L., Ahmadimoghaddam D., Ceckova M. (2013) Multidrug and toxin extrusion proteins (MATE/SLC47); role in pharmacokinetics. Int. J. Biochem. Cell Biol. 45, 2007–11.

Tsuda M., Terada T., Ueba M., Sato T., Masuda S., Katsura T., Inui K. (2009) Involvement of human multidrug and toxin extrusion 1 in the drug interaction between cimetidine and metformin in renal epithelial cells. J. Pharmacol. Exp. Ther. 329, 185–91.

Vogelgesang S., Cascorbi I., Schroeder E., Pahnke J., Kroemer H. K., Siegmund W., Kunert-Keil C., Walker L. C., Warzok R. W. (2002) Deposition of Alzheimer’s beta-amyloid is inversely correlated with P-glycoprotein expression in the brains of elderly nondemented humans. Pharmacogenetics 12, 535–41.

Watson C. P., Dogruel M., Mihoreanu L., Begley D. J., Weksler B. B., Couraud P. O., Romero I. A., Thomas S. A. (2012) The transport of nifurtimox, an anti-trypanosomal drug, in an in vitro model of the human blood–brain barrier: Evidence for involvement of breast cancer resistance protein. Brain Res. 1436, 111–121.

Watson C. P., Pazarentzos E., Fidanboylu M., Padilla B., Brown R., Thomas S. A. (2016) The transporter and permeability interactions of asymmetric dimethylarginine (ADMA) and L-arginine with the human blood–brain barrier in vitro. Brain Res. 1648, 232–242.

Weksler B., Romero I., Couraud P.-O. (2013) The hCMEC/D3 cell line as a model of the human blood brain barrier. Fluids Barriers CNS 10, 16.

Wijesuriya H. C., Bullock J. Y., Faull R. L. M., Hladky S. B., Barrand M. A. (2010) ABC efflux transporters in brain vasculature of Alzheimer’s subjects. Brain Res. 1358, 228–238.

Wittwer M. B., Zur A. A., Khuri N., Kido Y., Kosaka A., Zhang X., Morrissey K. M., Sali A., Huang Y., Giacomini K. M. (2013) Discovery of Potent, Selective Multidrug and Toxin Extrusion Transporter 1 (MATE1, SLC47A1) Inhibitors Through Prescription Drug Profiling and Computational Modeling. J. Med. Chem. 56, 781–795.

Wu K.-C., Lu Y.-H., Peng Y.-H., Hsu L.-C., Lin C.-J. (2015) Effects of lipopolysaccharide on the expression of plasma membrane monoamine transporter (PMAT) at the blood-brain barrier and its implications to the transport of neurotoxins. J. Neurochem.

