## Supplementary information for "Region-specific blood-brain barrier transporter changes leads to increased sensitivity to amisulpride in Alzheimer’s disease"

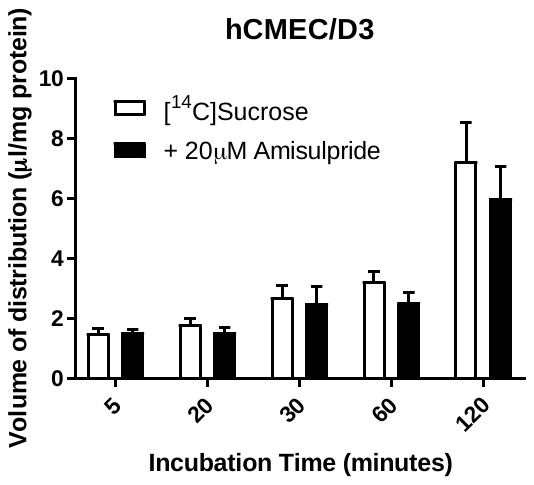

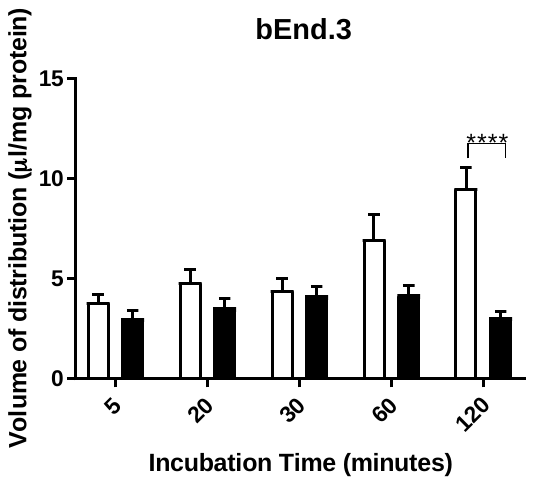

**Figure S1:** The effect of self-inhibition on the accumulation of [^14^C]sucrose was determined in hCMEC/D3 and bEnd.3 cell lines. No differences compared to control was observed, except at 120 minutes in b.End3 cells (p<0.0001). All data are expressed as mean ± S.E.M, n = 3 to 7 plates with 6 replicates (wells) per timepoint per plate. 5 time points.

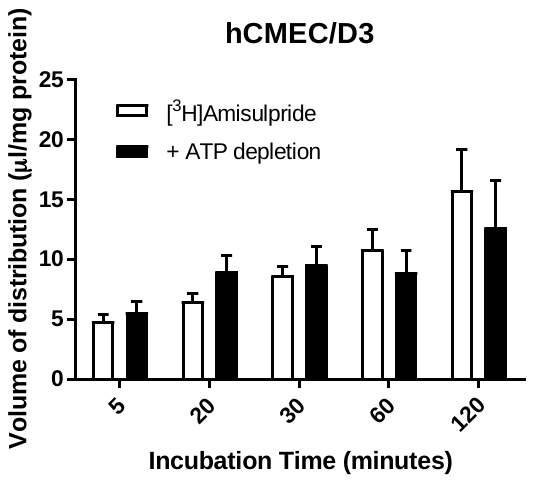

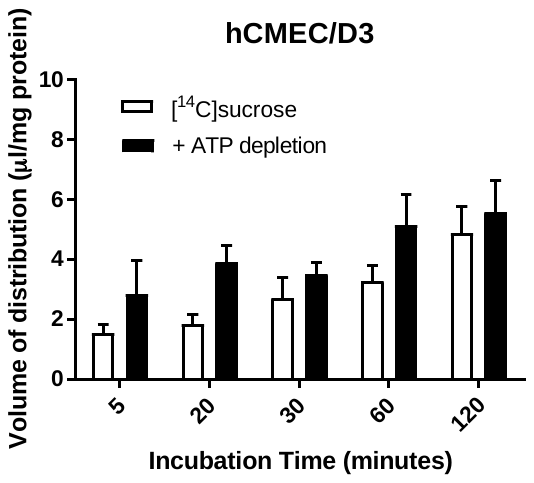

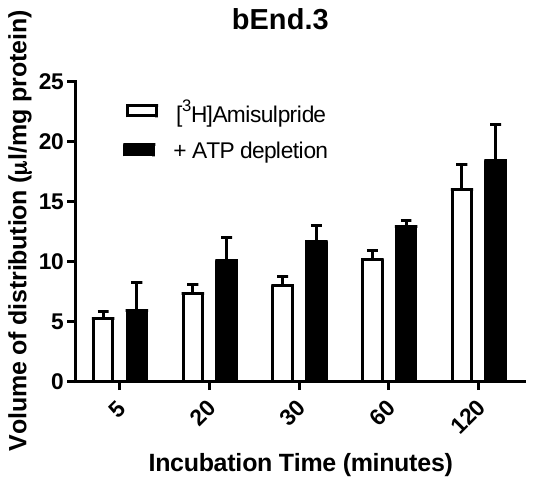

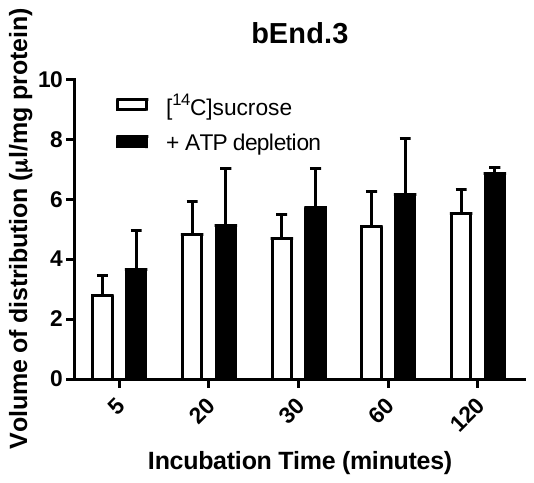

**Figure S2.** The effect of ATP depletion on the accumulation of [^3^H]amisulpride (6.5nM) and [^14^C]sucrose was determined in hCMEC/D3 (upper) and bEnd.3 (lower) cell lines. No significant differences were observed compared to control i.e. radiolabelled molecule alone. The [^3^H]amisulpride data has been corrected for [^14^C]sucrose and are expressed as mean ± S.E.M, n = 3 passages with 6 replicates (wells) per timepoint per plate. 5 time points. Data were analysed with SigmaPlot version 13.

| 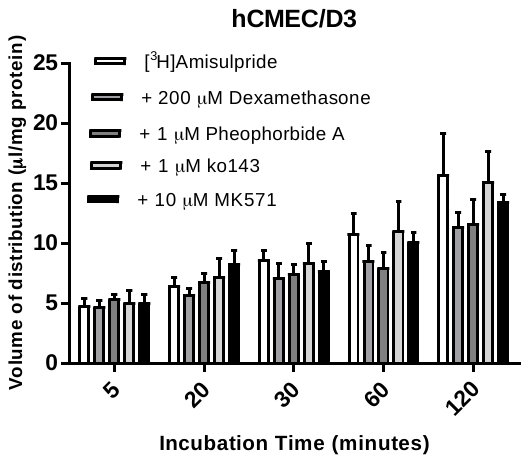 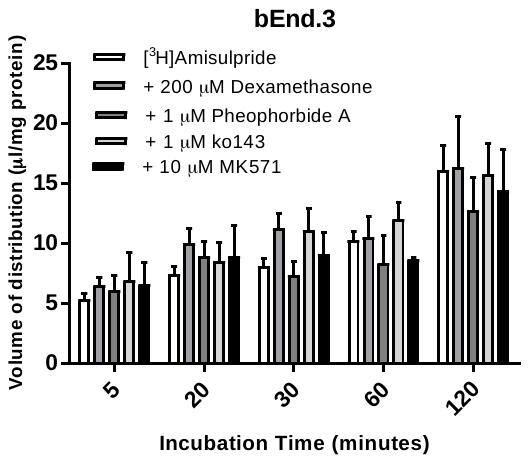 | 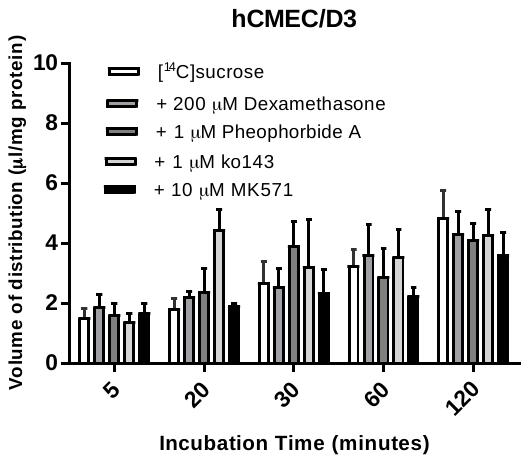 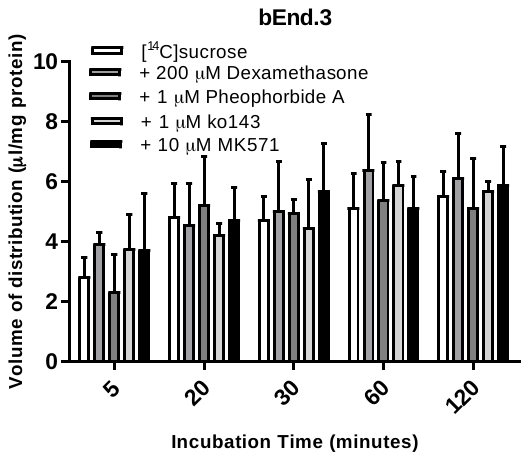 |
| --- | --- |

**Figure S3:** The role of ABC transporters on the accumulation of [^3^H]amisulpride (6.5nM) and [^14^C]sucrose was determined in hCMEC/D3 (upper) and bEnd.3 (lower) cell lines. No significant differences were observed compared to control i.e. radiolabelled molecule alone. The [^3^H]amisulpride data has been corrected for [^14^C]sucrose and are expressed as mean ± S.E.M, n = 3 passages with 6 replicates (wells) per timepoint per plate. 5 time points.

| 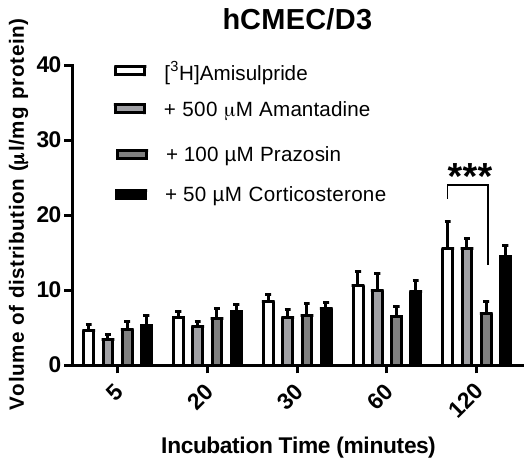 | 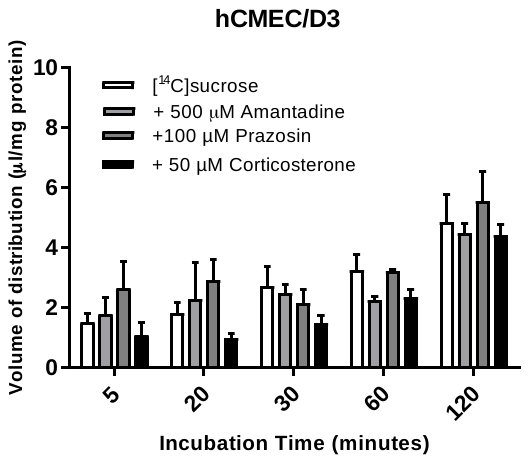 |
| --- | --- |
| 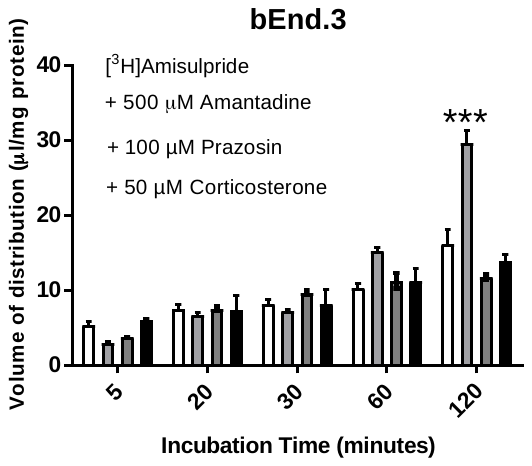 | 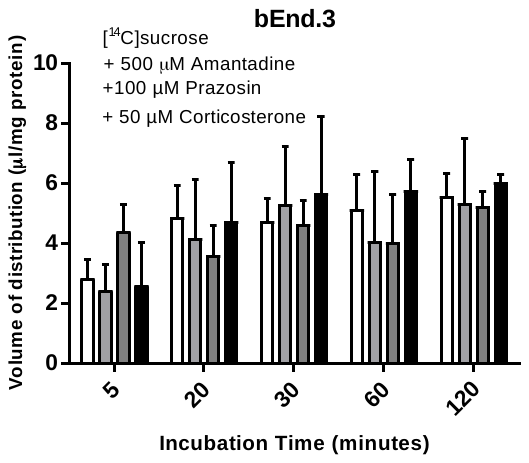 |

**Figure S4**: The effect of OCT-inhibition on the accumulation of [^3^H]amisulpride (6.5nM) and [^14^C]sucrose was determined in hCMEC/D3 and bEnd.3 cell lines. Significant differences were observed compared to control - ***p<0.001. [^3^H]amisulpride data has been corrected for [^14^C]sucrose All data are expressed as mean ± S.E.M, n = 3 passages with 6 replicates (wells) per timepoint per plate. 5 timepoints.

| 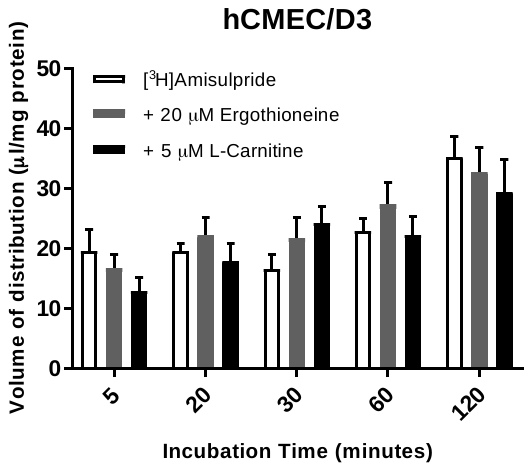 | 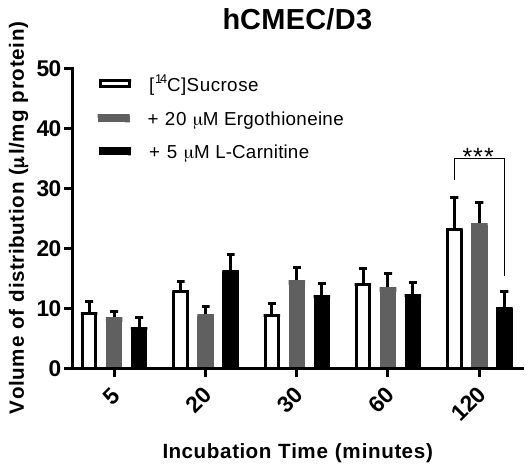 |
| --- | --- |

**Figure S5**: The effect of OCTN-inhibition on the accumulation of [^3^H]amisulpride (3.7-7.7nM) and [^14^C]sucrose (0.7-1.5 µM) was determined in hCMEC/D3. No differences were observed when inhibitor groups were compared to [^3^H]amisulpride control at all time points and [^14^C]sucrose control up to 60 minutes. Significant differences were observed between [^14^C]sucrose and *L*-carnitine at 2 hours suggestive that this time point for [^3^H]amisulpride should be ignored. [^3^H]amisulpride data has been corrected for [^14^C]sucrose. All data are expressed as mean ± S.E.M. n =3 passages (p31, p32 and p34) with 6 replicates (wells) per timepoint per plate for [^3^H]amisulpride with ergothioneine (5 time-points). n = 4 passages (p30, p31, p32 and p34) with 6 replicates (wells) per timepoint per plate for [^3^H]amisulpride alone and with L-carnitine (5 time-points). Data were analysed with GraphPad Prism 7.03, where ns - P > 0.05; *** - p ≤ 0.001;.

**bEnd.3	bEnd.3**

| 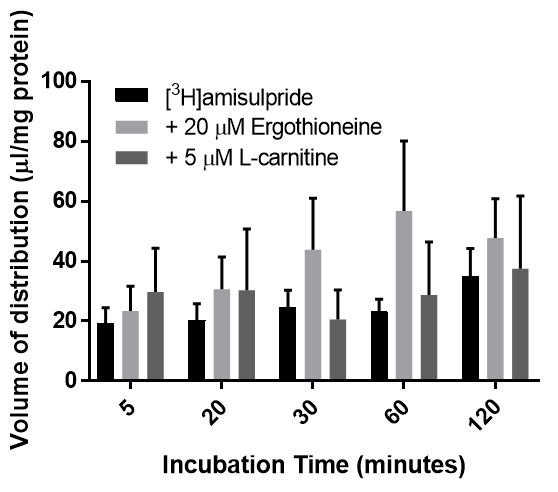 | 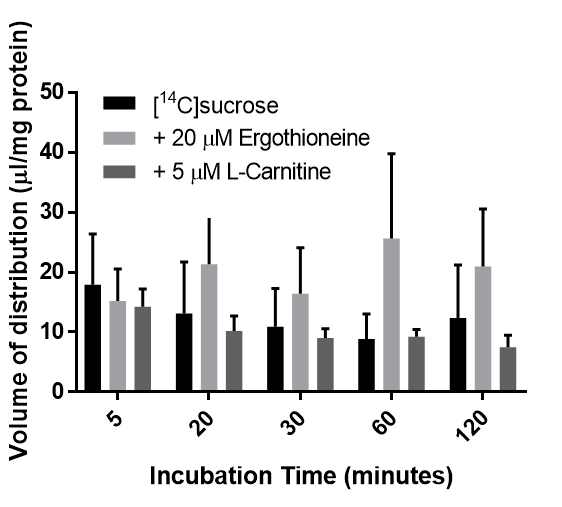 |
| --- | --- |

**Figure S6**. The effect of OCTN1 and OCTN2 substrates on [^3^H]amisulpride accumulation and [^14^C]sucrose values in bEnd.3 cells. Cells were incubated with OCTN1 substrate ergothioneine (20 µM) or OCTN substrate L-carnitine (5 µM). No significant differences in [^3^H]amisulpride accumulation or [^14^C]sucrose values were observed compared to control conditions following incubation for 5, 20, 30, 60, and 120 minutes. All data expressed as mean ± SEM, n = 3 (ergothioneine 20 µM, L-carnitine 5 µM) and n = 4 (control) passages of cells with 6 replicates per incubation timepoint per plate (5 time-points). Values in (A) have been corrected for sucrose values. Data were analysed by two-way ANOVA with Sidak post-hoc test using GraphPad Prism 7.

**bEnd.3	bEnd.3**

| 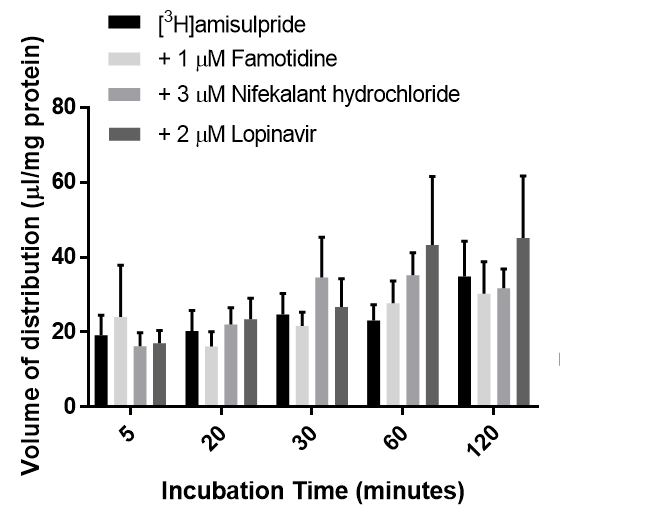 | 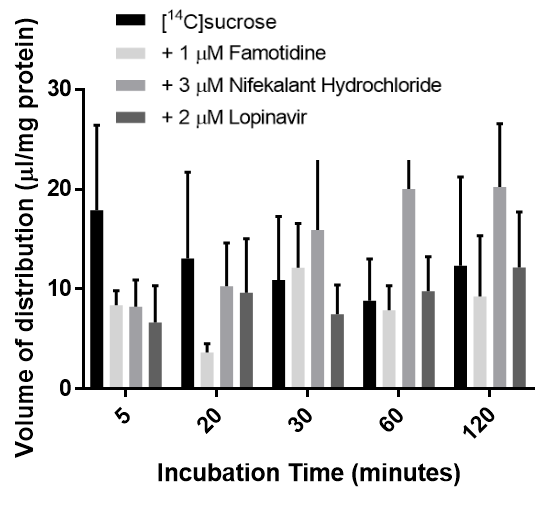 |
| --- | --- |

**Figure S7.** The effect of MATE1, MATE2 and PMAT inhibitors on [^3^H]amisulpride accumulation and [^14^C]sucrose values in bEnd.3 cells. Cells were incubated with MATE1 inhibitor, famotidine, MATE2 inhibitor, nifekalant hydrochloride (3 µM), or PMAT inhibitor, lopinavir (2 µM). No significant differences were observed compared to control conditions following incubation for 5, 20, 30, 60, and 120 minutes. All data expressed as mean ± SEM n = 3 (famotidine 1 µM) and n = 4 (control, nifekalant hydrochloride 3 µM, lopinavir 2 µM) passages of cells with 6 replicates per incubation time-point per plate. 5 time-points. [^3^H]amisulpride values have been corrected for [^14^C]sucrose values. Data were analysed by two-way ANOVA with Sidak post-hoc test using GraphPad Prism 7.

| 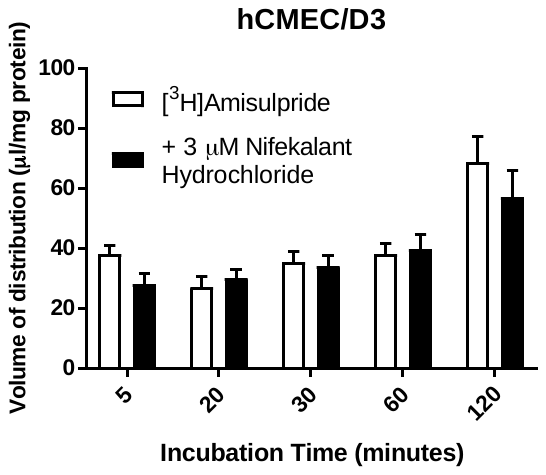 | 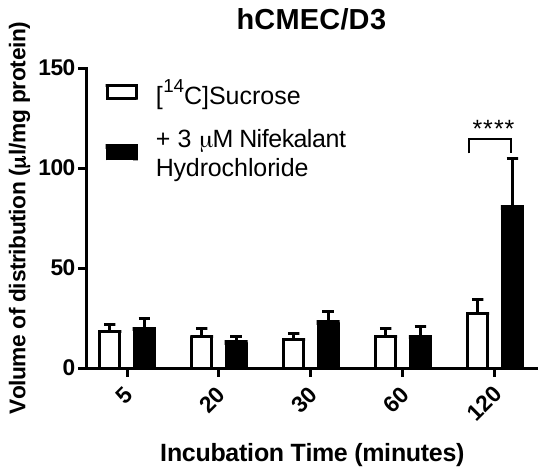 |
| --- | --- |

**Figure S8:** The effect of MATE2 inhibition on the accumulation of [^3^H]amisulpride (3.7-7.7nM) was determined in hCMEC/D3 cell lines. Significant differences were observed compared to [^14^C]sucrose control-****p<0.0001. [^3^H]amisulpride data has been corrected for [^14^C]sucrose and are expressed as mean ± S.E.M, n = 3 passages (p30, p31 and p34) with 6 replicates (wells) per timepoint per plate (5 time-points).

| 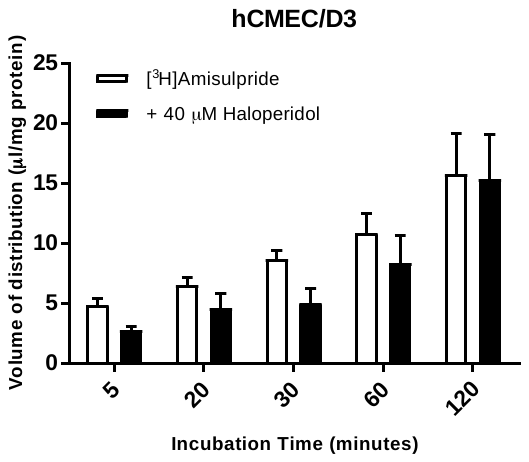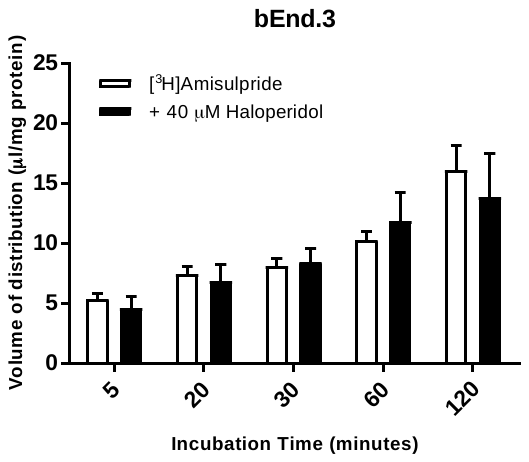 | 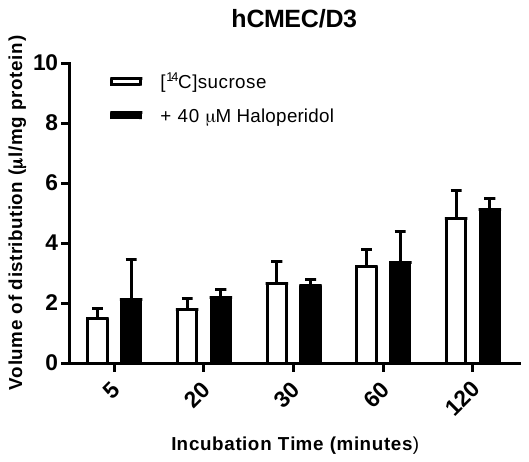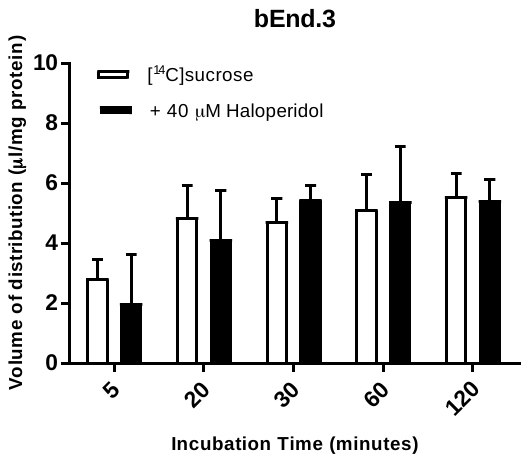 |
| --- | --- |

**Figure S9**: The effect of unlabelled haloperidol on the accumulation of [^3^H]amisulpride and [^14^C]sucrose

was determined in hCMEC/D3 (upper) and bEnd.3 (lower) cell lines. No significant differences were

observed compared to control. All data have been corrected for [^14^C]sucrose and are expressed as

mean ± S.E.M, n = 3 passages with 6 replicates (wells) per timepoint per plate. Data were

analysed with SigmaPlot version 13.

| 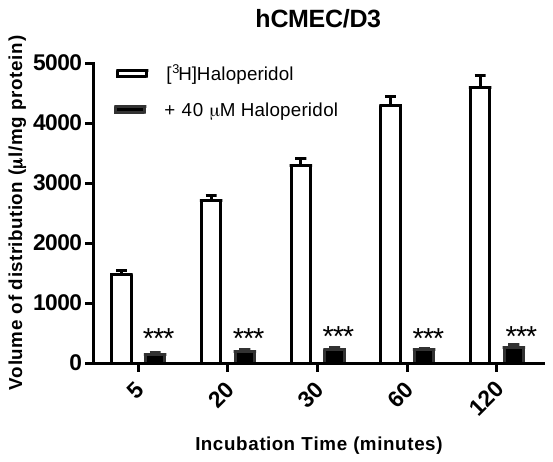 | 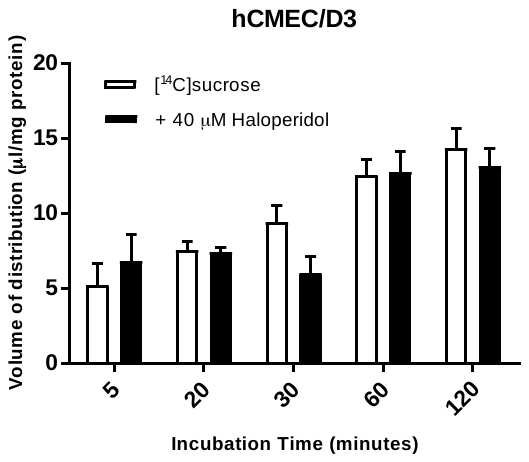 |
| --- | --- |
| 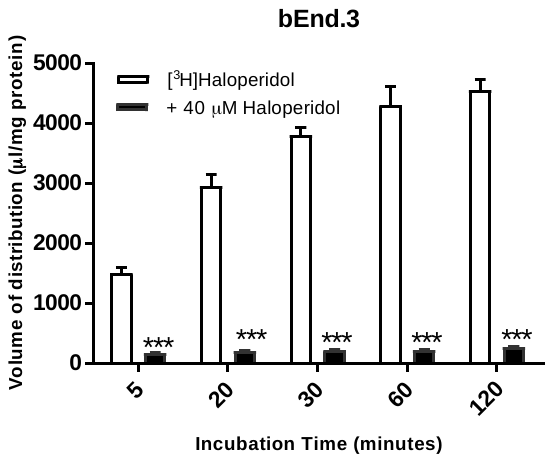 | 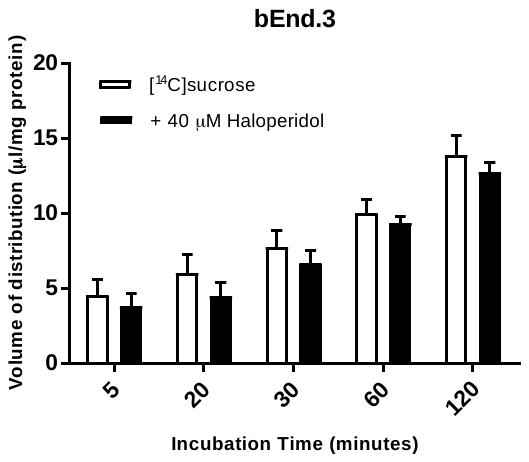 |

**Figure S10**: The effect of self-inhibition on the accumulation of 10 nM [^3^H]haloperidol and [^14^C]sucrose was investigated in hCMEC/D3 and bEnd.3 cell lines. Significant differences compared to control were observed - ***p<0.001. All [^3^H]haloperidol data have been corrected for [^14^C]sucrose. All data is expressed as mean ± S.E.M, n = 3 passages with 6 replicates (wells) per timepoint per plate. Data were analysed with SigmaPlot version 13.

| 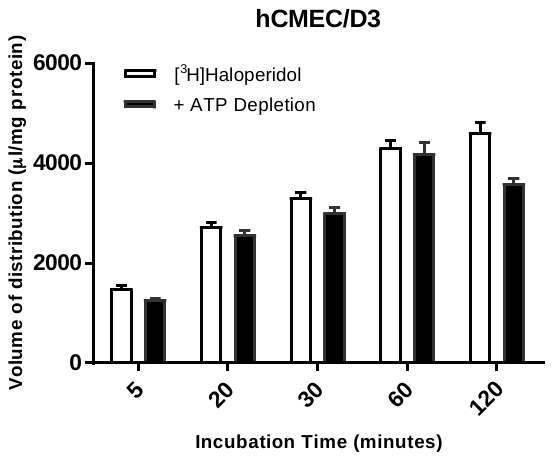 | 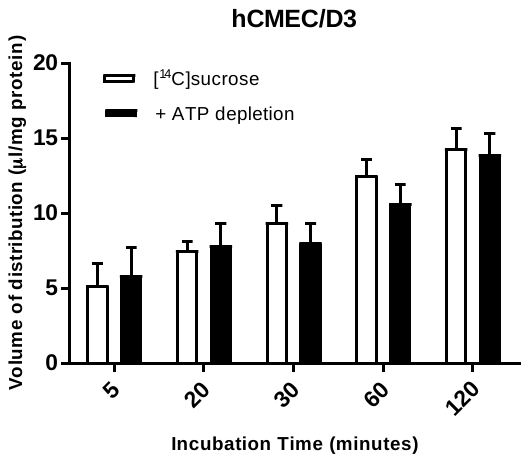 |
| --- | --- |
| 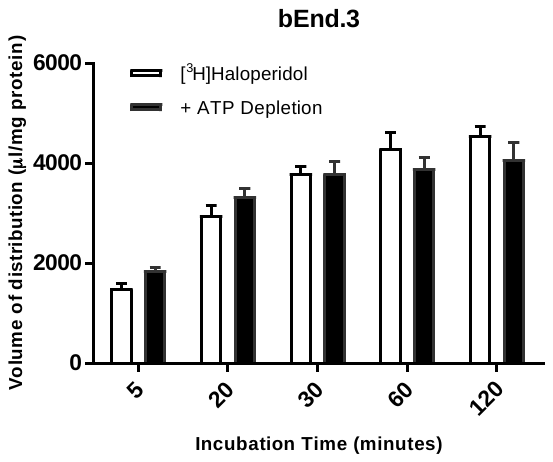 | 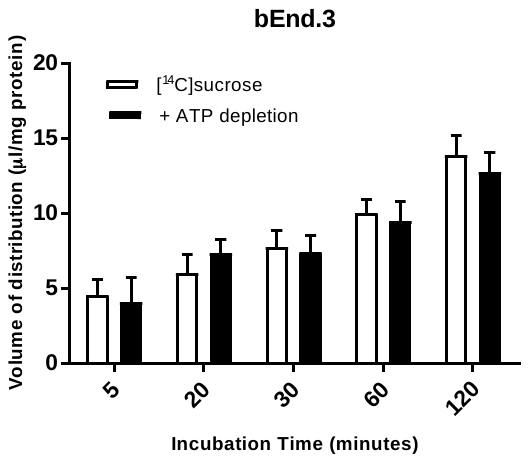 |

**Figure S11**: The effect of ATP depletion on the accumulation of 10nM [^3^H]haloperidol and 3.8μM [^14^C]sucrose was determined in hCMEC/D3 and bEnd.3 cell lines. No significant differences compared to control were observed. [^3^H]haloperidol data has been corrected for [^14^C]sucrose. All data is expressed as mean ± S.E.M, n = 3 passages with 6 replicates (wells) per timepoint per plate. Data were analysed with SigmaPlot version 13.

| 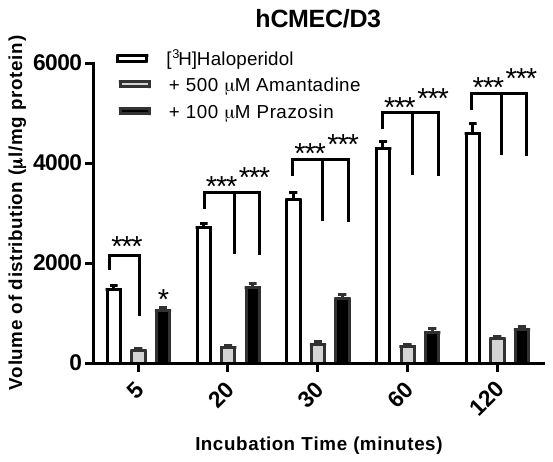 | 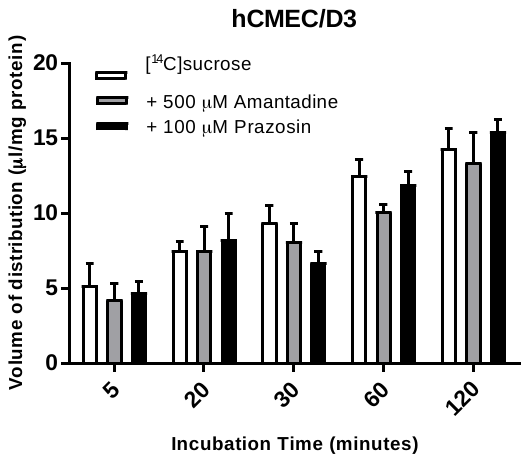 |
| --- | --- |
| 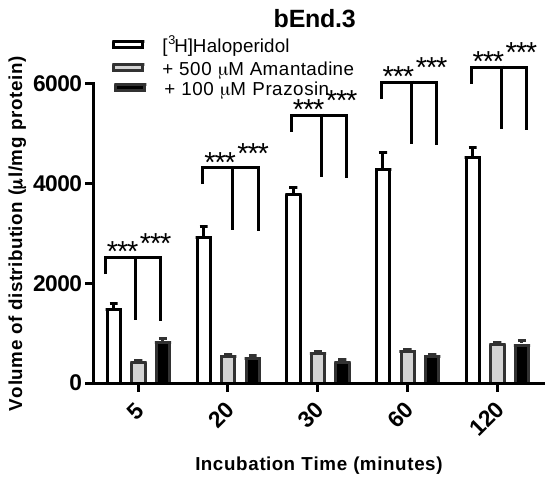 | 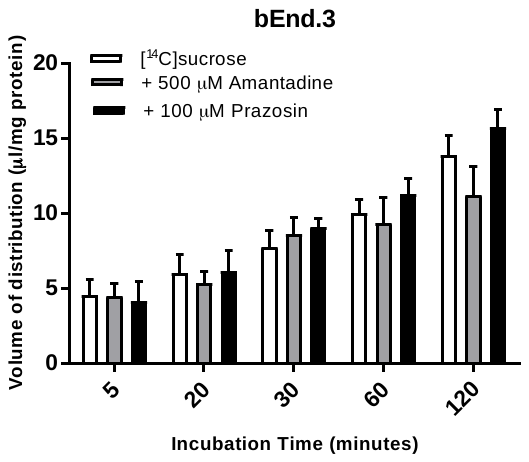 |

**Figure S12:** The effect of OCT inhibition on the accumulation of 10 nM [^3^H]haloperidol and 3.8μM [^14^C]sucrose in hCMEC/D3 and bEnd.3 cell lines. Significant differences were observed compared to control - ***p<0.001. [^3^H]haloperidol data has been corrected for [^14^C]sucrose and are expressed as mean ± S.E.M, n = 3 passages with 6 replicates (wells) per timepoint per plate. Data were analysed with SigmaPlot version 13.

| 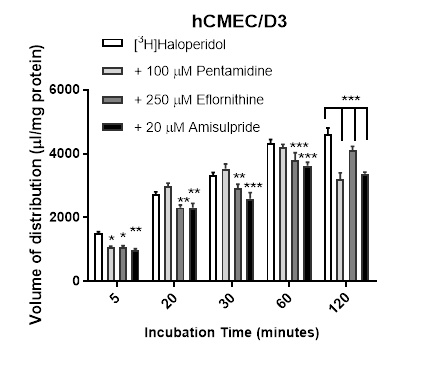 | 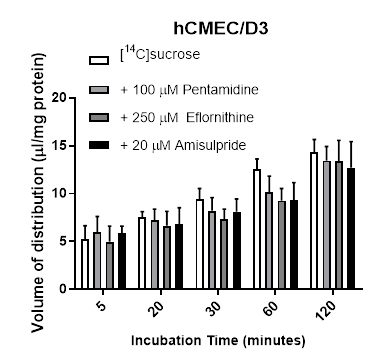 |
| --- | --- |

**Figure S13**: The effect of other cationic drugs on the accumulation of 10 nM [^3^H]haloperidol and 3.8μM [^14^C]sucrose. Significant differences were observed compared to control - ***p<0.001,**p<0.01, and *p<0.05. All data have been corrected for [^14^C]sucrose and are expressed as mean± S.E.M, n = 3 passages with 6 replicates (wells) per timepoint per plate. Data were analysed with SigmaPlot version.

13.

| **A.** | 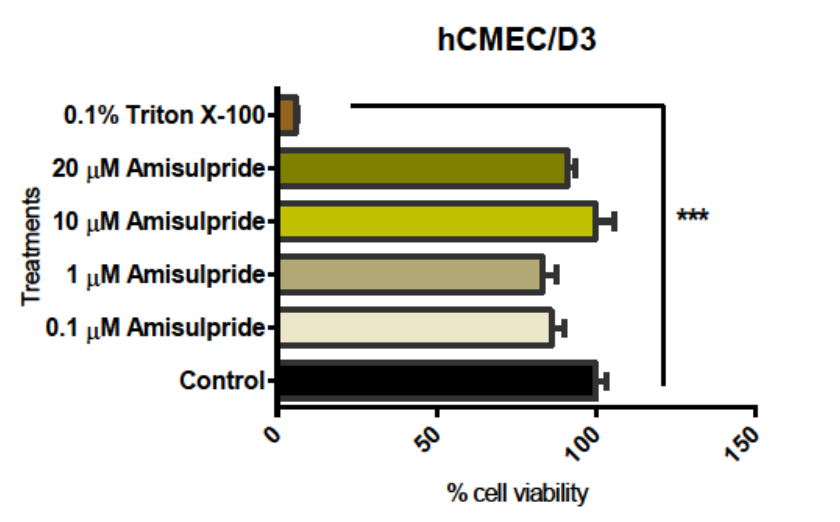 |
| --- | --- |
|  | 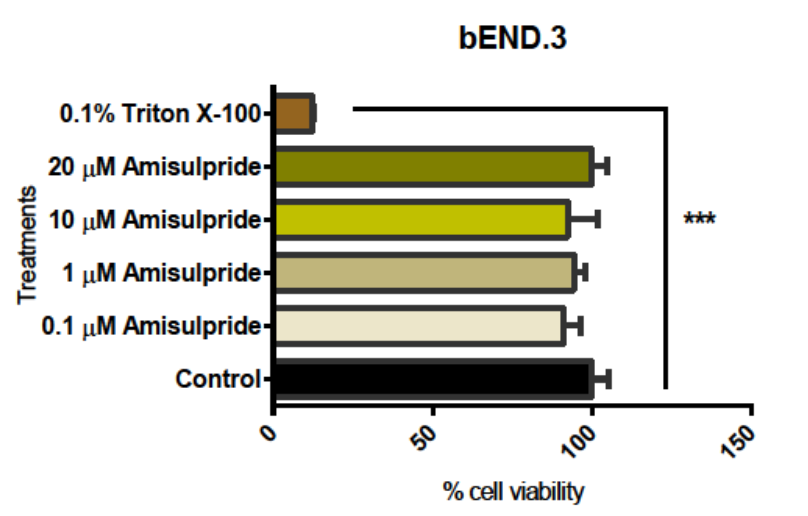 |
| **B.** | 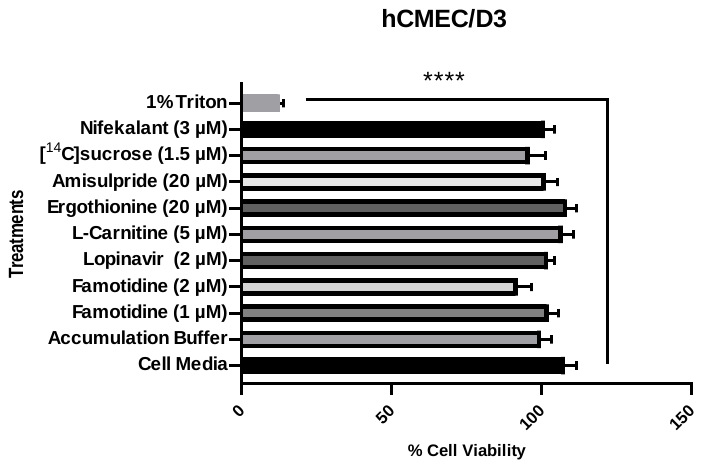 |

**Figure S14**: The cytotoxic effects of treatments on cells - no treatment except for the positive control Triton X-100 had a cytotoxic effect on cells (0.1%; ***p<0.001) or (1%; ****p<0.0001). All data are expressed as mean ± S.E.M [**A]** n = 3 passages with 6 replicates (wells) per timepoint per plate. Data were analysed with SigmaPlot version 13. **[B]** All data are expressed as mean ± S.EM., n=2 plates (P32 and P33) with 6 replicates (wells) per plate, except for [^14^C] sucrose (1.5µM) and nifekalant (3 µM) where n=1 plate (P33) was used. Data were analysed using mixed effects one-way ANOVA with GraphPad Prism 8.0.0.

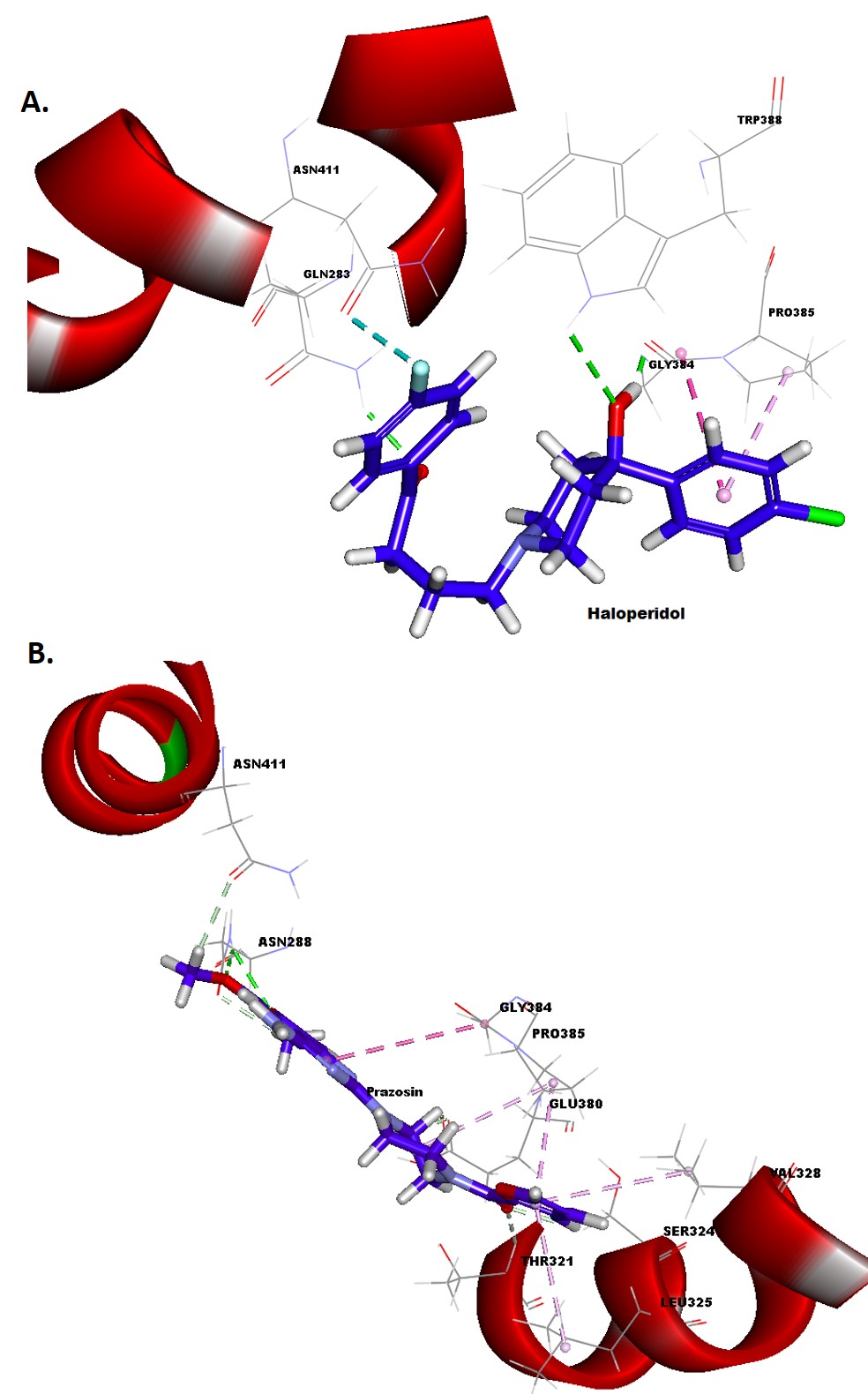

**Figure S15:** 3D molecular docking of A) haloperidol and B) prazosin with OCT-1 transporter showing the molecular level interactions. Hydrogen bonds are represented in green dotted lines, and hydrophobic interactions are represented in pink dotted lines.

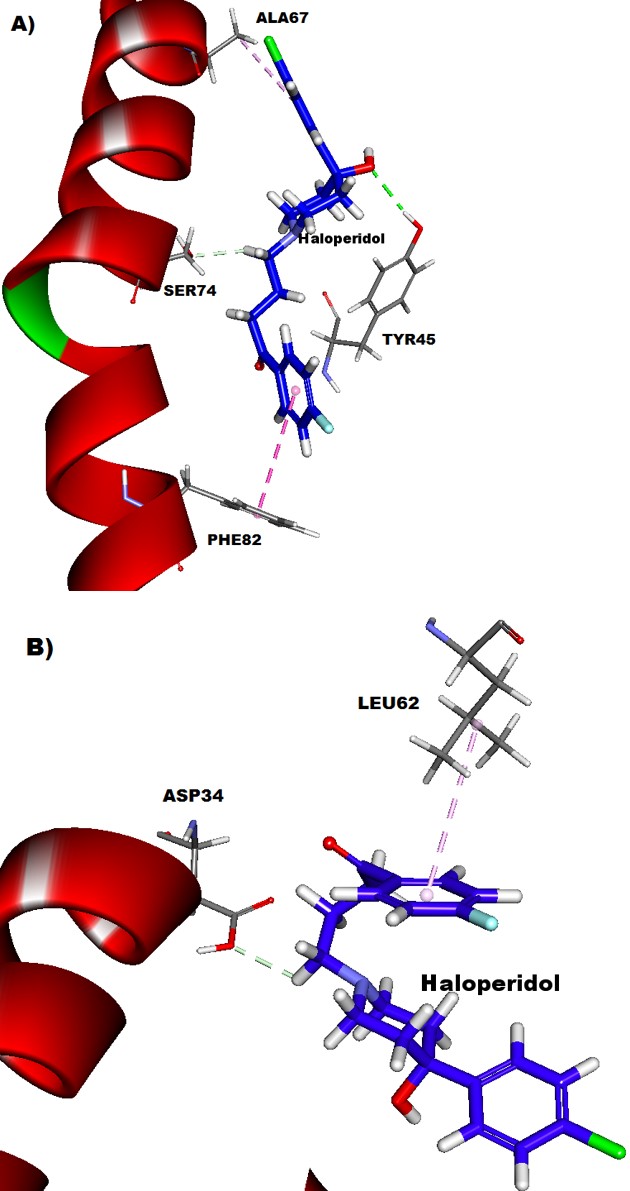

**Figure S16:** 3D molecular docking of haloperidol with A) MATE-1 and B) PMAT transporters showing the molecular level interactions. Hydrogen bonds are represented in green dotted lines, and hydrophobic interactions are represented in pink dotted lines.

A) **Amisulpride docking images**

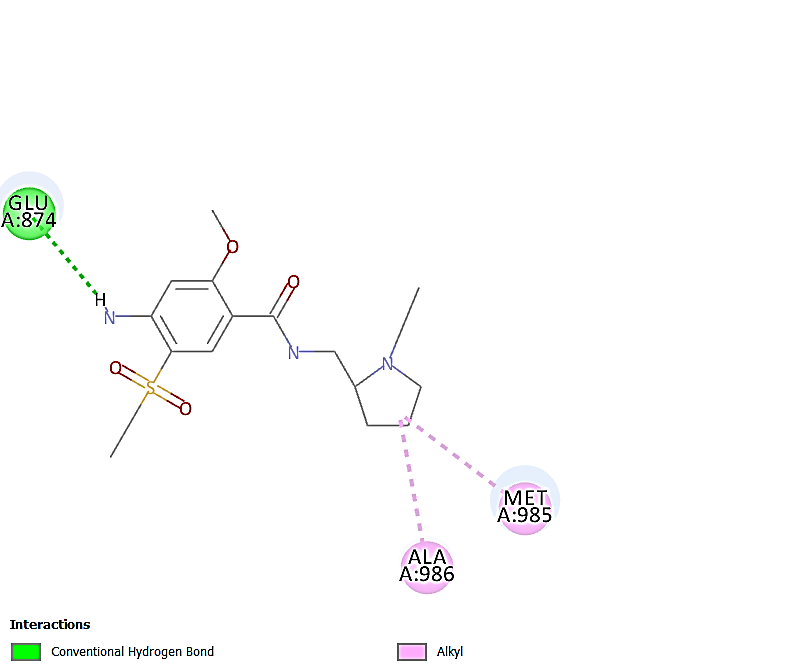

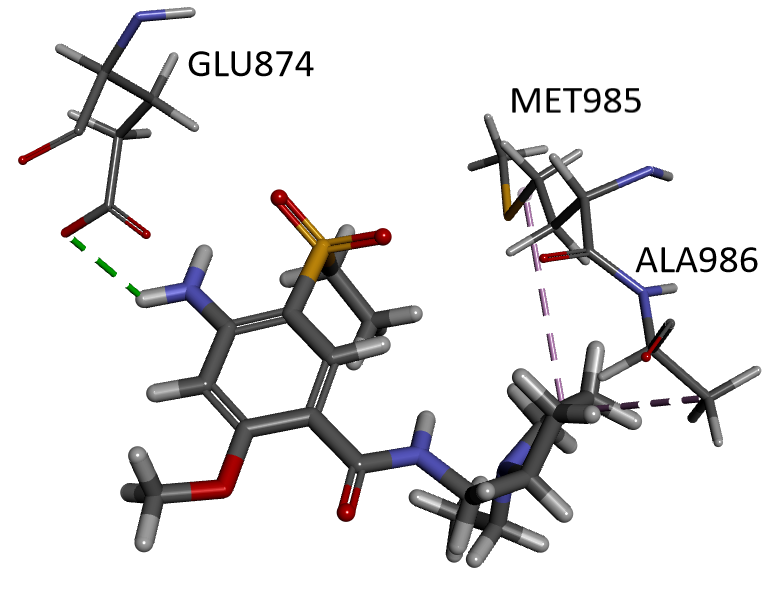

B) **Colchicine docking images**

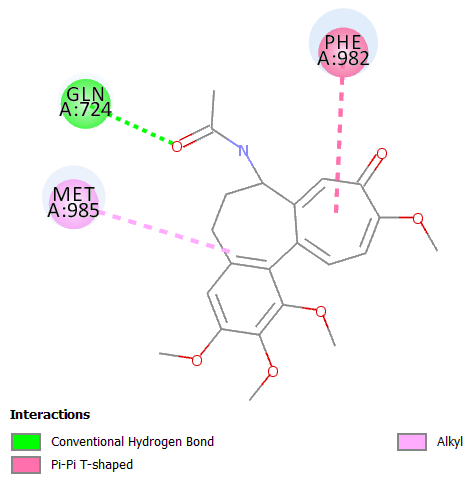

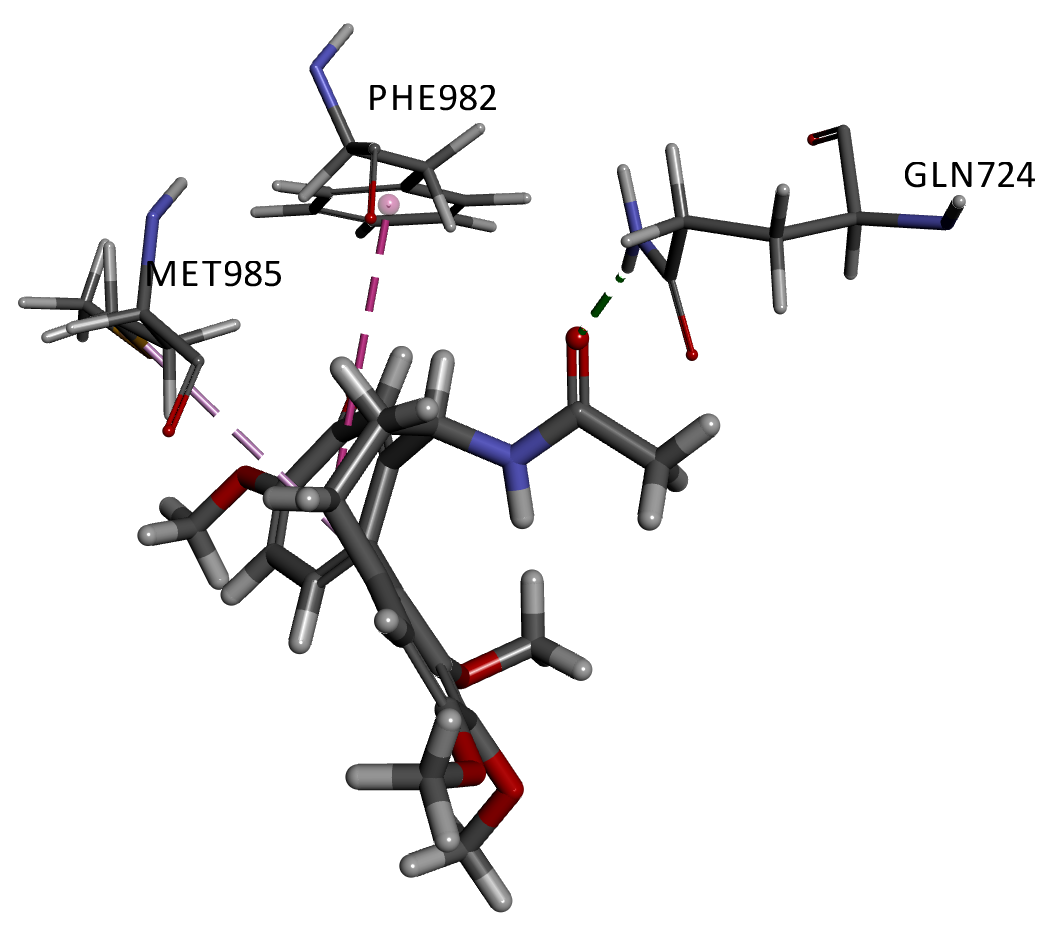

C) **Dexamethasone docking images**

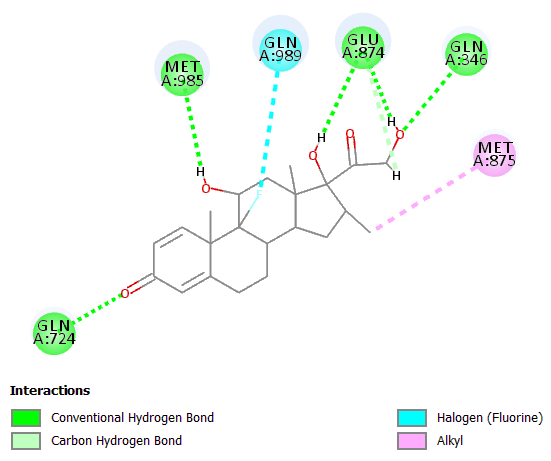

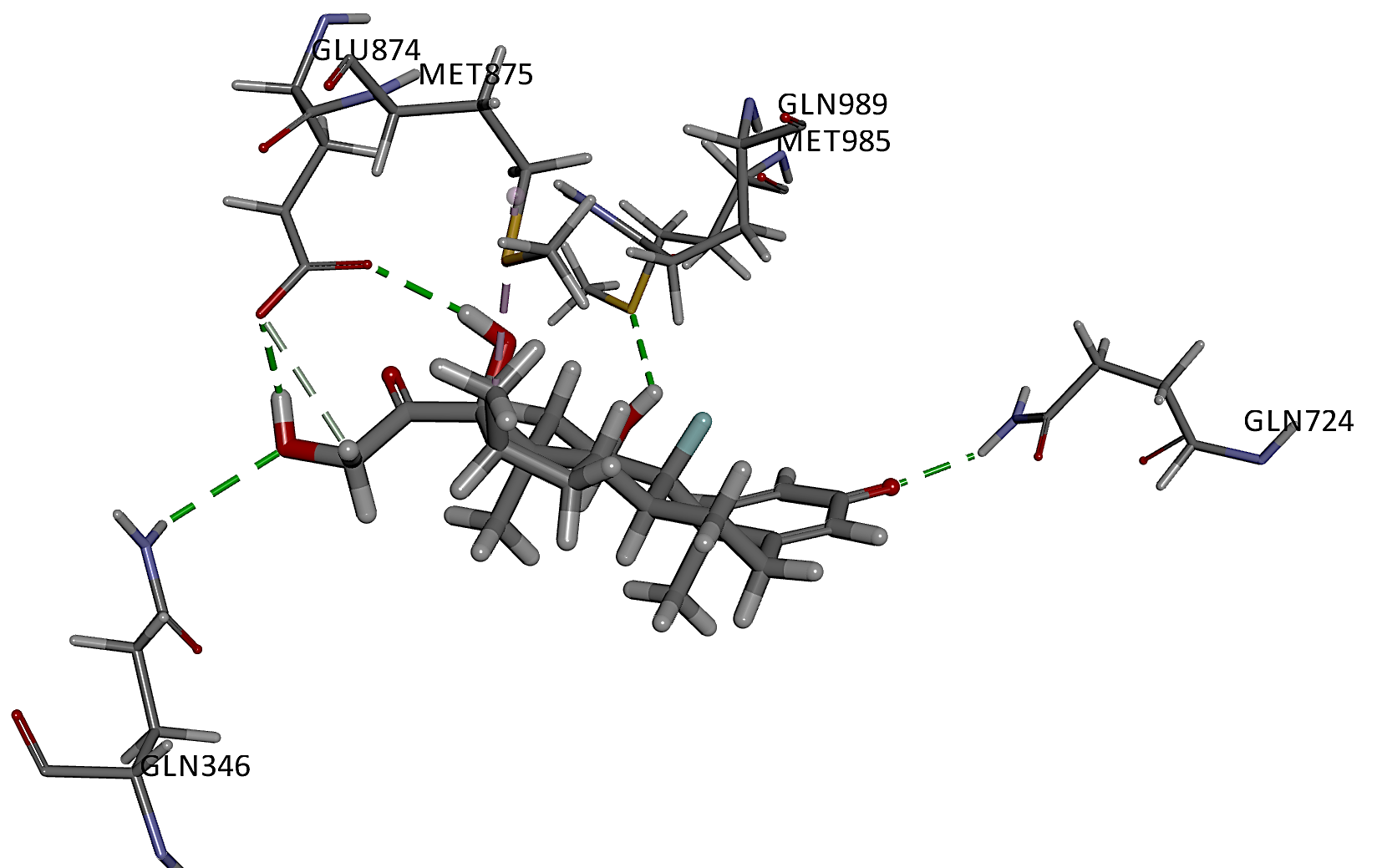

**Figure S17:** Molecular-level interactions of (A) amisulpride, (B) colchicine and (C) dexamethasone with P-gp. Amisulpride in stick-representation and amino acid residues in line-representations. Hydrogen bonds are represented in green dotted lines, and hydrophobic interactions are represented in pink dotted lines.

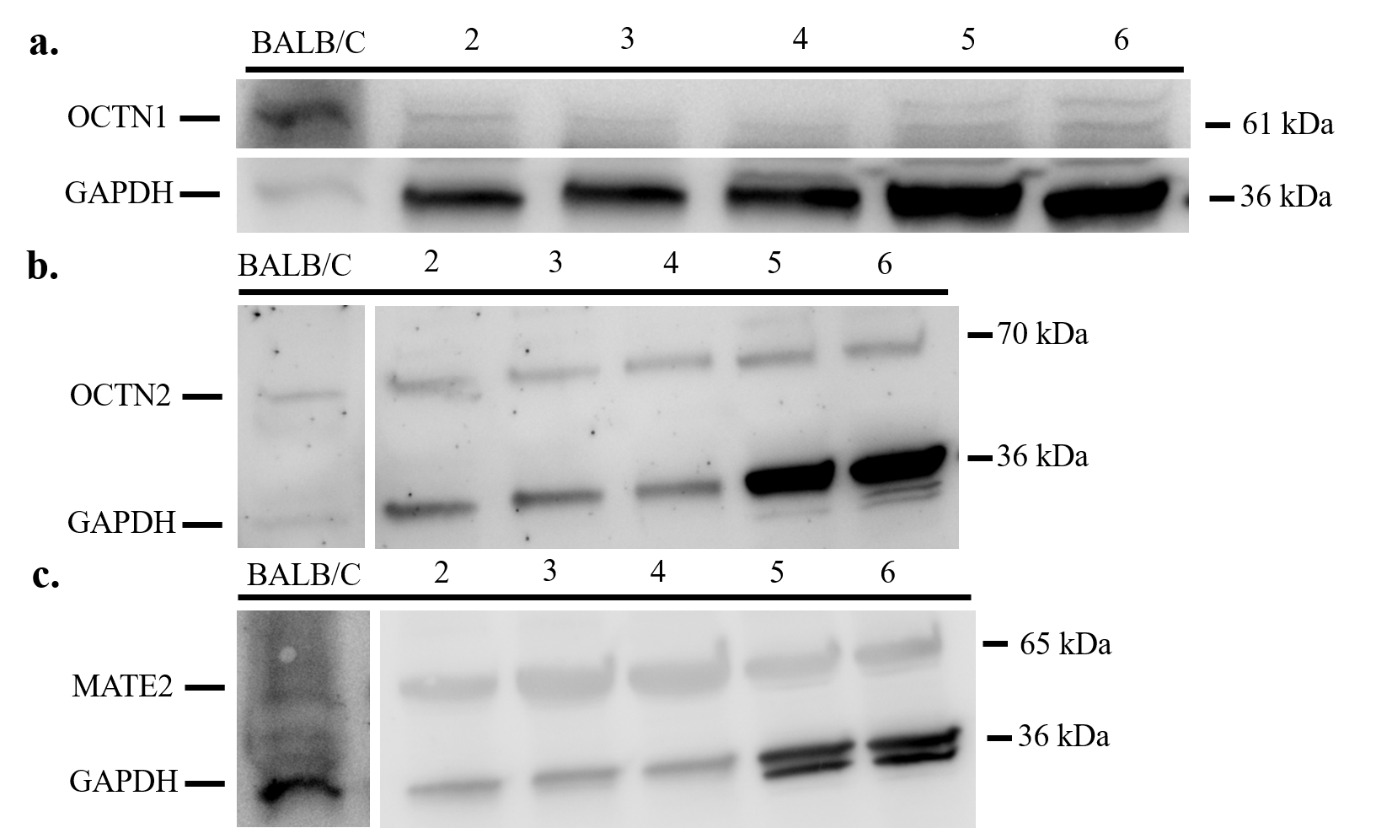

**Figure S18:** OCTN and MATE2 expression in confluent bEnd.3 cells and hCMEC/D3 cells. a. OCTN1 expression was observed in both cell lines at all passages at 61 kDa and in BALB/C (positive control) b. OCTN2 bands were also observed at 70 kD all lanes c. MATE2 expression was observed in all cell lines at 65 kDa. GAPDH expression was used as a loading control at 36 kDa. BALB/C – Brain endothelial cells isolated from mouse (positive interblot control), 2 - bEnd.3 passage 18, 3 – bEnd.3 passage 19, 4 – bEnd.3 passage 23, 5 - hCMEC/D3 passage 28, 6 - hCMEC/D3 passage 33. MATE1 expression was confirmed in bEnd3 cells using a recently available mouse-reactive antibody (data not shown).

**A.**

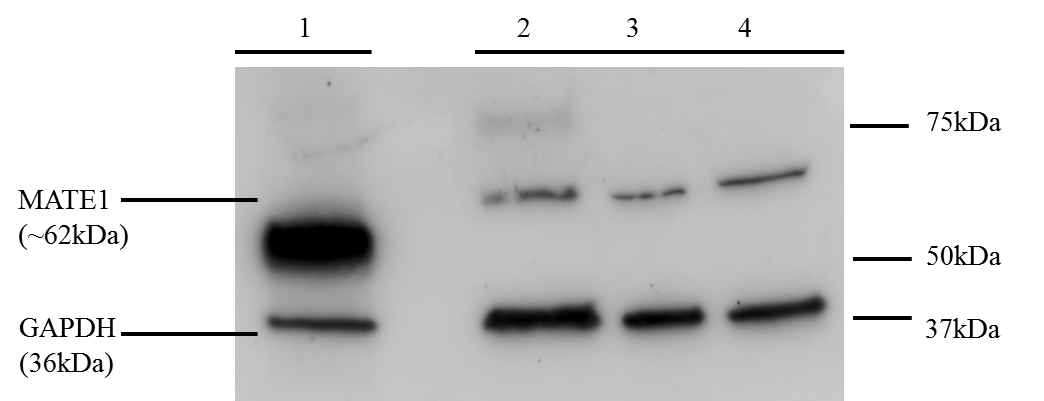

PMAT

GAPDH

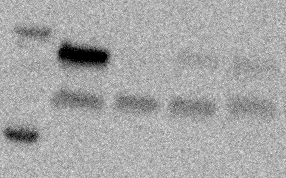

**P28	P30	P31	P32**

**Human**

**capillaries**

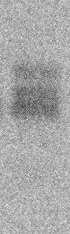

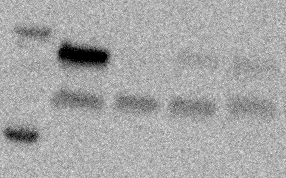

**B.**

58 kDa

**58 kD**

**36 kD**

36 kDa

**36 kD**

**C.**

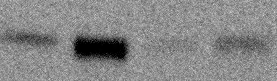

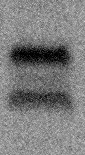

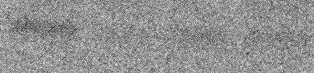

**Human**

**capillaries**

PMAT

GAPDH

**P17	P20	P24	P18**

58 kDa

**58 kD**

**36 kD**

36 kDa

**36 kD**

**Figure S19** **[A]** MATE1 expression in confluent hCMEC/D3 cells at passages 27 (lane 2), 28 (lane 3) and 33 (lane 4) was observed at 62kDa. Lane 1 is MATE1 expression in human cortex endothelium (positive control). GAPDH expression was used as a loading control at 36kDa. **[B]** An example Western blot of PMAT expression in confluent hCMEC/D3 cells at passages 28, 30, 31, 32 were tested for PMAT (58.1 kD) expression. GAPDH (35.8 kDa) was used as a loading control. PMAT was detectable in P28, P31 and P32, but not P30, when 20 μg of protein was utilized. 3 technical replicates were performed. 20 μg of protein was also utilized for the positive control (human capillaries) in the PMAT assay. Anti-PMAT antibody – Cat#:bs-4176R: RRID:AB_11108960. Capillary lysate isolated from the frontal cortex of a healthy aging adult (DPM 14/09; BBN20006; Table S3) was used as a positive control. **[C]** An example Western blot of PMAT (58 kDa) expression in confluent monolayers of b.End3 cells (P17, 20, 24, 18). GAPDH (36 kDa) was used as a loading control. 3 technical replicates were performed. 10μg of protein was utilized for the positive control (human capillaries) in the PMAT assay. Antibodies used: anti-PMAT antibody – 1:650, Cat#bs-4176R, RRID:AB_11108960. ; anti-GAPDH antibody – 1:10,000 (for human capillaries) and 1:2,500 for bEnd.3 cell lysates, Cat#ab9485, RRID:AB_307275; secondary anti-rabbit IgG, HRP-linked antibody – 1:1000, Cat#7074, RRID:[AB_2099233](http://antibodyregistry.org/AB_2099233). Capillary lysate isolated from the frontal cortex of a healthy aging adult (DPM 14/09; BBN20006; Table S3) was used as a positive control.

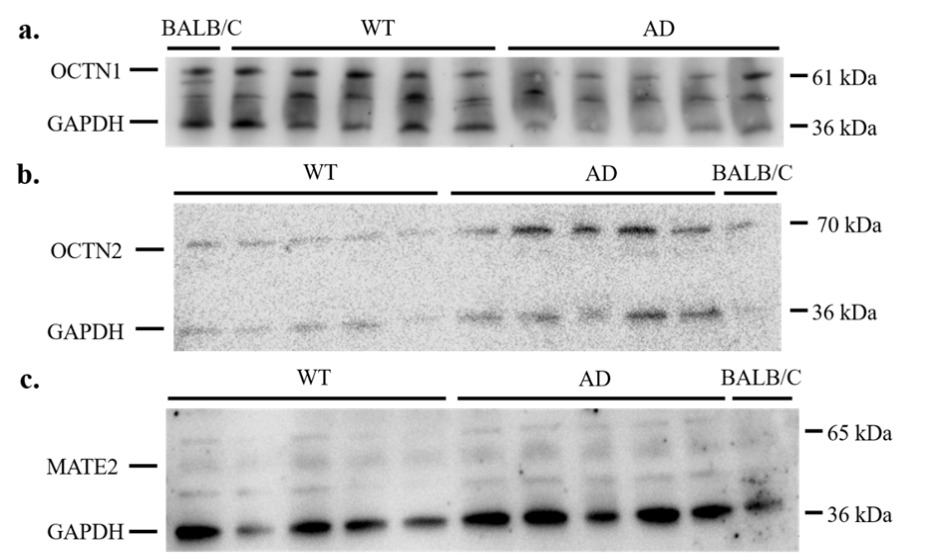

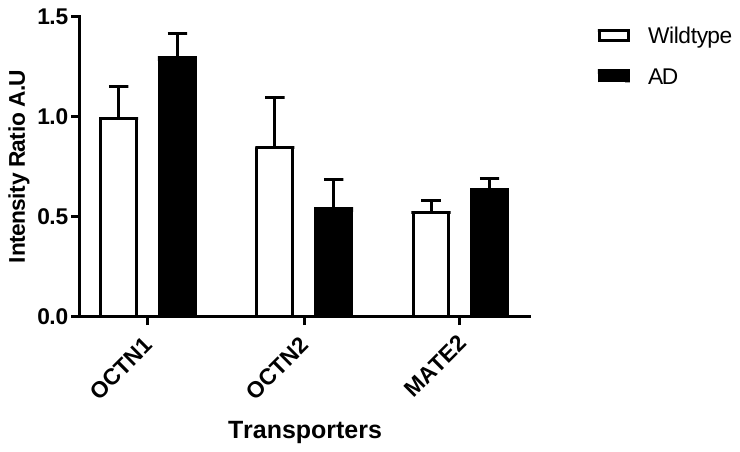

**Figure S20**. Brain capillaries isolated from aged (24 months) C57BL6/129 and 3xTg-AD age-matched mice were tested for a) OCTN1 (61 kDa) and b), OCTN2 (70 kDa) and MATE2 (65 kDa) expression. GAPDH (loading control) detected at 36 kDa. c) Samples plotted for transporter expression. No significant differences were detected by Two-way ANOVA. n=5 for WT and AD mice. Means ± S.E.M.

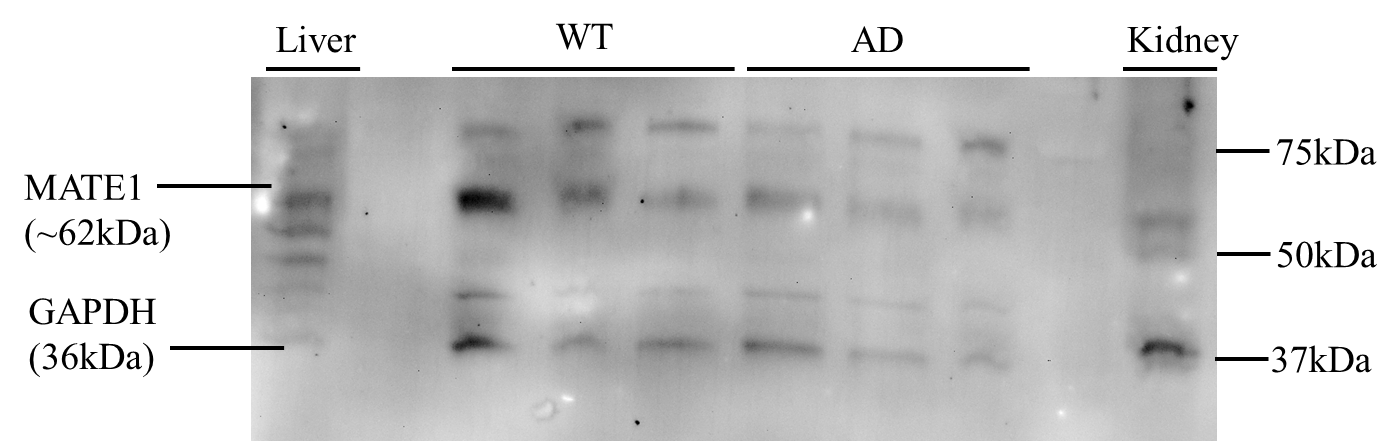

|  | Intensity Ratio A.U. |
| --- | --- |
| Wildtype | 1 |
| AD | 0.750986 |

**Figure S21**. An example Western blot of brain capillaries isolated from old-aged (16 months) C57BL6/129 mice (WT; n=3 different animals) and age-matched 3xTg-AD mice (AD n=4 different animals, this blot shows 3) showing the expression of MATE1 (62kDa) as detected by a recently available mouse-reactive antibody. Mouse (BALB-C) liver and kidney were used as positive controls. GAPDH expression was used as a loading control at 36kDa. Band intensity ratio analysis was conducted using ImageJ software. Samples plotted for MATE1 transporter expression. No significant difference detected by Student’s t-test.

a) Frontal Cortex

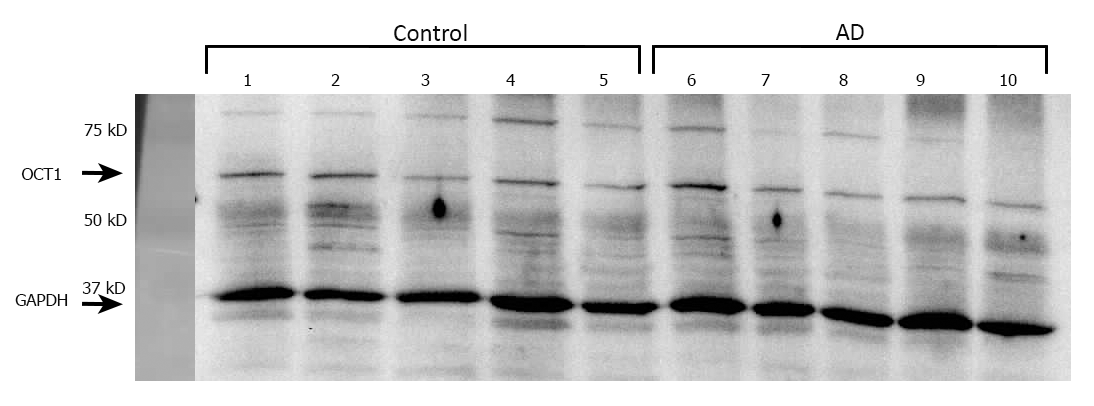

b) Caudate nucleus

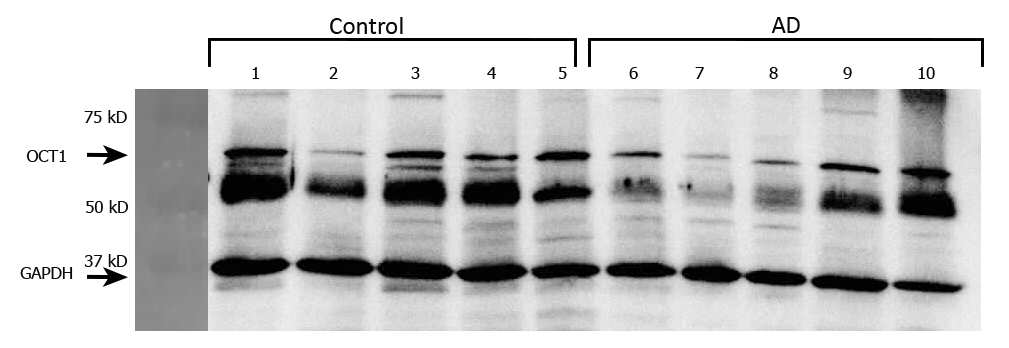

c) Putamen

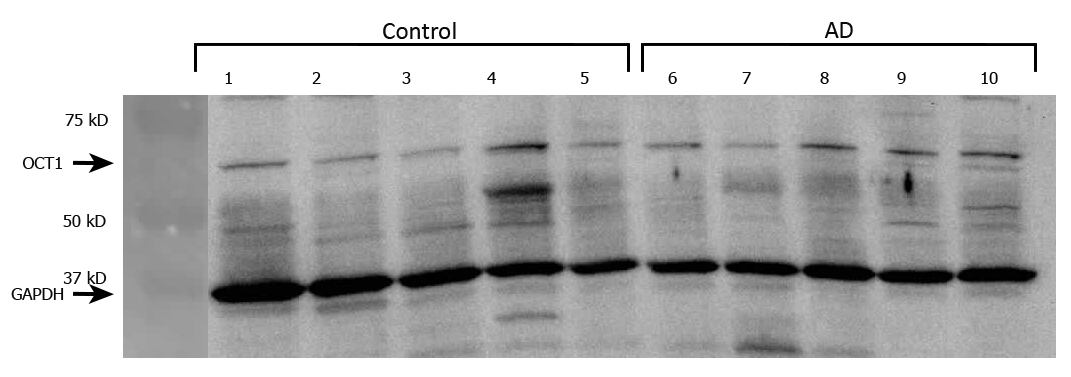

**Figure S22**: Capillaries isolated from the (a) cortex (b) caudate nucleus and (c) putamen of age-matched healthy (lanes 1-5) and AD aﬀected (lanes 6-10) individuals were tested for OCT1 expression using anti-OCT1 antibody (1:400) at 61 kD. GAPDH (loading control) detected at 36 kD.

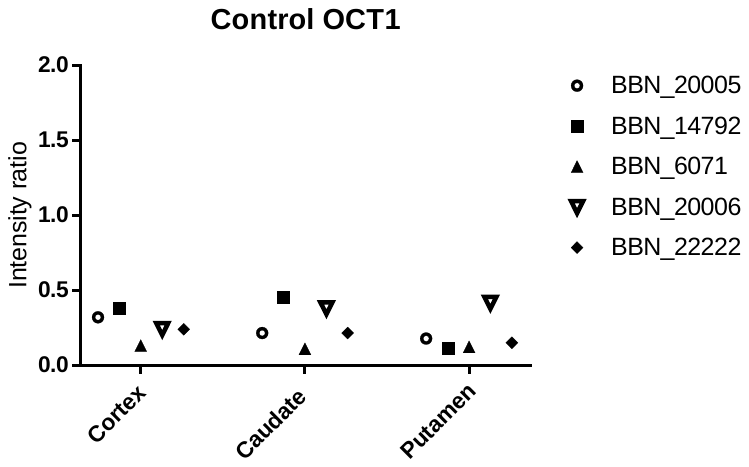

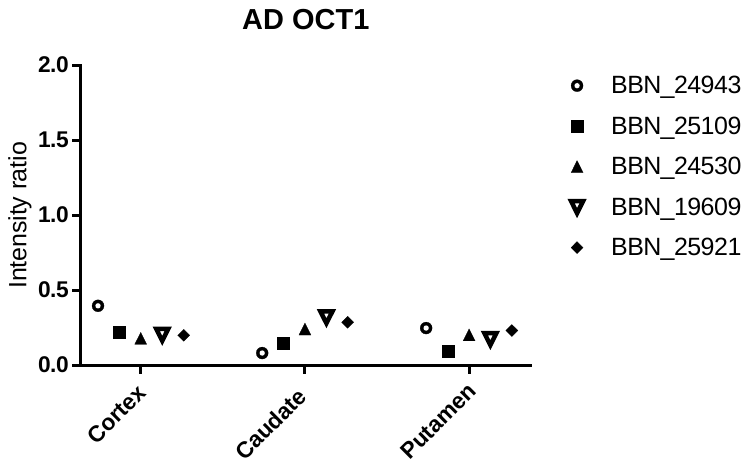

**Figure S23**: Individual values have been plotted for the transporter expression in the capillaries of frontal cortex, caudate nucleus and caudate putamen samples from healthy and AD affected individuals. The numbers in the key indicate the MRC ID designated to each sample. Details of the cases can be found in the Tables S3 and S5.

(a) **Frontal Cortex**

36 kDa

61 kDa

GAPDH

OCTN1

BALB/C

Control

AD

36 kDa

70 kDa

GAPDH

OCTN2

BALB/C

Control

AD

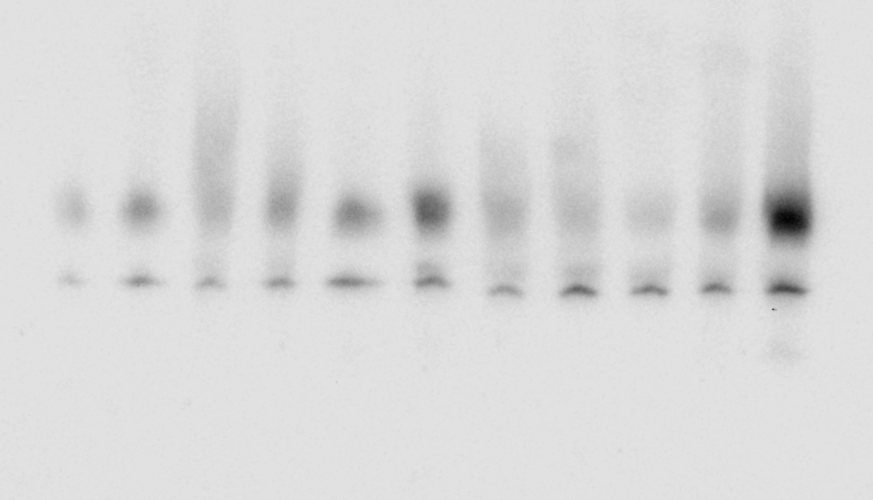

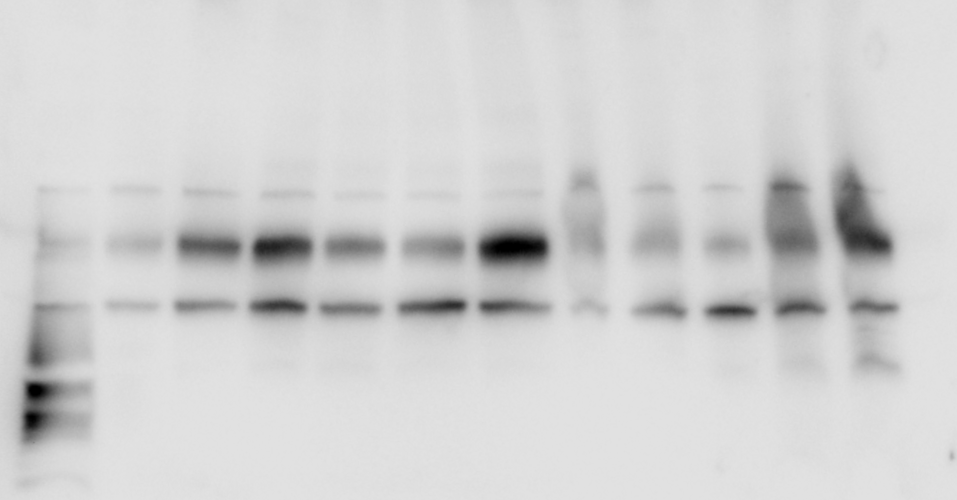

(b) **Caudate Nucleus**

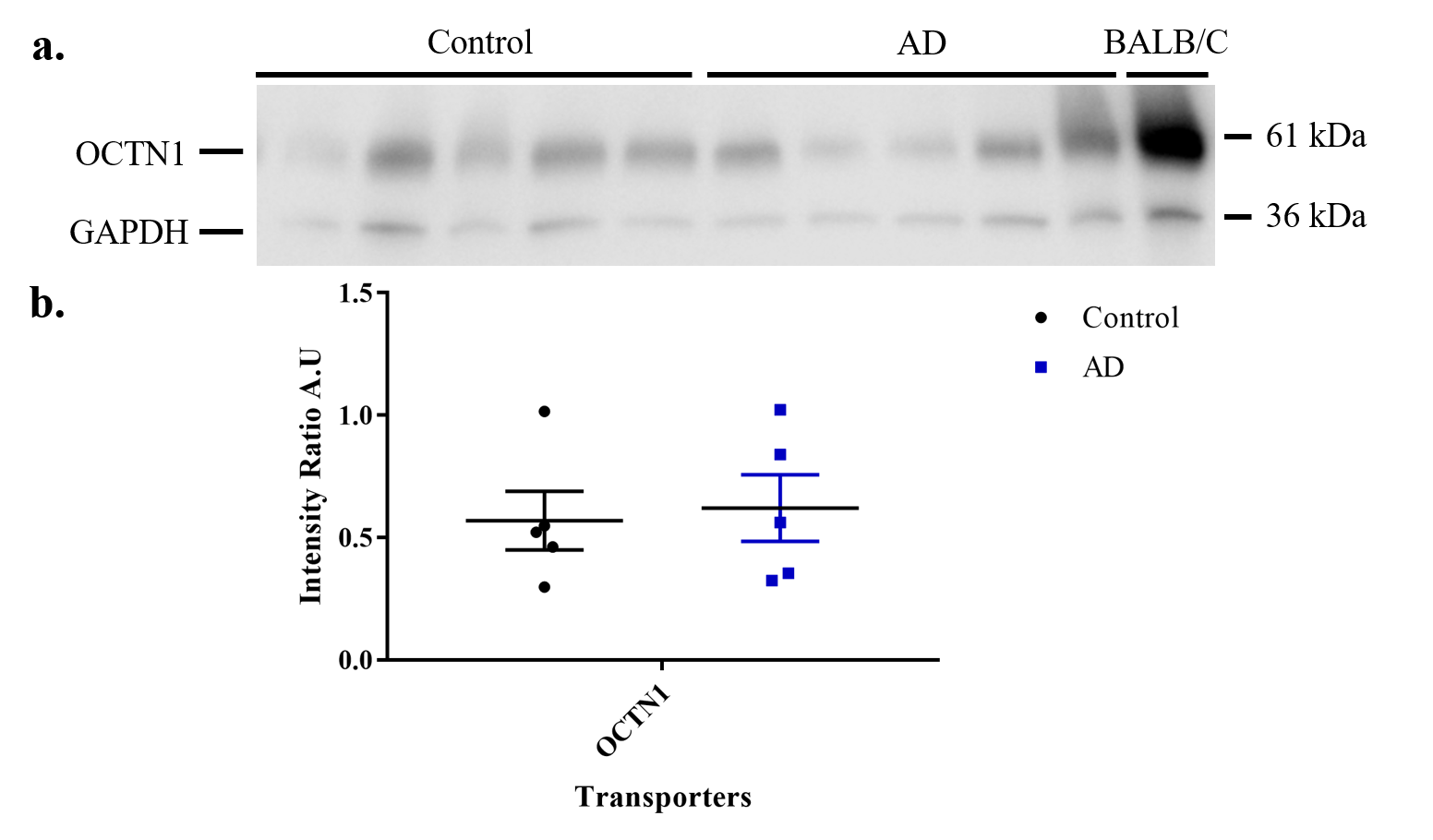

(c) **Putamen**

36 kDa

61 kDa

GAPDH

OCTN1

BALB/C

Control

AD

36 kDa

70 kDa

GAPDH

OCTN2

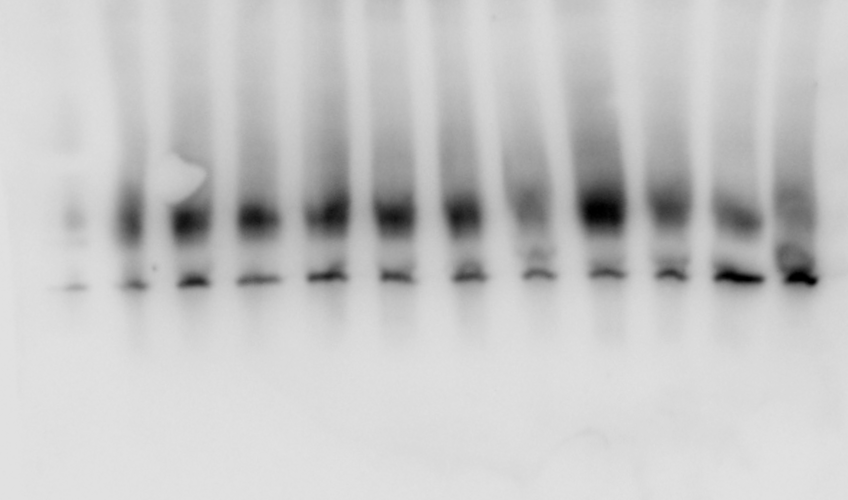

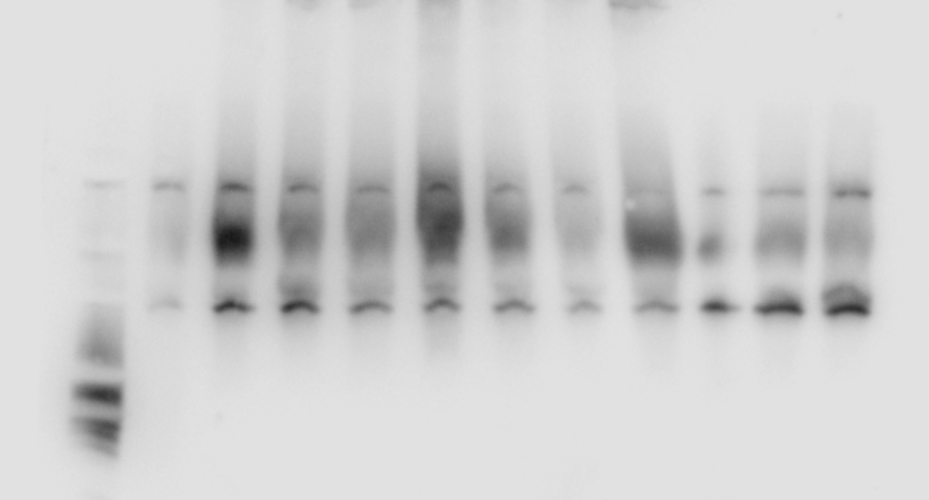

BALB/C

Control

AD

**Figure S24.** Brain capillaries isolated from the frontal cortex, caudate nucleus and caudate putamen of age-matched healthy and AD affected individuals were tested for OCTN1 (61 kDa) and OCTN2 (70 kDa) expression. GAPDH (loading control) detected at 36 kDa.. OCTN2 was not identified in the any of the caudate nucleus samples. This is likely due to technical limitations.

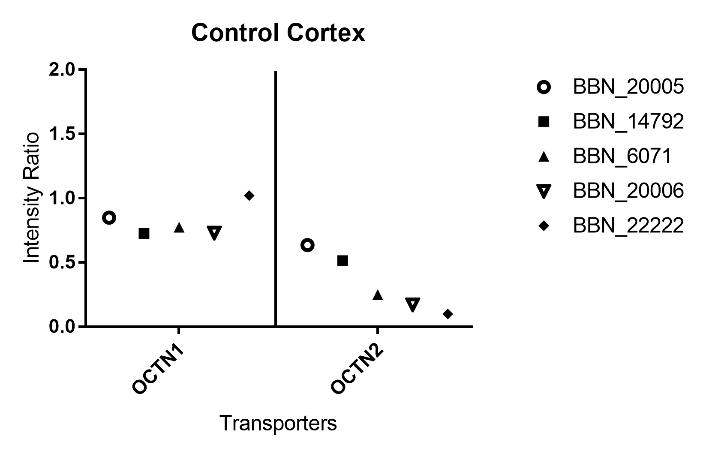

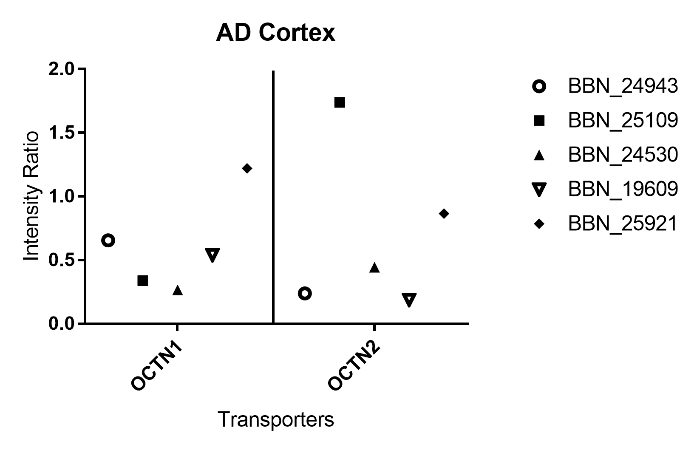

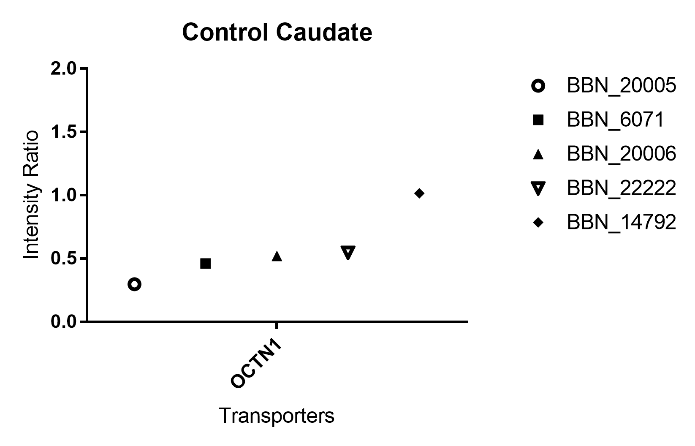

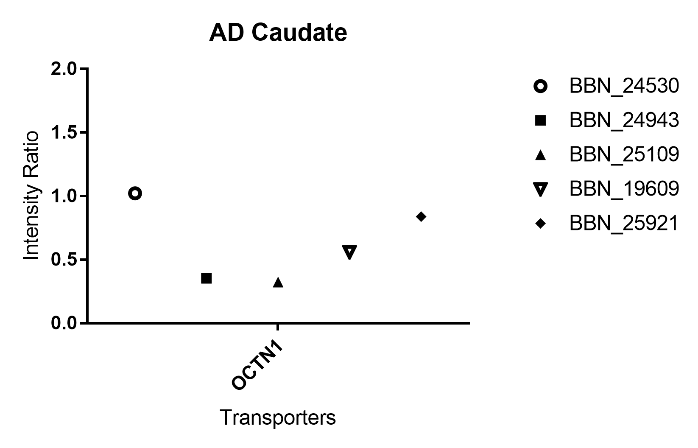

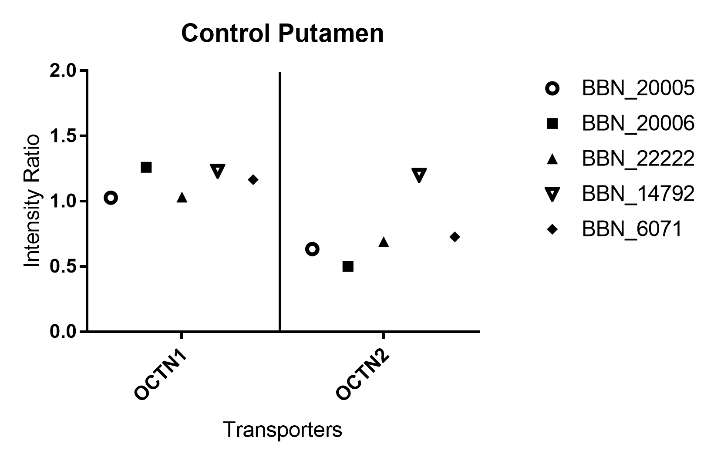

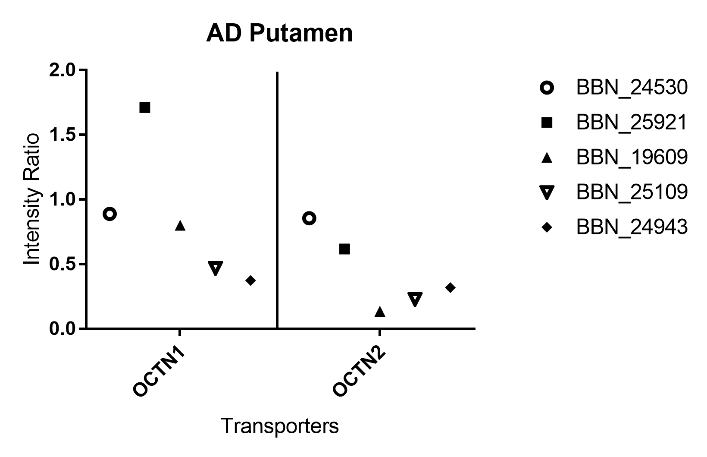

**Figure S25**: Individual values have been plotted for the transporter expression in the capillaries of frontal cortex, caudate nucleus and caudate putamen samples from healthy and AD affected individuals. The numbers in the key indicate the MRC ID designated to each sample. Details of the samples can be found in the Tables S3 and S5.

(a) Frontal cortex

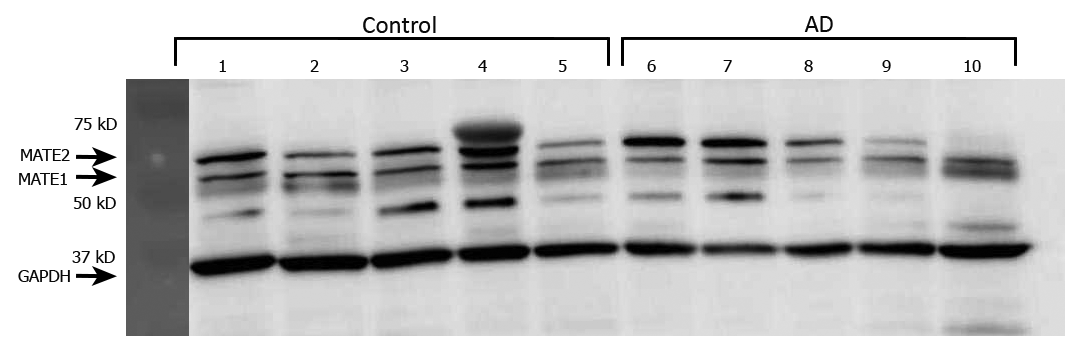

(b) Caudate nucleus

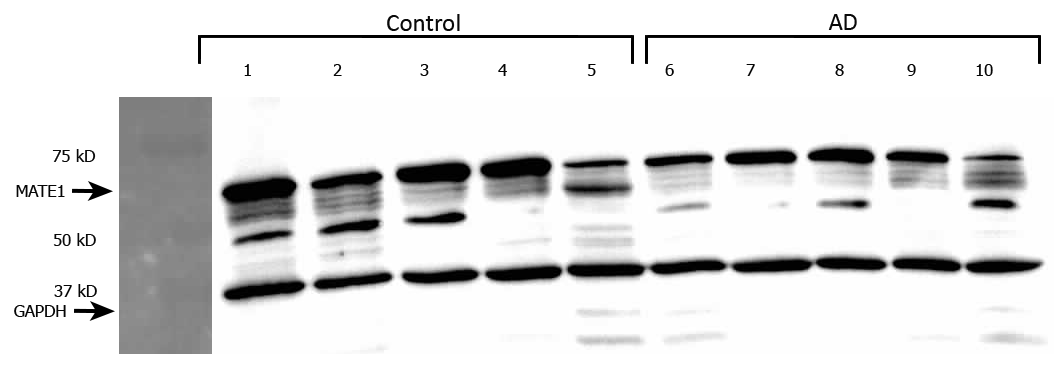

(c) Caudate putamen

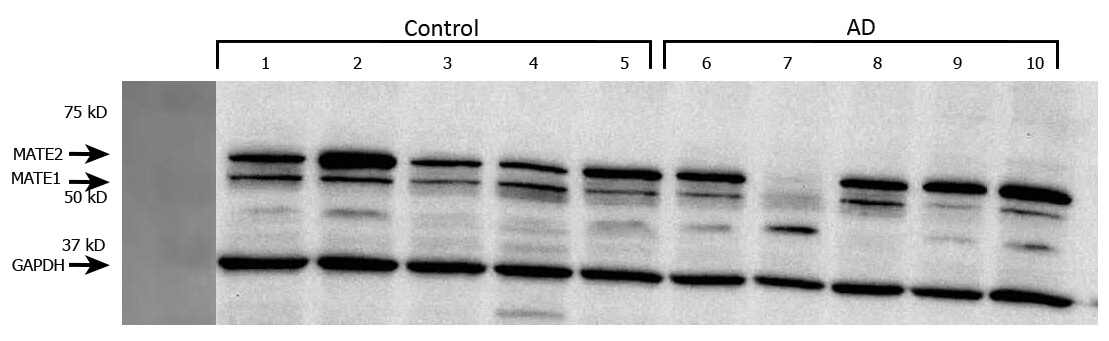

**Figure S26**: Capillaries isolated from the (a) cortex (b) caudate nucleus and (c) caudate putamen of age-matched healthy (lanes 1-5) and AD aﬀected (lanes 6-10) individuals were tested for MATE1 expression using anti-MATE1 antibody (1:500) at 62 kD. GAPDH (loading control) detected at 36 kD.

(a) Frontal cortex

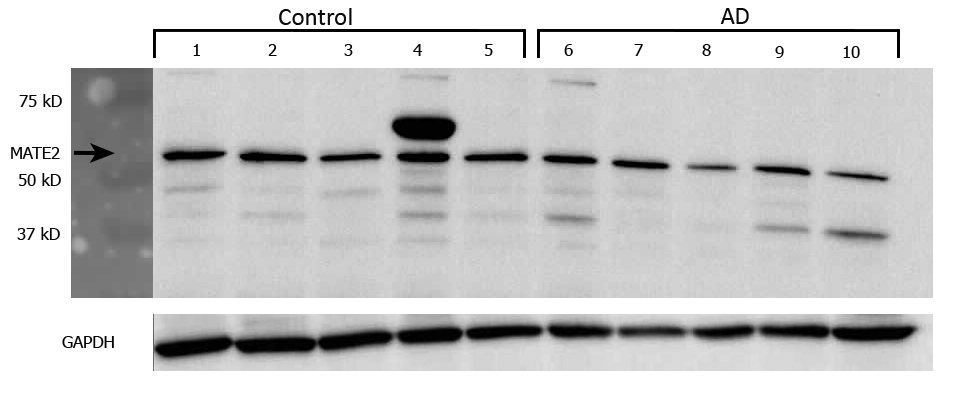

(b) Caudate nucleus

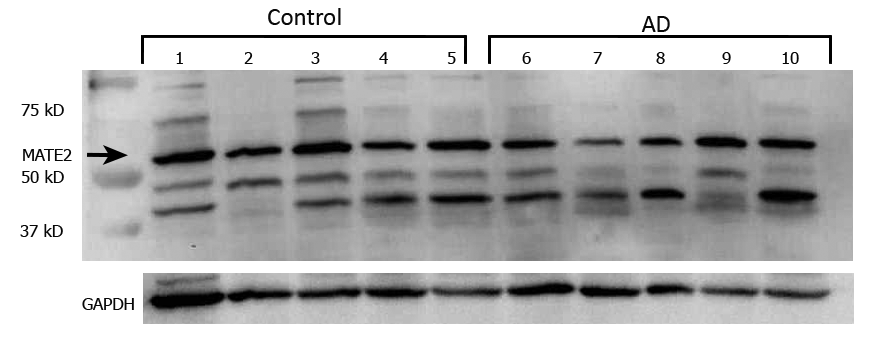

(c) Caudate putamen

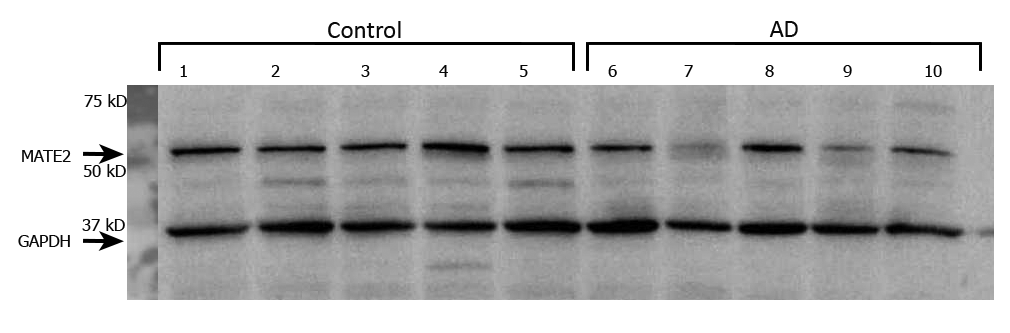

**Figure S27**: Capillaries isolated from the putamen of age-matched healthy (lanes 1-5) and AD aﬀected (lanes 6-10) individuals were tested for MATE2 expression using anti-MATE2 antibody (1:500) at 65 kD. GAPDH (loading control) detected at 36 kD.

(a) Frontal cortex

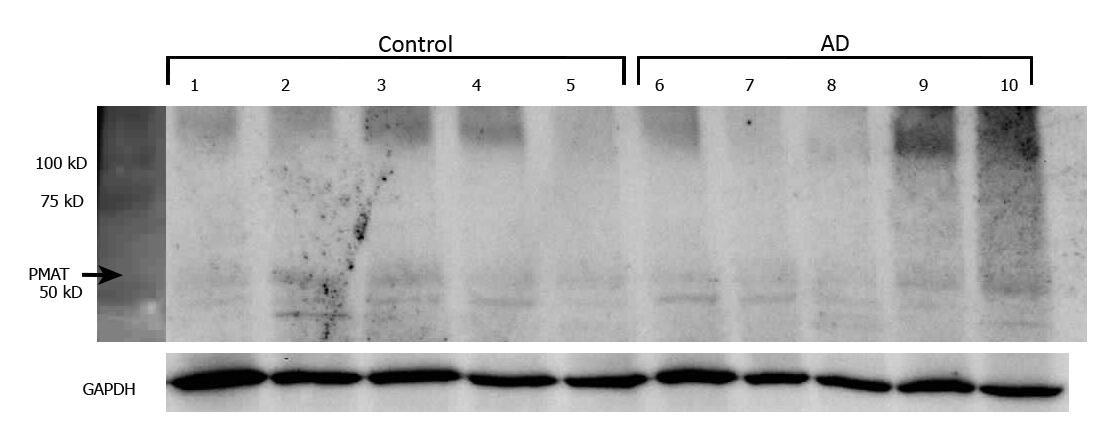

(b) Caudate nucleus

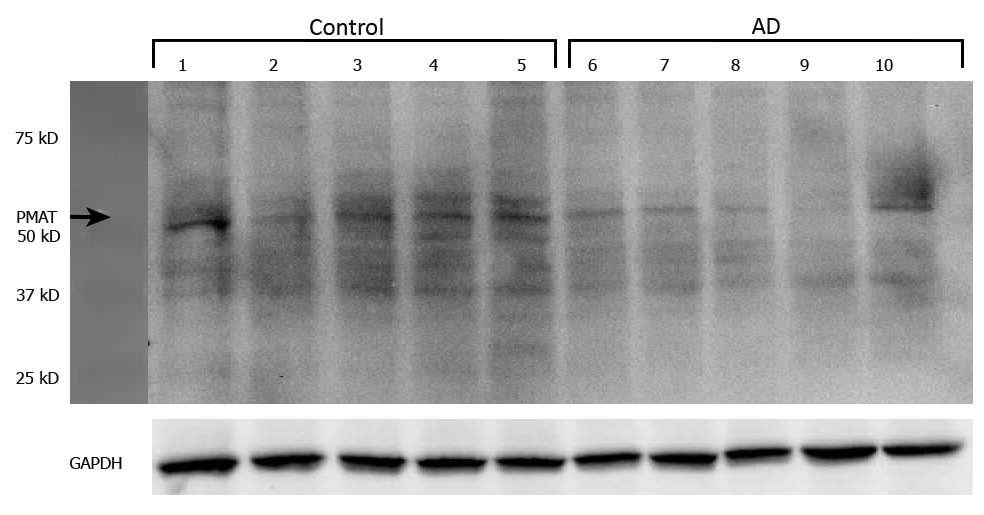

(c) Caudate putamen

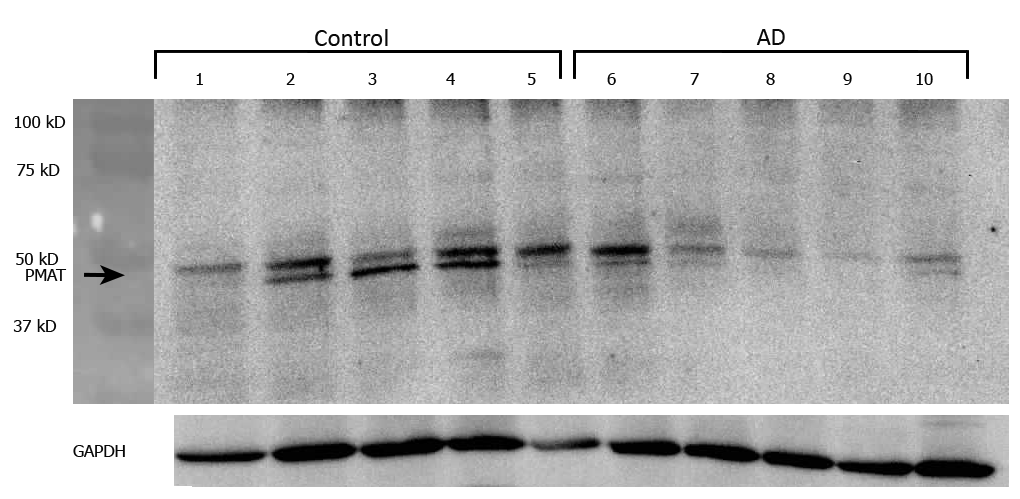

**Figure S28**: Capillaries isolated from the (a) cortex, (b) caudate and (c) putamen of age-matched healthy (lanes 1-5) and AD aﬀected (lanes 6-10) individuals were tested for PMAT expression using anti-PMAT antibody (1:500) at 58 kD. GAPDH (loading control) detected at 36 kD.

**Table S1**: Colchicine and dexamethasone binding interactions with the binding site of the multidrug transporter ABCB1 (P-glycoprotein) newly released pdb code 6FN1.

| **COLCHICINE** | **Amino Acid** | **Type of interaction** | **Distance** |
| --- | --- | --- | --- |
|  | GLN724 | Hydrogen Bond | 2.04567 |
|  | PHE982 | Hydrophobic | 5.13144 |
|  | MET985 | Hydrophobic | 5.3631 |
| **DEXAMETHASONE** | **Amino Acid** | **Type of interaction** | **Distance** |
|  | GLN346 | Hydrogen Bond | 2.8001 |
|  | GLN724 | Hydrogen Bond | 2.10989 |
|  | MET985 | Hydrogen Bond | 2.58018 |
|  | GLU874 | Hydrogen Bond | 2.18713 |
|  | GLU874 | Hydrogen Bond | 2.28905 |
|  | GLU874 | Hydrogen Bond | 3.02681 |
|  | MET875 | Hydrophobic | 4.10009 |

**Table S2:** C57BL6/129 mice (wild-type) were perfused with [^3^H]amisulpride and [^14^C]sucrose for 10 minutes and R_Tissue_ ml.100g^-1^ calculated. All data are expressed as mean ± S.E.M. [^3^H]amisulpride entry into the occipital cortex or caudate was not significantly different to [^14^C[sucrose at any age and ageing did not affect drug distribution or vascular space. (Two-way ANOVA for age (three groups) and molecule (two groups) on brain distribution (dependent variable) followed by Tukey’s multiple comparison data).

| **Occipital Cortex** | **Gender** | **Weight**  **(g)** | **[^3^H]Amisulpride** | **[^14^C]Sucrose** |
| --- | --- | --- | --- | --- |
| Mid-Age (12-13 mths; n=4) | Male | 33.3±1.3 | 2.16±0.36 | 1.94±0.50 |
| Old-Age (16 mths; n=3) | Male | 34.9±0.5 | 1.70±0.02 | 1.56±0.11 |
| Elderly  (24 mths; n=6) | 3 Males and 3 Females | 37.0±1.8 | 1.49±0.22 | 1.39±0.25 |
| **Caudate** |  |  |  |  |
| Mid -Age (12-13 mths; n=4) | Male | 33.3±1.3 | 2.23±0.90 | 1.98±0.29 |
| Old Age (16 mths; n=3) | Male | 34.9±0.5 | 2.35±0.22 | 2.17±0.94 |
| Elderly  (24 mths; n=3) | 2 Males; 1 Female | 35.0±1.0 | 1.63±0.46 | 1.67±0.49 |

**Table S3**: Human brain tissue samples were supplied by The Manchester Brain Bank, which is part of the Brains for Dementia Research programme, jointly funded by Alzheimer’s Research UK and Alzheimer’s Society. The cases were divided into two groups based on Braak staging. The mean age of cases with a Braak stage 0-II was 86.8±1.5years (healthy controls), which was not significantly different (unpaired t-test) to the cases with a Braak-stage of V-VI, which was 79.4±3.7 years (AD affected individuals). The mean post-mortem delay was 91.7±8.9 (healthy controls) versus 90.8±12.3 hours (AD affected individuals). Values are not significantly different to each other (unpaired t-test). The mean post-mortem pH was 6.00±0.15 (healthy controls) and 6.17±0.10 (AD affected individuals). Values are not significantly different to each other (unpaired t-test). Three brain regions (frontal cortex, caudate nucleus, and putamen) were obtained from each of the cases.

| **Manchester ID (DPM)** | **MRC ID Number** | **BDR ID** | **Braak stage** | **Post-mortem pH** | **Post-mortem delay (hours)** | **Gender** | **Age at death** |
| --- | --- | --- | --- | --- | --- | --- | --- |
| 12/20 | BBN_6071 | 426 | II | 5.63 | 92 | Male | 84 |
| 13/25 | BBN_14792 | 612 | II | 6.47 | 78 | Female | 91 |
| 14/08 | BBN_20005 | 226 | 0-I | 6.27 | 98 | Male | 85 |
| 14/09 | BBN_20006 | 255 | I | 5.84 | 69.5 | Male | 84 |
| 14/27 | BBN_22222 | 480 | I-II | 5.81 | 121 | Female | 90 |
| 14/03 | BBN_19609 | 608 | VI | 6.05 | 62 | Male | 75 |
| 15/07 | BBN_24530 | 403 | VI | 6.02 | 131 | Female | 91 |
| 15/20 | BBN_24943 | 410 | VI | 6.21 | 96 | Male | 71 |
| 15/21 | BBN_25109 | 484 | VI | 6.54 | 97 | Female | 79 |
| 15/29 | BBN_25921 | 830 | V-VI | 6.03 | 68 | Male | 81 |

**Table S4**: *Medication history of the cases (those with dementia-Braak stage V-VI) and healthy controls (Braak stage 0-II)*. Medication details came from a variety of sources including a combination of face to face and telephone assessments by BDR team and from GP information after death. The BDR information relates to the medications the patient was receiving on the day of the assessment. BDR assessments were carried out on an annual basis for Cases (those with dementia) and every two years for healthy controls. For Cases their nominated representative (normally a spouse or offspring) would always be present and once they were too ill to be assessed (due to the progression of the dementia), just the nominated representative was interviewed. The nominated representative guided BDR as to the suitability/appropriateness of interviewing the Case. Tissue samples were supplied by The Manchester Brain Bank, which is part of the Brains for Dementia Research programme, jointly funded by Alzheimer’s Research UK and Alzheimer’s Society. *Identified as OCT1 inhibitors (Ahlin et al., 2008; Bourdet et al., 2005).

| ***Manchester ID (DPM)*** | ***MRC***  ***ID*** | ***BDR ID*** | ***Braak Stage*** | ***Sedatives, antidepressants and antipsychotics*** | ***Other medications*** |
| --- | --- | --- | --- | --- | --- |
| 12/20 | BBN_6071 | 426 | II | * Haloperidol- ***first generation antipsychotic***  Oxynorm, - ***codeine/opioid*** | Aspirin- ***non-steriodal anti-inflammatory (NSAID)***  Gaviscon -***antacid***  Humulin injection- ***insulin***  Codanthramer – ***laxative***  Bisoprolol – ***beta blocker***  *Ranitidine – ***H2 blocker antacid***  *Metoclopramide- ***antiemetic***  Hyoscine – ***muscarinic antagonist***  Paracetamol |
| 13/25 | BBN_14792 | 612 | II |  | Amlodipine- ***calcium channel blocker, anti-hypertensive***  Warfarin- ***coumarin*** |
| 14/08 | BBN_20005 | 226 | 0-I |  | Digoxin – ***anti-arrythmic,***  Bumetamide - ***diuretic*** |
| 14/09 | BBN_20006 | 255 | I | Tramadol – ***opioid***  Pregabalin, gabapentin – same drug, ***anti-epileptic with sedating and analgesic action***  *Citalopram **– *antidepressant (SSRI)*** | Aspirin- ***NSAID***  Furosemide- ***diuretic***  Lisinipril – ***ACE inhibitor***  Lansoprazole – ***antacid/blocks Na^+^ pump***  Calcichew – ***calcium supplement***  Madapar – ***antiparkinson’s***  Atorvastatin- ***lipid lowering agent***  Humilin M3 injection – ***insulin***  Novorapid, insulatard – ***insulin long acting*** |
| 14/27 | BBN_22222 | 480 | I-II |  | Bendroflumethiazide- ***diuretic***  Levothyroxine- ***thyroxine supplement***  Calcichew- ***calcium supplement***  Alendronic acid – ***bisphosphonate (bone health)***  Clopidogril- ***antithrombotic***  Simvastatin – ***lipid lowering*** |
| 14/03 | BBN_19609 | 608 | VI |  | Chlormethiazide - ***diuretic***  Furosemide - ***diuretic***  Paracetamol- ***anti pyretic*** |
| 15/07 | BBN_24530 | 403 | VI |  | Aspirin - ***NSAID***  Lactulose- ***laxative***  Lercanipine- ***anti-hypertensive***  Lisinipril- ***ACE inhibitor (antihypertensive)***  Methocarbamol- ***muscle relaxant, sedative*** |
| 15/20 | BBN_24943 | 410 | VI | Mirtazapine**- antidepressant (SNRI)**  Risperidone- **second generation antipsychotic**  Sodium valproate- **anti-epileptic**, sometimes used to treat agitation, | Lactulose, senna – ***laxatives*** |
| 15/21 | BBN_25109 | 484 | VI | Mirtazapine**- antidepressant** | *Loratadine- ***antacid, H2 antagonist***  Adcal- ***calcium supplement***  Amlodipine- ***anti-hypertensive***  Aspirin- ***NSAID***  Canderstartin- ***angiotensin antagonist (anti-hypertensive)***  Clopidrogel - ***antithrombotic***  Furosemide **– *diuretic***  Levothyroxine- ***thyroxine***  Omeprazole- ***antacid***  Raloxifene – ***bone health***  Simvastatin- ***lipid lowering***  Galantamine- ***Acetyl cholinesterase inhibitor (ChEI)*** |
| 15/29 | BBN_25921 | 830 | V-VI | Fluoxetine- **antidepressant (SSRI)**  Zopiclone- **sedating, anxiolytic** | Tansulosin – ***alpha 1 adrenergic blocker***  Omeprazole- ***antacid*** |

**Table S5**: Total protein expression of brain samples obtained from the cases (those with dementia-Braak stage V-VI) and healthy controls (Braak stage 0-II) described in Table S3. All data is expressed as mean±SEM. Three brain regions (frontal cortex, caudate nucleus and putamen) were obtained from each of the cases. Two-Way ANOVA indicated there was a significant difference between the controls and dementia cases (P<0.0001) and also a difference between the brain regions (P<0.001). Post-hoc analysis was by Tukey’s multiple comparison and revealed that the healthy controls had significantly higher protein expression in the caudate nucleus and putamen compared to similar samples from those cases with dementia. No difference between the groups was observed for the frontal cortex.

|  | **Healthy controls (Braak stage 0-II)**  **n=5**  **μg of protein/100μl of RIPA buffer** | **Dementia**  **(Braak stage V-VI)**  **n=5**  **μg of protein/100μl of RIPA buffer** | **Post-hoc analysis**  **multiple comparisons** |
| --- | --- | --- | --- |
| **Frontal cortex** | 88.8±7.3 | 77.3±5.9 | P>0.05 |
| **Caudate nucleus** | 107.8±6.9 | 67.5±4.5 | P<0.001 |
| **Putamen** | 128.8±3.5 | 86.9±7.3 | P<0.0001 |

**Table S6:** Primary and Secondary antibodies used for protein expression studies. The predicted antibodies with their predicted molecular weight (MW) WB were all made up in PBS-T with 5% BSA. Actual band sizes may differ due to post translational modifications or cleavages. (Validation data is available from the Abcam website (<https://www.abcam.com/nav/primary-antibodies>), Cell signalling technology website (<https://www.cellsignal.co.uk>) and Alonome labs website (<https://www.alomone.com>). Accessed 19.10.18. Validation data is available from the BIOSS USA antibodies website (<http://www.biossusaantibodies.com>). Accessed 4.12.18.

| **PROTEIN** | **Primary Antibody** | **Secondary Antibody WB** |
| --- | --- | --- |
| **OCT-1 (SLC22A1)** | Rabbit polyclonal anti-human and mouse (Abcam, Cat#ab55916; RRID:AB_882579)  WB Dilution- 1:250 | Goat anti-rabbit HRP (Abcam, cat#ab6721: RRID:AB_955447)  Dilution - 1:1000 |
| **OCT-2 (SLC22A2)** | Rabbit monoclonal to human and mouse (Abcam, Cat#ab170871: RRID:AB_2751021)  WB Dilution – 1:2000 | Goat anti-rabbit HRP (Abcam, cat#ab6721: RRID:AB_955447)  Dilution - 1:2000 |
| **OCT -3 (SLC22A3)** | Rabbit polyclonal to human and mouse (Abcam, Cat#ab183071; RRID:AB_2751016),  WB Dilution – 1:600 | Goat anti-rabbit HRP (Abcam, cat#ab6721: RRID:AB_955447)  Dilution - 1:2000 |
| **OCTN1**  **(SLC22A4)** | Rabbit polyclonal to human and mouse (Abcam, Cat#ab200641; RRID:AB_2751017),  WB Dilution – 1:1000 | Goat anti-rabbit (IgG)-HRP (Cell Signalling. Cat#7074S:[AB_2099233](http://antibodyregistry.org/AB_2099233)) Dilution -1:1000 |
| **OCTN2**  **(SLC22A5)** | Rabbit polyclonal to human and mouse (Abcam, Cat#ab180757; RRID:AB_2751018),  WB Dilution – 1:1000 | Goat anti-rabbit (IgG)-HRP (Cell Signalling. Cat#7074S: [AB_2099233](http://antibodyregistry.org/AB_2099233)) Dilution -1:1000 |
| **MATE1**  **(SLC47A1)** | Rabbit polyclonal to human  from Abcam ( Cat#ab104016: RRID [AB_10711136](http://antibodyregistry.org/AB_10711136)), WB Dilution - 1:500 | Goat anti-rabbit HRP (Abcam, Cat#ab6721: RRID:AB_955447) Dilution - 1:2000 |
| **MATE1**  **(SLC47A1)** | Rabbit polyclonal to mouse from Alomone labs ( Cat#ANT-131: RRID:AB_2751020),  WB Dilution – 1:800 | Goat anti-rabbit HRP (Abcam, Cat#ab6721: RRID:AB_955447) Dilution - 1:2000 |
| **MATE2**  **(SLC47A2)** | Rabbit polyclonal to human and mouse (Abcam, Cat#ab174344: RRID:AB_2751019),  WB Dilution – 1:500 | Goat anti-rabbit HRP (Abcam, Cat#ab6721: RRID:AB_955447) Dilution - 1:2000 |
| **PMAT**  **(SLC29A4)** | Mouse monoclonal to human and rat (Abcam, Cat#ab56554: RRID:AB_2190909),  WB Dilution - 1:500 | Rabbit anti-mouse HRP (Abcam, Cat#ab6728: RRID:AB_955440) Dilution 1:2000 |
| **PMAT**  **(SLC29A4)** | Rabbit polyclonal to human, mouse and rat (Bioss Antibodies; Cat#:bs-4176R: RRID:AB_11108960) ,  WB Dilution  -1:800 for hCMEC/D3  -1:650 for b.End3 | Goat anti-rabbit (IgG)-HRP (Cell Signalling. Cat#7074S: RRID:[AB_2099233](http://antibodyregistry.org/AB_2099233)) Dilution -1:1000 |
| **GAPDH** | Rabbit polyclonal to GAPDH (Abcam, Cat#ab9485: RRID:AB_307275)  WB Dilution 1: 2500 or 1:10,000 |  |

| **Table S7:** The transporter inhibitors used in this study along with [^3^H]amisulpride. All inhibitors were used in the presence of 0.05% DMSO and used at the published concentration ranges where they affect transporter activity. | | | | |
| --- | --- | --- | --- | --- |
| **Target Transporter** | **Transporter**  **Inhibitor/substrate** | **Concentration** | **Catalogue number (where available) and Source** | **References** |
| - | Amisulpride | 0.1-20 µM | 71675-85-9  Cayman Chemicals, UK |  |
| P-gp /MDR1  (ABCB1) | Dexamethasone | 200 µM | Sigma, UK | (Watson *et al.*, 2012) |
| BCRP  (ABCG2) | ko143 | 1 µM | 3241  Tocris Bioscience, UK | (Matsson *et al.*, 2009) |
| BCRP  (ABCG2) | Pheophorbide A | 1 µM | sc-264070  Santa Cruz Biotechnology, Germany | (Watson *et al.*, 2012) |
| MRPs | MK571 | 10 µM | CAY10029  Cambridge Bioscience, UK | (Poller *et al.*, 2008) |
| P-gp /MDR1  (ABCB1) and OCT including OCT1 (SLC22A1) | Haloperidol | 40 µM | H1512  Sigma, UK | (Iwaki *et al.*, 2006)(Kang *et al.*, 2006; Ahlin *et al.*, 2008)(Sekhar *et al.*, 2017) |
| OCT1 (SLC22A1) and  OCT3 (SLC22A3) | Prazosin | 100 µM | P7791  Sigma, UK | (Sekhar *et al.*, 2017)(Dickens *et al.*, 2012) |
| OCT1 (SLC22A1) and  OCT2 (SLC22A2)  MATE1 (SLC47A1) and MATE 2 (SLC47A2)  PMAT (SLC29A4) | Amantadine | 500 µM | 1339  Tocris, UK | (Sekhar *et al.*, 2017), (Nies *et al.*, 2011), (Tsuda *et al.*, 2009)(Itagaki *et al.*, 2012) |
| OCT3 (SLC22A3) | Corticosterone | 50 µM | 27840  Sigma, UK | (Dickens *et al.*, 2012) |
| OCTN1 (SLC22A4) | Ergothioneine | 20 µM | sc-200814  Santa Cruz Biotechnology, Inc, USA. | (Gründemann *et al.*, 2005) |
| OCTN2 ( SLC22A5) | L-carnitine | 5 µM | sc-205727  Santa Cruz Biotechnology, Inc, USA. | (Tamai *et al.*, 1998) |
| PMAT (SLC29A4) | Lopinavir | 2 µM | SC-297831  Santa Cruz Biotechnology, Inc, USA. | (Duan *et al.*, 2015) |
| MATE1 (SLC47A1) | Famotidine | 1 µM | SC-205691  Santa Cruz Biotechnology, Inc, USA. | (Wittwer *et al.*, 2013) |
| MATE2 (SLC47A2) | Nifekalant hydrochloride | 3 µM | SC-204819  Santa Cruz Biotechnology, Inc, USA. | (Wittwer *et al.*, 2013) |
